# Oncogenic PI3Kα variants reveal graded conformational spectrum with mutation-specific cryptic pockets

**DOI:** 10.1101/2025.06.26.661751

**Authors:** Hyunbum Jang, Bengi Ruken Yavuz, Mingzhen Zhang, Yonglan Liu, Ruth Nussinov

## Abstract

Cancer-prone alleles exhibit single hotspot mutations. However, the combination of a cancer hotspot and a weak or moderate mutation (the ‘one-two punch’ hypothesis) produces same-allele double variants with a significantly different and potentially graded clinical phenotypic spectrum. Oncogenic PI3Kα variants, which are also associated with benign tumors and neurodevelopmental disorders, offer statistical support for this model. Using atomistic molecular dynamics (MD) simulations, we revealed that PI3Kα variants with single and double mutations exhibit expanded conformational profiles. Double mutations significantly shift the conformational ensembles toward the active form—a more pronounced effect than a single mutation. These double mutants facilitate nSH2 release, iSH2 shift, and A-loop protrusion in solution, promoting PIP_2_ substrate recruitment at the membrane. We discovered that observable, potentially drug-targetable cryptic pockets are PI3Kα mutation-specific. *Ab initio* discovery of these pockets through simulations is essential for AI-aided virtual drug screening. A key challenge is that a single drug is often ineffective against PI3Kα variants due to their diverse conformational spectra. To address this, we propose a conformational selection strategy involving a combination of allosteric drugs for variants with graded conformational spectra, particularly those with strong double mutations; *we identified such potentially targetable cryptic pockets in double mutants conformers*.

## Introduction

Mutations in oncogenic proteins commonly influence their tendency to adopt active conformations. Multiple observations confirm that cancer emergence requires more than a single mutation^1^. Often one of them is a hotspot. A strong hotspot mutation shifts the equilibrium toward the active state; this shift may be observable^2^. A weak pathogenic mutation minimally shifts the equilibrium toward the active state. An observable phenotypic presentation is likely to be clinically more severe. While different mutations (and their combinations) can populate the same conformations, the extent to which each mutational variant occupies each conformation is unclear. This information is critical because proteins function properly when they are predominantly in their active state. Here, we aim to delineate conformational variants of phosphoinositide 3-kinase α (PI3Kα) by comparing an inactive wild-type form to its mutants, which exhibit a graded conformational spectrum toward an active form. We propose that weak mutations have a mild effect and tend to resemble the wild-type conformational distribution. In contrast, stronger oncogenic mutations stabilize the active state and/or destabilize the inactive state, driving the ensemble toward the active state.

Elevated PI3Kα signaling is a hallmark of cancer^3^. Under physiological conditions, PI3Kα signaling is initiated downstream by stimulated RTKs and small GTPases of the Ras superfamily^4^. However, mutations can work by either bypassing the action of RTK in relieving the PI3Kα autoinhibition or bypassing the contribution of Ras to the enhancement of membrane attachment^5–7^. In case of oncogenic charge reversal by mutations E542K and E545K, strong electrostatic repulsion in the helical domain disengages it from nSH2, shifting the equilibrium toward the nSH2-relieved state of p110α. These conformational changes result in exposure of the kinase domain for membrane interaction and activation^8–10^. Ras binding to the Ras binding domain (RBD) of p110α enhances the membrane interaction of wild-type PI3Kα by promoting its localization and optimal orientation. The oncogenic H1047R hotspot mutation replaces Ras actions by increasing the positive charge on the membrane-interacting surface of the kinase domain^11^. This stabilizes the open, well-positioned conformation of the kinase domain and shifts the equilibrium toward the catalytically preorganized favored state. Oncogenic mutations of p110α increase membrane recruitment, which is driven by the reorientation of iSH2 and the disengagement of the adaptor-binding domain (ABD)^12^. This suggests that mutation-specific fluctuations on iSH2 and ABD likely attenuate the inhibitory contacts of p85α against p110α. With the membrane-interacting surface of the kinase domain exposed, PI3Kα increases its affinity for the membrane, leading to a population shift toward the catalytically preorganized, membrane-bound open state. This increased affinity stabilizes the primed open state, thereby extending its residence time and enhancing its catalytic action in the conversion of phosphatidylinositol 4,5-bisphosphate (PIP_2_) to phosphatidylinositol 3,4,5-trisphosphate (PIP_3_). PTEN converts PIP_3_ back to PIP_2_^13^.

Here, we provide detailed conformational evolution of PI3Kα mutants in solution and at the membrane. The conformations of oncogenic single hotspot mutants differ from those of weak/moderate mutants, even when combined. We examined two hotspot (driver) mutations, E545K and H1047R, in the helical and kinase domains, respectively. We also considered four weak/moderate mutations in combination with hotspot mutations, R93W in the ABD, E453K in the C2 domain, and E726K and M1043I in the kinase domain. As detailed below, we subscribe to a combination of allosteric drugs with a conformational selection strategy^14–17^, in which the mechanisms of single mutations may complement each other, as with E453K/E545K, or enhance each other’s effects, as with E726K/H1047R. Our comprehensive molecular dynamics (MD) simulations of PI3Kα variants with single and double mutations reveal conformational profiles related to combinations of coexisting mutations. Double mutations can include a hotspot and a weak/moderate mutation of varying strength. Recent computational studies have demonstrated the rescuability of allosteric hotspots^18^. Hotspot mutations can modulate the distinct structural and energetic properties of proteins at distant sites through allosteric coupling. We define mutation strength by the tendency of the conformational ensembles to move toward the active conformation. These profiles can help explain the graded clinical severity of the mutation outcome, ranging from more severe to milder^19–21^, as observed from clinical phenotypic spectrum in biochemical studies^20–22^. Graded clinical phenotypic distributions reproduce the conformational heterogeneity of different combinations of mutations, including single hotspot, weak/moderate mutation, or their combination. We propose designing mutant-selective allosteric drugs that can selectively target the heterogeneous conformations of PI3Kα mutants with high precision. PI3Kα is a large protein and the second most mutated oncogene in cancer. A two-drug approach may be more effective than a single drug against variants with different conformational spectra^23^. This approach is particularly relevant in the context of a combination of allosteric drugs^24^, especially for strong double mutants.

## Results

PI3Kα forms as an obligate dimer containing two subunits, the catalytic p110α and the regulatory p85α. The p110α subunit is encoded by the *PIK3CA* gene located on the 3q26 band of chromosome 3 (**Fig. 1a**). The p85α subunit is encoded by the *PIK3R1* gene located on the 5q13.1 band of chromosome 5 (**Fig. 1b**). *PIK3CA* contains 20 exons with mutation hotspots, exons 9 and 20, which are frequently mutated in various cancers^25^. *PIK3R1* contains 16 exons, and unlike *PIK3CA*, mutations are less frequent and occur in exons 11 and 13^26^. In the cytoplasm, the p110α catalytic subunit binds to the p85α regulatory subunit to form the inactive p85α–p110α complex (**Fig. 1c**). We performed comprehensive computational studies using MD simulations for PI3Kα variants with a single oncogenic hotspot (driver) mutation, a weak/moderate mutation, or a combination of the two. The simulations consisted of inactive PI3Kα in solution and an active PI3Kα at an anionic lipid bilayer, composed of the phosphatidylcholine (PC), phosphatidylserine (PS), and phosphatidylinositol 4,5-bisphosphate (PIP_2_). Seven missense mutations on p110α were examined: R88Q and R93W in the ABD, E453K in the C2 domain, E545K in the helical domain, and E726K, M1043I, and H1047R in the kinase domain. E545K and H1047R are oncogenic driver mutations. Six combinations of double mutations were considered: five for solution simulations (R93W/E545K, E453K/E545K, E545K/M1043I, R93W/H1047R, and E453K/H1047R) and one for membrane simulation (E726K/H1047R). Details of each mutation are summarized in **Supplementary Table 1.**

**Fig. 1.**
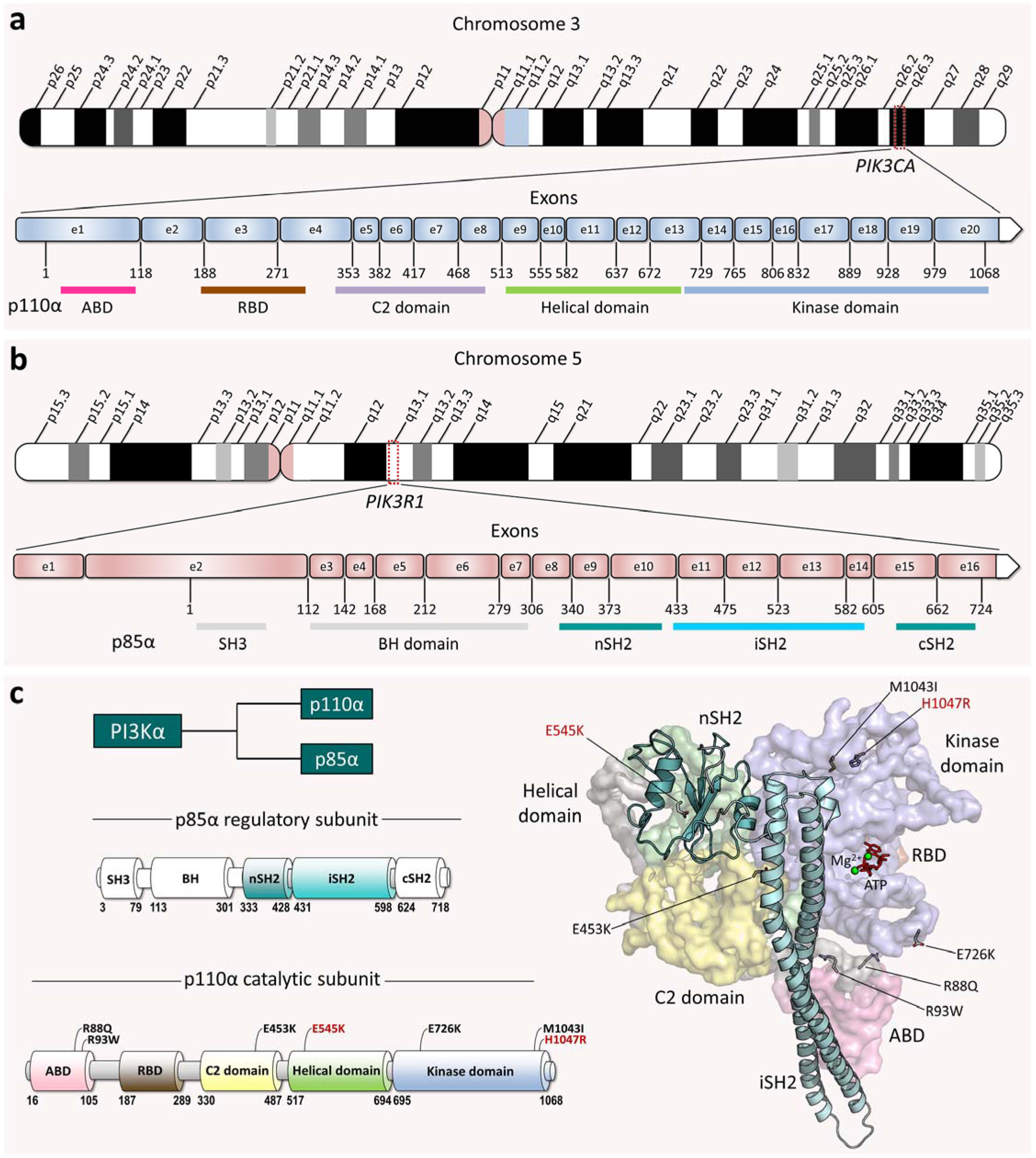
The PI3K*α*-encoding genes (*PIK3CA* and *PIK3R1*) and the protein structure. **a**, Schematic of chromosome 3 with the location of the human *PIK3CA* gene on the 3q26 band. The *PIK3CA* gene contains 20 exons and expresses the p110α catalytic subunit. **b**, Schematic of chromosome 5 with the location of the human *PIK3R1* gene on the 5q13.1 band. The *PIK3R1* gene contains 16 exons and expresses the p85α regulatory subunit. **c**, The domain structures of p85α and p110α (*left panel*). The p110α subunit is composed of ABD (residues 16-105), RBD (residues 187-289), and C2 (residues 330-487), helical (residues 517-694), and kinase (residues 695-1068) domains. The p85α subunit is composed of the SH3 (residues 3-79), breakpoint-cluster region homology (BH, residues 113-301), nSH2 (residues 333-428), iSH2 (residues 431-598), and cSH2 (residues 624-718) domains. The PI3Kα conformation modeled *in silico* (*right panel*), with the mutations mapped onto the structure as marked in the p110α domain structure. The p110α is shown in surface representation with different color codes: The ABD is pink, the RBD is brown, the C2 domain is yellow, the helical domain is green, and the kinase domain is blue. The p85α is depicted as a cartoon, with nSH2 colored light teal and iSH2 colored cyan.

### nSH2 release is a key requirement for PI3Kα activation

Our previous MD simulations provided the mechanism of wild-type PI3Kα activation at the atomic level^8^. PI3Kα is large, so atomistic simulations may not capture all relevant conformational changes related to population shifts due to high kinetic barriers. While this can prevent the evaluation of graded mutational effects, particularly reliable capture of minor conformational changes, the relative stability of the conformations of variants and the wild type can delineate the mutational effects, as discussed below. In our earlier work, we removed the nSH2 to observe the conformational changes^8^. Its absence, as well that of the phosphorylated motif of the C-terminal RTK, biases an accurate comparison of the wild type’s larger population of the closed state with the minor open state species. In this study, the nSH2 of p85α initially interacts with the helical domain of p110α, indicating an inactive PI3Kα. To observe the mutational effects on nSH2 release, we performed simulations of two single mutations, E453K and E545K, and one double mutation with the combination of E453K/E545K. While wild-type PI3Kα (PI3Kα^WT^; WT denotes wild type) retains its nSH2 at the helical domain and remains as an inactive form, PI3Kα mutants, PI3Kα^E453K^, PI3Kα^E545K^, PI3Kα^E453K/E545K^ exhibit large conformational changes of p85α as compared to PI3Kα^WT^, shifting nSH2 for its release (**Supplementary Fig. 1a**). In PI3Kα^WT^, E545^HD^ forms a salt bridge with K379^nSH2^, preventing nSH2 release, and E453^C2^ recruits K942^KD^, which forms a salt bridge with E345^nSH2^, leaving nSH2 intact (**Fig. 2a**). Here, HD and KD denote the helical and kinase domains, respectively. Charge inversion mutations disrupt this salt bridge network. In PI3Kα^E453K^, K453^C2^ repels K942^KD^ and forms a salt bridge directly with E345^nSH2^. In PI3Kα^E545K^, the hotspot mutant residue K545^HD^ loses contact with K379^nSH2^ as expected. While single mutations only affect the interactions at their own mutation sites, the double mutation E453K/E545K completely removes all salt bridges defined in the wild-type system. As a result, an increase in the distance of K379^nSH2^–K545^HD^ at the nSH2/helical domain interface and a decrease in the distance of F934^KD^–K942^KD^ in the activation loop (A-loop) can be observed (**Fig. 2b**). The decreased distance for the pair of residues in the A-loop indicates a conformational change of the loop and a shift towards the active site. A profile of increasing distance for E/K453^C2^–K942^KD^ and E345^nSH2^–K942^KD^ follows the trend, wild type < E545K < E453K/E545K, except for the single mutation E453K (**Supplementary Fig. 1b**). In PI3Kα^E453K^, an alternative formation of the E345^nSH2^–K453^C2^ salt bridge disrupts the interaction of K453^C2^ with K942^KD^ and increases the distances of these residues with K942^KD^. With the substituted salt bridge, PI3Kα^E453K^ can retain the nSH2 at the helical domain as the wild type, indicating the weak mutational effect. Greater displacement of nSH2 by the double mutation E453K/E545K results in a significant reduction in the surface area of the interface between nSH2 and the p110α subunit (**Supplementary Fig. 1c**).

**Fig. 2.**
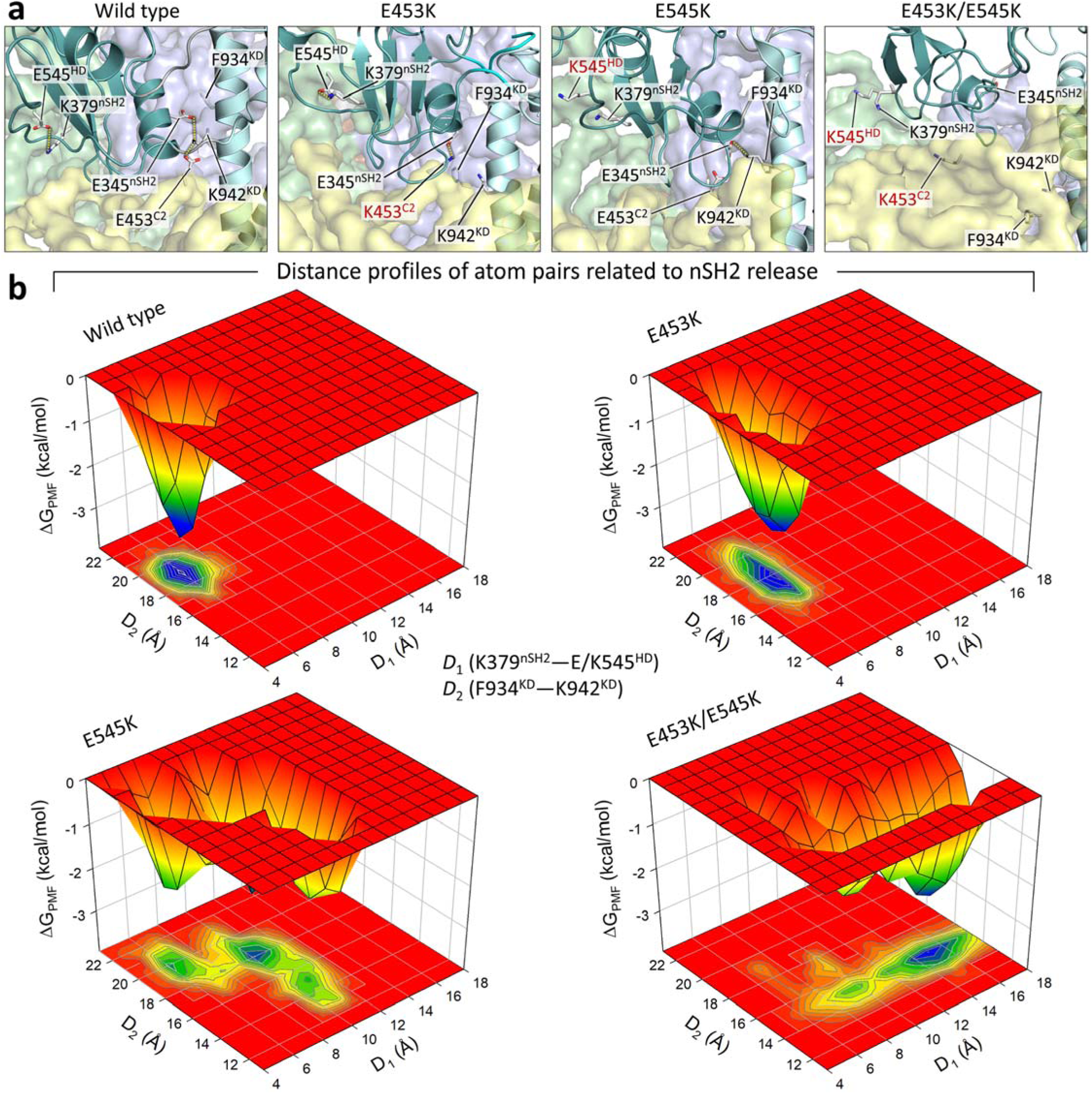
Different strengths of nSH2 release in PI3K*α* with single and double mutations. **a**, The best representative conformations from the ensemble clusters for PI3Kα^WT^, PI3Kα^E453K^, PI3Kα^E545K^, and PI3Kα^E453K/E545K^ are shown, with highlights of the salt bridge interactions at the nSH2/helical/C2/kinase domain interfaces. The p110α structure is shown in surface representation with color codes. The C2, helical, and kinase domains are colored yellow, green, and blue, respectively. The p85α structure is depicted as a cartoon, with nSH2 colored light teal and iSH2 colored cyan. **b**, The three-dimensional potential of mean force, Δ*G*_PMF_, and the projection onto a two-dimensional subspace representing the relative free energy profile along reaction coordinates, *D*_1_ and *D*_2_. In the calculation of the probability distributions for two atom pair distances, *D*_1_ is defined as the distance from K379^nSH2^ to E/K545^HD^, and *D*_2_ is defined as the distance from F934^KD^ to K942^KD^. HD and KD denote the helical and kinase domains, respectively. Unstable profiles of these distances indicate nSH2 release. The double mutation E453K/E545K exhibits pronounced destabilization of these atomic pair distances, demonstrating a more severe effect of the mutation on nSH2 release compared to single mutations or the wild type.

In summary, nSH2 is latched to p110α by salt bridge interactions with E545^HD^ and K942^KD^ in PI3Kα^WT^. Loss of these latches by charge inversion mutations allows nSH2 to be displaced from the p110α subunit (**Supplementary Fig. 2**). In all cases, the double mutation E453K/E545K severely disrupts the salt bridge network at the nSH2/helical/C2/kinase domain interface. This clearly indicates the mutation effect on nSH2 release in the trend: *single weak mutation < single driver mutation < double mutation*. The significant impact of the double mutation on PI3Kα conformation toward activation is consistent with cell experiments that illustrate increased proliferation of *cis* PI3Kα double mutants compared with single hotspot mutants^27^.

### The dynamics of iSH2 due to weakened interdomain interaction and allosteric control leads to PI3Kα activation

The mutation effect can also be observed in the dynamics of iSH2. iSH2 is also latched to p110α by the salt bridge interaction with E453^C2^. However, the E453K mutation destabilizes the interaction of the C2 domain with iSH2, allowing iSH2 to move slightly away from p110α (**Fig. 3a**). In PI3Kα^WT^, E453^C2^ forms a salt bridge R574^iSH2^ or alternatively with K567^iSH2^ and K575^iSH2^. The charge inversion mutation E453K disrupts these salt bridges, resulting in a shift of iSH2 from the C2 domain, which is more pronounced in the double mutation E453K/E545K (**Fig. 3b**). The wild-type residue E453^C2^ restricts the dynamics of the A-loop and iSH2 through salt bridges with K942^KD^ and R574^iSH2^, respectively. The mutant residue K453^C2^ can affect iSH2 dynamics but not nSH2 release. Conversely, the mutant residue K545^HD^ can affect nSH2 release but not iSH2 shift from the C2 domain (**Supplementary Fig. 3**). However, it is important to note that mutations of both E453^C2^ and E545^HD^ to lysine synergically affect nSH2 release and iSH2 shift from the C2 domain.

**Fig. 3.**
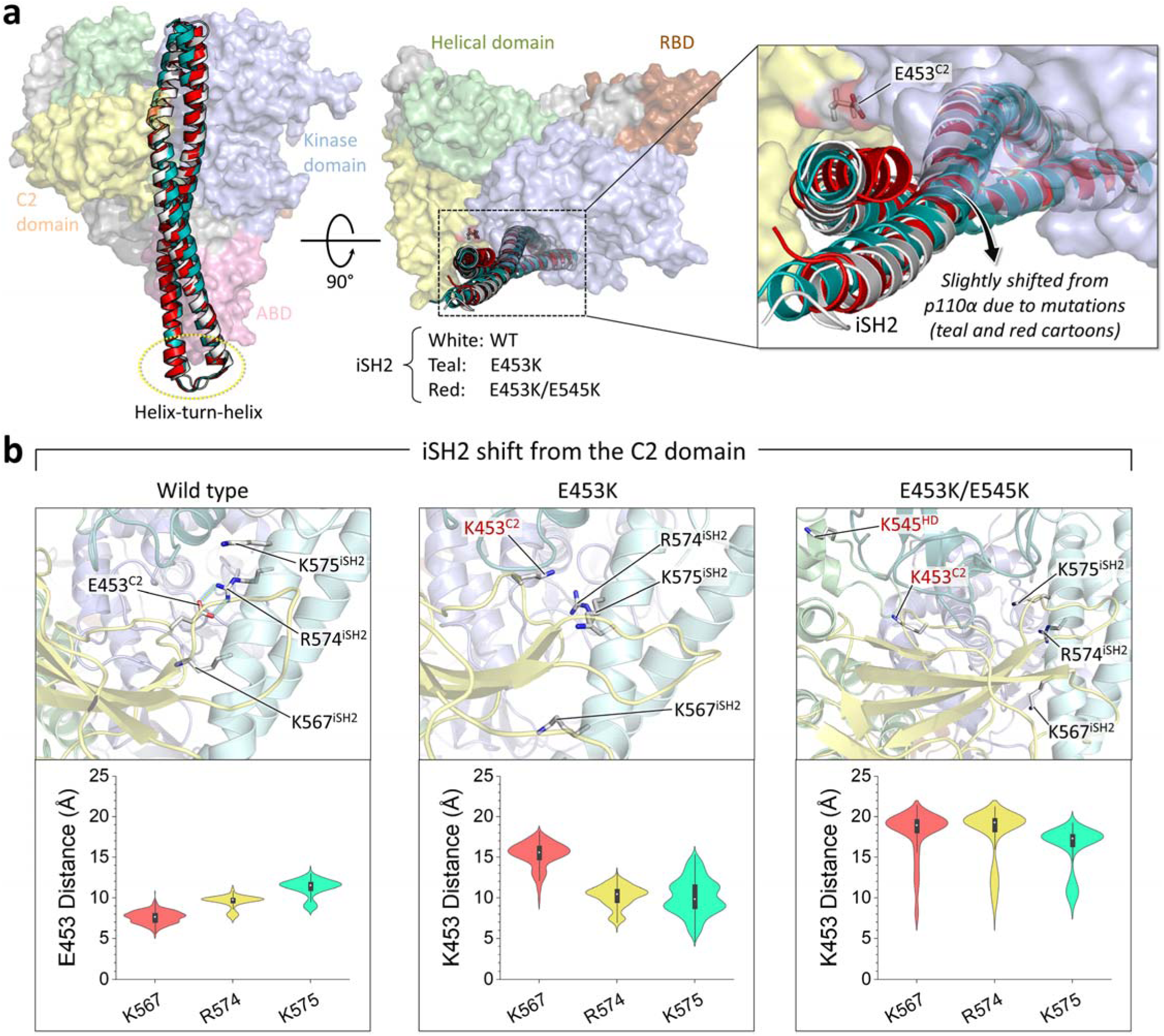
The E453K mutation affects the iSH2 movement. **a**, Superimposition of the best representative conformations of PI3Kα^WT^, PI3Kα^E453K^, and PI3Kα^E453K/E545K^ highlighting iSH2 movement. The p110α is shown in surface representation with color codes: The ABD is pink, the RBD is brown, the C2 domain is yellow, the helical domain is green, and the kinase domain is blue. The iSH2 structure is depicted as a cartoon, showing the wild type in white, the single mutation E453K in teal, and the double mutation E453K/E545K in red. **b**, Highlights of the salt bridge interactions at the iSH2/C2 domain interfaces are shown alongside violin plots representing the atomic pair distances of E/K453^C2^–K567^iSH2^, E/K453^C2^–R574^iSH2^, and E/K453^C2^– K575^iSH2^. The protein structure is depicted as a cartoon: the C2 domain is yellow, the helix domain is green, the kinase domain is blue, and iSH2 is cyan. The double mutation E453K/E545K exhibits a greater synergistic effect on the iSH2 shift followed by the nSH2 release. The iSH2 shift relieves the constraint on the collapsed A-loop conformation, promoting its exposure.

The iSH2 domain in the regulatory subunit of p85α is a long coiled-coil with the motif of two long α-helices coiled together. The iSH2 helices extend from the ABD and reach the interface between the kinase domain and the helical and C2 domains. The helix-turn-helix region of iSH2 interacts with the ABD, and the R93W mutation on the ABD can disrupt this interaction. Unlike E453^C2^, which is in direct contact with iSH2, R93^ABD^ is distal to iSH2 (**Fig. 4a**). In PI3Kα^WT^, R93^ABD^ forms a salt bridge with E710^KD^, thereby attaching the ABD to the kinase domain (**Fig. 4b**). In PI3Kα^R93W^, W93^ABD^ removes the interaction with the kinase domain by replacing the salt bridge to π-π stacking with F119^ID^ in an interdomain (ID) helix (residues 108-122) (**Fig. 4c**). These local changes in the interaction can cause fluctuations in the helix-turn-helix region of iSH2. While the conformational changes for the single mutation R93W appear to be minor or wild-type-like behavior, the large deviation of the helix-turn-helix region of iSH2 for the double mutation R93W/E545K is eminent compared to the wild type (**Supplementary Fig. 4a**). Consistent with this, greater fluctuations in ABD are observed in R93W/E545K compared to both wild type and R93W (**Fig. 4d**). It appears that the single hotspot mutation E545K alone can disrupt the R93^ABD^–E710^KD^ salt bridge and slightly increase the fluctuations of ABD compared to the wild type, but not as much as the double mutation R93W/E545K (**Supplementary Fig. 4b**). To observe how E545^HD^ allosterically controls the ABD conformation at such a distal site, we identified the optimal propagation pathways through the protein by calculating the dynamic correlated motion among residues using the weighted implementation of suboptimal paths (WISP) algorithm^28^. In PI3Kα^WT^, the iSH2 regulates p110α autoinhibition by allosterically constraining the domains in the p110α subunit. It can be observed that both PI3Kα^WT^ and PI3Kα^R93W^ propagate the allosteric signal between E545^HD^ and R/W93^ABD^, connecting the signaling nodes in nSH2, iSH2, and ABD (**Fig. 4e**). However, PI3Kα^R93W/E545K^ disrupts the propagation of the allosteric signal between K545^HD^ and W93^ABD^. Instead, it establishes an alternative allosteric pathway between K545^HD^ and E710^KD^ through the kinase domain nodes, which bypasses the signaling nodes in nSH2 and iSH2, altogether suggesting weakened interdomain associations between nSH2 and helical domain and between iSH2 and ABD.

**Fig. 4.**
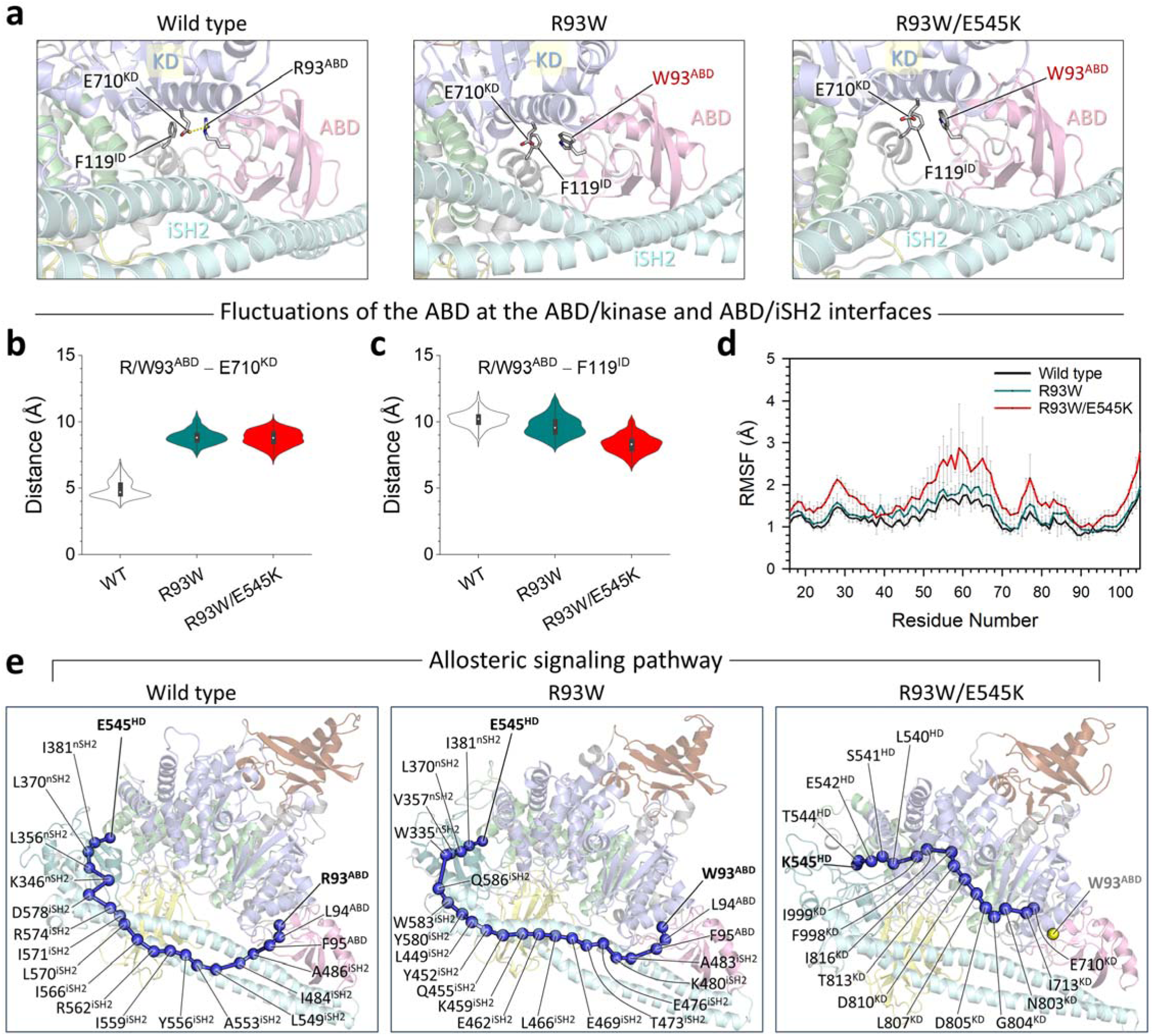
The R93W mutation disrupts the ABD/kinase domain interface, inducing movement of the helix-turn-helix region of iSH2. **a**, The best representative conformations from the ensemble clusters for PI3Kα^WT^, PI3Kα^R93W^, and PI3Kα^R93W/E545K^ are shown, with highlights of the residue interactions at the ABD/kinase domain interfaces. The protein structure is depicted as a cartoon with the different colors: the ABD is pink, the helix domain is green, the kinase domain is blue, and iSH2 is cyan. **b**, Violin plot representing the atomic pair distance of R/W93^ABD^–E710^KD^, and **c**, the same of R/W93^ABD^–F119^ID^. KD and ID refer to the kinase domain and interdomain, respectively. **d**, The root-mean-squared-fluctuations (RMSFs) of the ABD for PI3Kα^WT^ (black), PI3Kα^R93W^ (teal) and PI3Kα^R93W/E545K^ (red). **e**, The allosteric signaling pathways between the source residue E/K545^HD^ and the sink residue R/W93^ABD^. Blue beads denote the allosteric signal nodes, and yellow bead for W93^ABD^ in the R93W/E545K double mutation indicates the broken allosteric signal. The greater fluctuations observed in the ABD of PI3Kα^R93W/E545K^ indicate a loss of interaction with the kinase domain as well as with iSH2, inducing movement of the helix-turn-helix region of iSH2. This results in disruption of the allosteric signals passing through iSH2, in turn, abolishing the regulatory role of iSH2, which allosterically acts in p110α autoinhibition.

As with R93W, the single mutation R88Q disrupts the interaction between the ABD and the kinase domain. In PI3Kα^WT^, R88^ABD^ forms a salt bridge with D746^KD^, or occasionally with D743^KD^, attaching the ABD to the kinase domain (**Supplementary Fig. 5**). However, R88Q disrupts the interaction with the kinase domain and increases the fluctuations of ABD to as much as those of the double mutation R93W/E545K. These fluctuations may interfere with the interaction of the ABD with the helix-turn-helix region of iSH2, leading to instability of p85α in its regulatory role. PI3Kα^WT^ propagates the allosteric signal between E545^HD^ and R88^ABD^, connecting the signaling nodes in nSH2, iSH2, and ABD, which is similar to the allosteric pathway between E545^HD^ and R/W93^ABD^ for PI3Kα^WT^ and PI3Kα^R93W^. However, PI3Kα^R88Q^ disrupts the propagation of the allosteric signal between E545^HD^ and R88^ABD^. An alternative allosteric pathway between E545^HD^ and D746^ABD^ through the kinase domain nodes fails to connect ABD as observed in PI3Kα^R93W/E545K^. Bypassing the allosteric signaling nodes in nSH2 and iSH2 indicates that single mutation R88Q alone is capable of inducing iSH2 instability as the double mutation R93W/E545K, suggesting a stronger mutational effect than R93W, the same ABD mutation.

### Protruding A-loop may facilitate substrate recruitment and promote membrane anchoring of kinase domain

The A-loop in the kinase domain of PI3Kα is highly basic and contains basic boxes, the first with the _941_KKKK_944_ and the second with the _948_KRER_951_ motifs. These basic residues are involved in electrostatic interactions with the nSH2 acidic motif (E341, E342, E345) and with iSH2 residue D464. The strong double mutation E453K/E545K is able to disrupt the C2/nSH2/kinase domain interface, allowing the conformational change of the A-loop (**Supplementary Fig. 2**). Similarly, the conformational change of the A-loop can be observed with the M1043I mutation on kα11 in the regulatory arch. In PI3Kα^WT^, M1043^kα11^ retains the hydrophobic cluster with adjacent hydrophobic residues, V906^kα6^, F909^kα6^, V952^A-loop^, V955^A-loop^, F1039^kα11^, and W1051^kα12^ (**Fig. 5a**). The latter part of the A-loop has the U-motif, and activation involves loss of the U-motif in the A-loop through reorientation of the U-motif residue^29^, F954^A-loop^. The M1043I mutation preserves the hydrophobic cluster with the same residues (**Supplementary Fig. 6a**), except for W1051^kα12^. Large fluctuations of W1051^kα12^ occur in the presence of the M1043I mutation, with even more pronounced effects in the case of the double mutation E545K/M1043I (**Fig. 5b**). The fluctuations may cause the latch of K942^A-loop^ on nSH2 to loosen, resulting in conformational change of the A-loop (**Fig. 5c**). In PI3Kα^WT^, M1043^kα11^ constrains K942^A-loop^ by propagating the allosteric signal through the first basic box residue (**Supplementary Fig. 6b**). However, in the M1043I mutation, I1043^kα11^ propagates the allosteric signal to K942^A-loop^ through other A-loop residues instead of the first basic box residues, and in the double mutation E545K/M1043I, I1043^kα11^ propagates the allosteric signal to K942^A-loop^ bypassing the A-loop residues. These distinct allosteric signaling networks caused by mutations are responsible for the protrusion of the A-loop from the kinase domain surface (**Fig. 5d**). A prominent protrusion of the first basic box and an adjacent hydrophobic residue F945^A-loop^ is observed in both single and double mutations (**Fig. 5e**). We propose that the first basic box of the protruding A-loop plays an important role in recruiting the lipid substrate to the catalytic site.

**Fig. 5.**
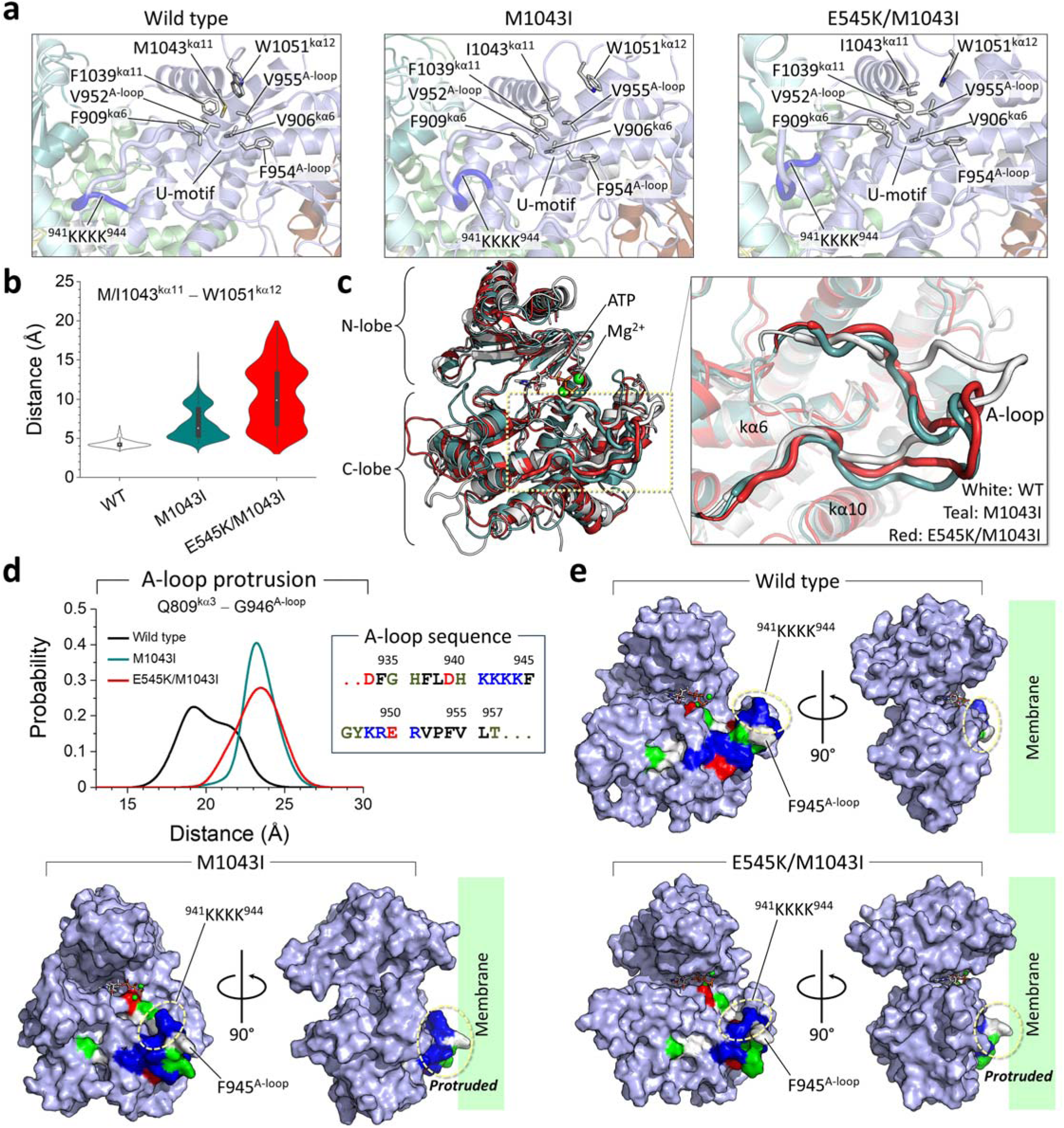
The M1034I mutation causes the A-loop to protrude. **a**, The best representative conformations from the ensemble clusters for PI3Kα^WT^, PI3Kα^M1043I^, and PI3Kα^E545K/M1043I^ are shown, with highlights of the residue interactions in the kinase domain. **b**, Violin plot representing the atomic pair distance of M/I1043^kα11^–W1051^kα12^. **c**, Superimposition of kinase domains from the best representative conformations highlights A-loop conformations. Wild-type is shown in white, the single mutation M1043I is shown in teal, and the double mutation E545K/M1043I is shown in red. **d**, The probability distribution of the distance between Q809^kα3^ and G946^A-loop^ representing the A-loop protrusion. The sequence of A-loop highlighting the first _941_KKKK_944_ and the second _948_KRER_951_ basic boxes. **e**, The kinase domain structures are highlighted with A-loop structures. The kinase domain is colored cyan. In the A-loop structure, residues are colored white, green, blue, and red to indicate their hydrophobic, polar, positively charged, and negatively charged properties, respectively. The protruding region of the A-loop toward the pseudo-membrane is marked. Both the single M1043I and the double E545K/M1043I mutations induce conformational change in the A-loop, suggesting that the M1043I mutation on kα11 can cause the A-loop to protrude. This A-loop protrusion from the kinase domain surface exposes the first basic box _941_KKKK_944_, facilitating the recruitment of the lipid substrate.

A similar conformational change of the A-loop is also observed for the E545K and E453K/E545K mutations (**Supplementary Fig. 7a**). It appears that the single hotspot mutation E545K alone can affect the conformation of the A-loop by protruding the first basic box from the kinase domain surface. M1043^kα11^ propagates the allosteric signal to K942^A-loop^ bypassing the first basic box residues (**Supplementary Fig. 7b**). The strong double mutation E453K/E545K also protrudes the first basic box of the A-loop (**Supplementary Fig. 7c**). In this case, only a weak allosteric signal from M1043^kα11^ constrains K942^A-loop^, and the release of nSH2 by the hotspot mutation removes the constraint on the A-loop, liberating it from the C2/nSH2 interface (**Supplementary Fig. 2**).

The conformational dynamics of the oncogenic hotspot mutation H1047R in kα11 of the C-terminal kinase domain differ from that of the oncogenic hotspot mutation E545K in the helical domain. Genetic and biochemical analyses suggested that mutations in the helical and kinase domains trigger a gain of p110α function through different molecular mechanisms^30^. Helical domain mutation E545K is not influenced by p85α binding but is dependent on Ras for its functionality. In contrast, kinase domain mutation H1047R, which replaces Ras binding, requires allosteric change mediated by p85α. To observe the effects of the H1047R mutation on PI3Kα activation, we considered double mutations with the combinations of R93W/H1047R and E453K/H1047R—similar combination of weak/strong mutations as performed for E545K. Unlike E545K, which uncouples nSH2 from the helical domain of p110α, H1047R is less effective in releasing nSH2 as a single mutation or in combination with R93W or E453K (**Supplementary Fig. 8a**). In addition, H1047R is also less effective for iSH2 movement, even in the double mutations (**Supplementary Fig. 8b**). However, we observed that H1047R slightly induces the A-loop protrusion as does the M1043I mutation, which is located at the same kα11 (**Supplementary Fig. 8c**). Based on our observations, we suggest that the H1047R mutation does not exert both nSH2 release and iSH2 movement but slightly affects the A-loop dynamics during the inactive-to-active transition in the p110α conformation.

### Mutations shift the equilibrium toward the catalytically favored state at the membrane

Oncogenic mutations in p110α facilitate PI3Kα activation by bypassing intrinsic constraints imposed by p85α. These include p85α nSH2 release and iSH2 movement, resulting in a significant conformational change of p110α to the active conformation. As we observed above, unlike other oncogenic hotspot mutations, H1047R is less effective in eliminating intrinsic constraints exerted by p85α. It was known that H1047R can regulate the membrane engagement of the active form of PI3Kα. To prove this, we performed simulations of active PI3Kα at the membrane. Our membrane-bound PI3Kα results implied that SH2 domains are recruited by the C-terminal phosphorylated tyrosine (pY) motifs of the RTK, leading to the active form of PI3Kα. Single-molecule total internal reflection fluorescence microscopy (TIRFM) measurements revealed that PI3Ks prioritize interaction with the pY motif, leading to SH2 release, before localizing to the membrane^31,32^. To take this into account, we adopted an active conformation of PI3Kα from our previous simulations^8^ (**Supplementary Fig. 9**). The active form of PI3Kα has both SH2 domains removed, allowing us to observe the structural rearrangement due to mutations via localization and orientation at the membrane. Here we consider the single oncogenic hotspot mutation H1047R and the combination of the hotspot mutation with the weak mutation E726K. In the presence of the membrane, PI3Kα diffuses to the membrane through contact of the p85α subunit with its long coiled-coil on the membrane surface and three anchor points of the p110α subunit located on the C2 domain and the N- and C-lobe kinase domains (**Supplementary Fig. 10**). We observed that all PI3Kα systems, whether wild type or mutants, which are all active forms, maintain these three anchor points when interacting with the membrane. For the C2 domain, the basic residues R349 in the C2β1-C2β2 loop and R412 and K413 in the C2β5-C2β6 loop form an anchor point by diffusing their sidechains into the amphipathic interface of the lipid bilayer. For the N-lobe kinase domain, the tandem lysine residues K723-K724 and D725 in the kα1-kα2 loop serve as an anchor point (**Fig. 6a**). For the C-lobe kinase domain, the hydrophobic residues W1057, I1058, and F1059 in kα12 form an anchor point by immersing their sidechains into the hydrophobic core of the lipid bilayer. However, the positions of these anchor point residues with respect to the membrane surface and some basic residues close to the active site show different patterns depending on the mutation profile (**Fig. 6b**). In PI3Kα^WT^ and PI3Kα^H1047R^, E726^kα1-kα2^ is close to the membrane surface, whereas in PI3Kα^E726K^ and PI3Kα^E726K/H1047R^, its mutant residue K726^kα1-kα2^ is slightly more distal from the membrane surface than its wild-type form. Similarly, in PI3Kα^WT^ and PI3Kα^E726K^, H1047^kα11^ is largely distal to the membrane surface, while in PI3Kα^H1047R^ and PI3Kα^E726K/H1047R^, its mutant residue R1047^kα11^ is closer to the membrane surface than its wild-type form. Surprisingly, R1047^kα11^ is not directly involved in interactions with any lipid, including PIP_2_, contrary to expectations for PIP_2_ recruitment. Instead, the H1047R mutation destabilizes the interaction of the kα11 helix with the kα7 and kα8 helices. In PI3Kα^WT^, these helical interactions are stabilized by the polar contact of H1047^kα11^ with Q958^kα7^ and π-π stacking of H1047^kα11^ with F977^kα8^. Uncoupling kα12 from the kα11 helix occurs dominantly with the H1047R mutation due to the destabilized helical interactions. This causes kα12 to move further toward the membrane than the wild type, facilitating the diffusion of the hydrophobic residues W1057, I1058, and F1059 in kα12 to the membrane, as reported as the C-terminal “closed-to-open” transition^33^. This suggests that the function of the H1047R mutation is to provide a stable C-lobe anchor point, thereby replacing the role of Ras, since the RBD is located above the C-lobe kinase domain from the membrane.

**Fig. 6.**
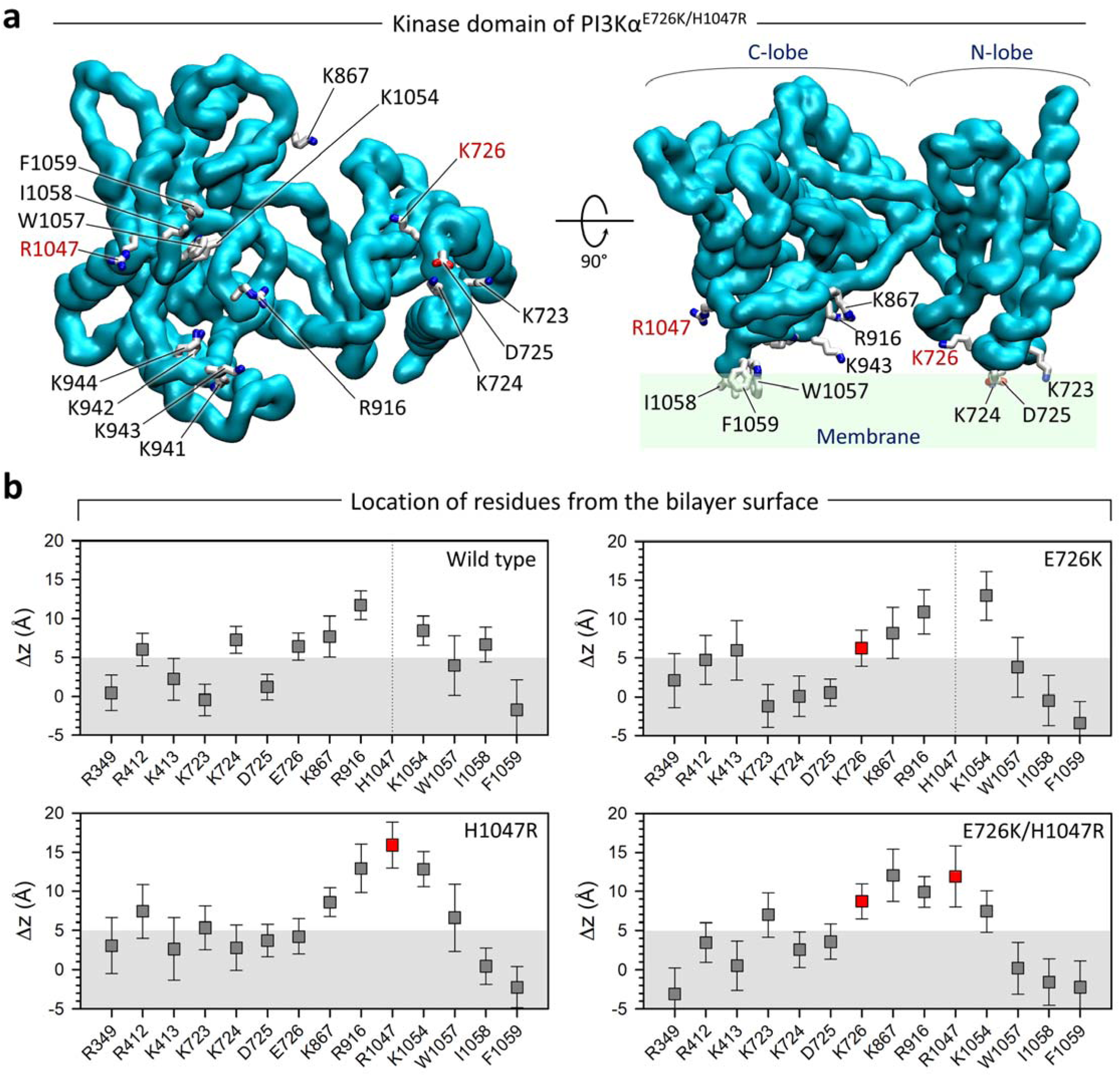
Active PI3K*α* interacts with the membrane through three anchor points. **a**, Mapping of the residues on the membrane-binding surface of the kinase domain of active PI3Kα^E726K/H1047R^. The mutant residues K726 and R1047 are marked in red. **b**, Averaged deviations of the residues from the bilayer surface for the active PI3Kα^WT^, PI3Kα^E726K^, PI3Kα^H1047R^, and PI3Kα^E726K/H1047R^. Red boxes denote the mutant residues K726 and R1047. The greater diffusion of the hydrophobic residues of kα12 (W1057, I1058, and F1059) into the hydrophobic core of the lipid bilayer suggests that the double mutation E726K/H1047R provides a more secure membrane anchor point at the C-lobe kinase domain than single mutations or the wild type. This secure anchor point in the C-lobe kinase domain can bring the active site closer to the membrane surface.

### The problem: PI3Kα catalytic action requires phosphate transfer, but the γ-phosphate (Pγ) of ATP is ∼19 Å from the membrane surface!

In lipid catalysis, PI3Kα phosphorylates PIP_2_ to form PIP_3_ by transferring a phosphate group from ATP. The substrate PIP_2_ must be close to the γ-phosphate of ATP for catalysis to occur, but the average position of the γ-phosphate (Pγ) of ATP is ∼19 Å from the membrane surface (**Supplementary Fig. 11**), which is too far for catalysis. However, our data show that PIP_2_ protrudes significantly from the membrane at the holo bilayer leaflet containing the protein compared to the apo bilayer leaflet (**Supplementary Fig. 12a**). In mutant systems, the phosphate group of PIP_2_ is located ∼3 Å above the average position of the phosphate groups of PC and PS, and it can move up to ∼8 Å as seen in the distribution. With the large size of the head group of the phosphorylated inositol ring, which has a size of ∼8.3 Å between the phosphate group and the phosphate at the 4th position of the inositol ring, the protrusion allows PIP_2_ to be close enough to ATP for catalysis. PIP_2_ protrusion occurs at the region of the membrane surface facing the active site of the kinase domain, followed by its recruitment by strong electrostatic attraction of the _941_KKKK_944_ motif in the A-loop and K867^kα4^ and R916^kα6-kβ9^ near the active site (**Supplementary Fig. 12b**). The more PIP_2_ coordination at the active site, the more pronounced the protrusion. The protrusion occurs during the simulations. In the double mutant simulation, for example, the coordination distance for a particular PIP_2_ at the starting point is ∼10 Å (**Supplementary Fig. 13a**). This PIP_2_ is observed as close as ∼5 Å in the simulation (**Fig. 7a**). The coordination distance for PIP_2_ in the wild-type system is ∼8 Å, suggesting that the mutations promote PIP_2_ protrusion and coordination. The radial distribution function, *g*(*r*), of the hydroxyl group at the 3rd position of the inositol ring of PIP_2_ with respect to the Pγ of ATP shows that the double mutation E726K/H1047R is better at coordinating PIP_2_ to ATP than the single mutations H1047R and E726K or the wild type (**Fig. 7b**). Similarly, PIP_2_ coordination at the _941_KKKK_944_ motif is better in the double mutation system than in the single mutation or wild-type systems. Mutant residue K726^kα1-kα2^ in PI3Kα^E726K^ and PI3Kα^E726K/H1047R^ is able to recruit PIP_2_, but its wild-type residue E726^kα1-kα2^ in PI3Kα^H1047R^ and PI3Kα^WT^ fails to coordinate PIP_2_. Near the active sites, all systems show PIP_2_ coordination at K867^kα4^ and R916^kα6-kβ9^ (**Supplementary Fig. 13b**). The ability of PI3Kα to bind an additional PIP_2_ molecule was confirmed *in vitro* by fluorescence quenching experiments^34^. Overall, the substrate coordination of the mutant systems is pronounced compared to the wild-type system. This suggests that the oncogenic hotspot H1047R mutation promotes the C-lobe kinase domain to contact the membrane and the weak E726K mutation increases the population of PIP_2_ at the active site, adding the role of substrate recruitment to the _941_KKKK_944_ motif.

**Fig. 7.**
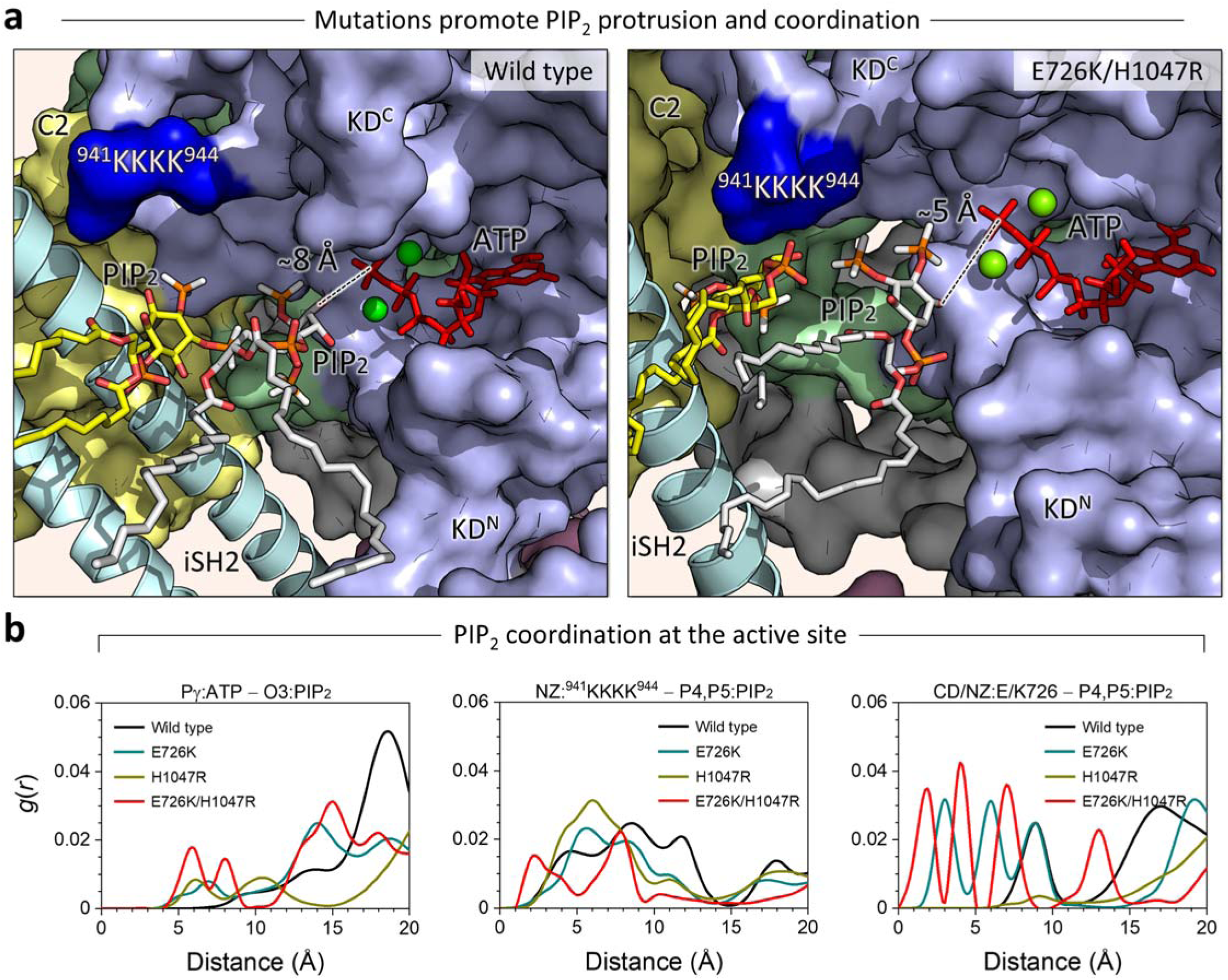
PI3K*α* mutations promote the coordination of PIP_2_ to the active site. **a**, A side-by-side comparison of the PIP_2_ protrusion and coordination. Snapshots highlighting the active sites of PI3Kα^WT^ and PI3Kα^E726K/H1047R^ from simulations. The coordination distances for a particular PIP_2_ are ∼8 Å and ∼5 Å for the wild-type and the double mutant systems, respectively. The coordination distance is defined as the distance between the hydroxyl group at the 3rd position of the inositol ring of PIP_2_ and the Pγ of ATP. The p110α structure is shown in surface representation with different color codes: The C2 domain is yellow, the helical domain is green, and the kinase domain is blue. The p85α structure is depicted as a cartoon. PIP_2_ is shown as sticks. ATP is shown as red sticks, and Mg^2+^ is shown as a green sphere. C2 and HD denote the C2 and helical domains, respectively. KD^N^ and KD^C^ represent the N-lobe and C-lobe kinase domain, respectively. The dark blue surface highlights the _941_KKKK_944_ motif in the A-loop. **b**, The radial distribution function, *g*(*r*), of selected atom pairs for PI3Kα^WT^, PI3Kα^E726K^, PI3Kα^H1047R^, and PI3Kα^E726K/H1047R^, representing the PIP_2_ coordination at the active site of the kinase domain. The distal location of ATP (∼19 Å) can be overcome by extracting the substate from the membrane, which enables its catalytic activity. The PIP_2_ extraction process begins with the initial recruitment of the head group by the _941_KKKK_944_ motif, which then protrudes it from the membrane through A-loop dynamics. The double mutation E726K/H1047R accelerates this process compared to single mutations or the wild type.

*In summary, how then is the ∼19 Å distance of ATP from the membrane surface spanned for the crucial phosphate transfer reaction*? Our results suggest that the _941_KKKK_944_ motif initially recruits the phosphorylated inositol ring of PIP_2_ via strong electrostatic attraction, extracting it from the membrane surface (**Supplementary Fig. 13a** and **Fig. 7a**). Subsequent A-loop fluctuations promote the protrusion of PIP_2_ from the bilayer surface and deliver it to the active site. The double mutation E726K/H1047R is more effective than single mutations or the wild type in recruiting, protruding, and coordinating PIP_2_ at the active site. This indicates that the impact of mutations on substrate recruitment is in the following order: *single weak mutation < single driver mutation < double mutations*. Our findings are consistent with experimental observations from liposome sedimentation assays. These experiments showed that *cis* PI3Kα double mutants exhibit increased binding to anionic and PIP_2_ liposomes compared to their respective single mutants^27^.

### PI3Kα double mutations are combinations of hotspot and weak/moderate mutations

Some genes can have multiple non-synonymous mutations in the same tumor^27,35,36^. These composite mutations combine frequent mutations with rare or weak mutations in the same gene. Our in-depth statistical analysis of *PIK3CA* (see the Methods section for details) revealed that a large number of tumors harbor mutations at following positions: H1047, E545, E542, R88, E726, M1043, E453, R93 in order of their frequencies (**Supplementary Fig. 14a**). Strong driver mutations occur at H1047, E545, and E542; weak driver mutations occur at R88, M1043, E453, and R93; and E726 is a strong latent driver. Strong driver mutations at E542, E545, and H1047 occur in >10% of all *PIK3CA* mutant tumors (*n* = 7284). Our analysis revealed 23 double mutations in the *PIK3CA* gene (**Supplementary Fig. 14b**). The strong latent driver mutation at E726 forms three double mutations with the strong driver mutations at H1047, E545, and E542. These mutations are present in 22, 22, and 20 tumors, respectively. Strong driver mutations in different domains can form double mutations with each other, such as E542/H1047 and E545/H1047. However, the odds ratio calculation implies that these doublets are mutually exclusive. Their relative rarity, particularly those involving two strong drivers, can be explained by the expectation that strong signals may result in oncogene-induced senescence (OIS). *PIK3CA* can be activated by combinations of strong and weak drivers. The strong driver mutation at H1047 is coupled with weak driver mutations at R108, V344, E365, and P539. Another strong driver at E542 forms a double mutation with the weak mutation at E453. The strong driver mutation at H1047 also forms doublets with the weak latent driver mutations at N107 and H1048. The nine co-occurring weak driver/weak driver combinations are R88/R108, R88/M1043, R88/T1025, R88/D350, R88/E81, R88/Y1021, R93/Y1021, R38/R108, and V344/T1025. There are two weak driver/weak latent driver combinations: R88/R357 and R93/E418. However, these weak combinations are not expected to fully activate *PIK3CA*. Overall, double mutations are rare, occurring in only ∼1% of *PIK3CA* mutant tumors.

### Discovering cryptic allosteric pockets of PI3Kα for mutant-selective drug strategy

A cryptic allosteric pocket, which is not visible in a protein in experimental structures, is a hidden binding site that can be exposed during protein dynamics. In principle, these cryptic pockets may be candidates for drug discovery. Using the PockDrug program^37^, we discovered multiple cryptic allosteric pockets, including gaps and cavities within the protein domains during the simulations. First, we selected pockets by screening them with a druggable probability of *p* > 0.5, removing larger pockets spanning the gaps between domains and smaller pockets comprising less than 14 residues, which yields a possible set of allosteric cryptic pockets allowing us to evaluate them (**Supplementary Fig. 15a**). We ignored pockets located within the cavities of the domains, such as the ABD, C2, or nSH2. For the strong double mutation E453K/E545K, we finally obtained two cryptic allosteric pockets, pocket #2 and #6 (**Fig. 8a and Supplementary Table 2**). Pocket #2 is encompassed by the C2, helical, kinase domains of p110α and iSH2 of p85α. An allosteric drug in this pocket can restrict the A-loop and iSH2 dynamics, preventing the kinase domain from being exposed to the membrane. Pocket #6 is encompassed by the C2, helical, and kinase domains, including the mutated residue K453^C2^. The fluorescence polarization assay combined with SiteMap prediction revealed an allosteric binding site encompassed by the nSH2, C2, and helical domains^38^, which is similar to Pocket #6. An allosteric drug in this pocket may constrain the C2 domain dynamics, thereby restricting the iSH2 movement. For the strong double mutation of PI3Kα, a combination of two allosteric drugs, each targeting a specific mutation, can be highly effective. For the double mutation R93W/E545K, we also obtained two possible cryptic allosteric pockets, pocket #2 and #48 (**Fig. 8b and Supplementary Table 3**). Interestingly, pocket #2 for R93W/E545K is located at the same site as in E453K/E545K, suggesting that the pocket is specific for the E545K hotspot mutation. Similarly, the E545K-derived pocket, pocket #4 (**Supplementary Table 4**), is also observed in the E545K single mutation system, but not in the E453K single mutation system, where pocket #10 (**Supplementary Table 5**) is predicted at the site between the C2 and kinase domains (**Supplementary Fig. 15b**). Pocket #48 is R93W specific because it is surrounded by the helix-turn-helix motif of iSH2 and ABD, where the W93^ABD^ mutation is located. An allosteric drug in this pocket may reduce the mutation-induced fluctuation of ABD and restore the wild-type-like allosteric pathway through iSH2 and nSH2 to the helical domain (**Fig. 4e**), leading to an enhanced regulatory effect of p85α. An allosteric binding site at the interface between the kinase domain and the ABD was also reported^38^, which is similar to Pocket #48. For PI3Kα^WT^, we obtained one possible cryptic allosteric pocket in the kinase domain, pocket #9 (**Fig. 8c and Supplementary Table 6**). The location of pocket #9 partially overlaps that of the allosteric drug RLY-2608 embedded in the crystal structure of inactive wild-type PI3Kα (PDB ID: 8TSD). This pocket #9 is wild-type specific; no similar pocket was predicted in PI3Kα mutant systems.

**Fig. 8.**
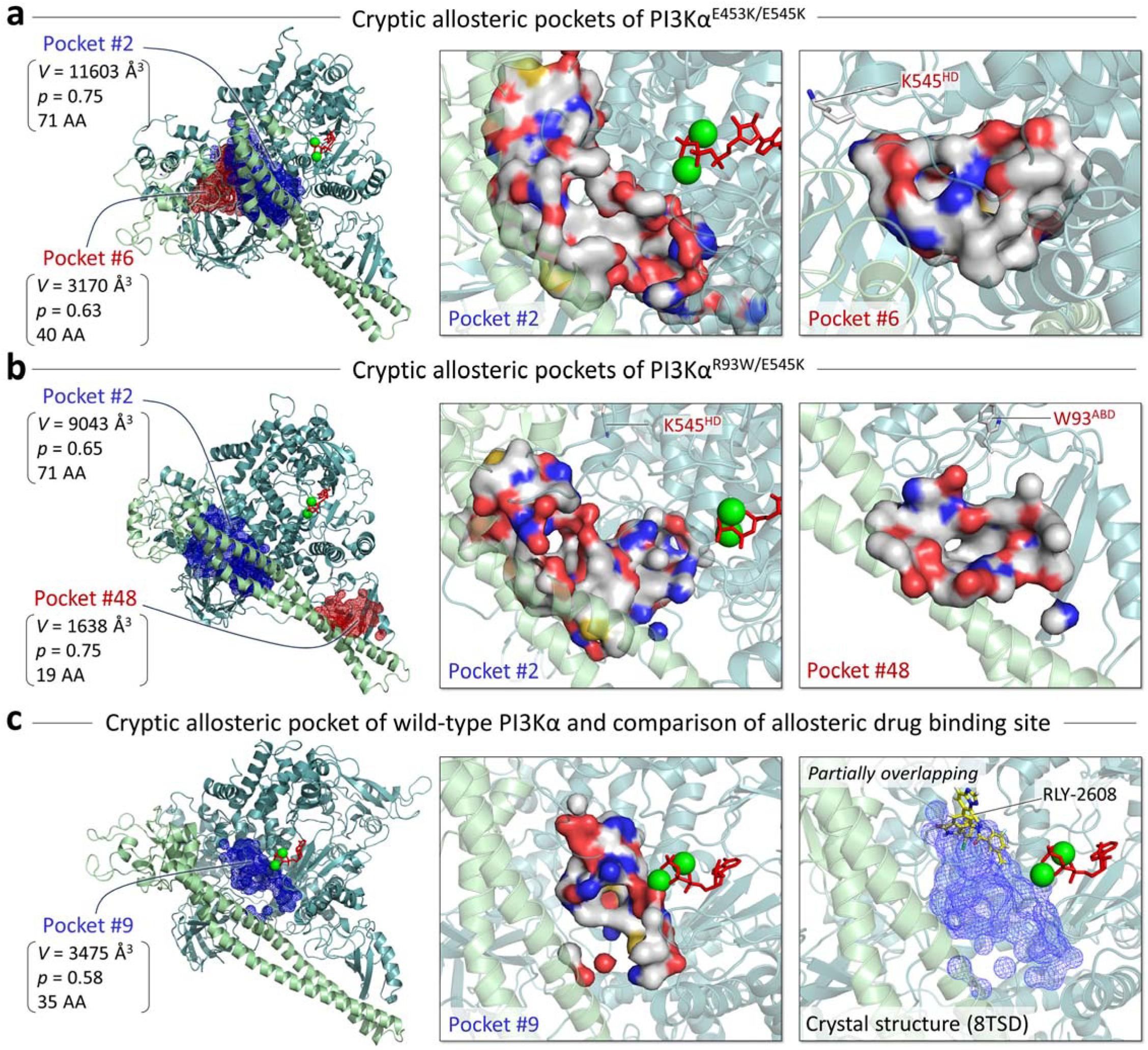
Mutant-specific cryptic allosteric pockets discovered during the dynamics of protein. **a**, Two cryptic allosteric pockets with high probability were predicted for PI3Kα^E453K/E545K^ (*left*). The blue mesh represents the pocket with the highest probability, and the red mesh represents the next highest probability pocket. The pocket volume, probability, and number of amino acids involved in the pocket are indicated. Highlights of these pockets are shown in the surface representation (*middle and right*). In the pocket structure, residues are colored white, green, blue, and red to indicate their hydrophobic, polar, positively charged, and negatively charged properties, respectively. **b**, The same for PI3Kα^R93W/E545K^. **c**, A predicted cryptic allosteric pocket with a high probability for PI3Kα^WT^ (*left*), highlight of the pocket in the surface representation (*middle*), and overlay of the allosteric drug RLY-2608 in the pocket. The cryptic allosteric pockets that we discovered are highly mutation-specific. As such, they can help design allosteric drugs that precisely target oncogenic variants. Combining allosteric drugs can target PI3Kα variants with double mutations.

Another oncogenic hotspot H1047R generates its own unique pocket in the C-lobe kinase domain of p110α. For the oncogenic hotspot mutation H1047R, whether in the double mutants PI3Kα^E453K/H1047R^ and PI3Kα^R93W/H1047R^, or in the single mutant PI3Kα^H1047R^, a cryptic allosteric pocket occurs in the region surrounded by kα6, kβ9, kβ10, kα7, kα8, and kα11 (**Supplementary Fig. 16a and Supplementary Tables 7-9**). The pocket can be labeled as H1047R specific. In PI3Kα^R93W/H1047R^, another deep allosteric pocket can be observed in the region encompassed by kα3, kα6, kα8, kα9, and kα10. In PI3Kα^WT^, the allosteric inhibitor PIK-108 was reported to bind to this region surrounded by kα6, kα7, kα8, and kα11 (PDB ID: 4A55)^39,40^. This pocket is very similar to our predicted H1047R-specific pocket, although slightly deeper (**Supplementary Fig. 16b**). Free energy landscapes of protein–inhibitor interactions revealed multiple binding poses of drug binding in encounter complexes that could not be resolved by X-ray crystallography^41^.

## Discussion

RTKs activate PI3Kα by recruiting SH2 to the C-terminal phosphorylated pYXXM motifs, which results in the release of p110α autoinhibition^8^. PI3Kα activation involves two critical components. First, nSH2 recruitment, which engenders the destabilization of the iSH2–C2 and ABD–kinase domain interactions. This produces a minor open state with an exposed, favorably oriented membrane-interacting surface. Second, membrane stabilization of this conformation, which results in a membrane-bound, catalysis-ready state. Oncogenic PI3Kα mutants mimic these components, aiming to elevate and strengthen them in the absence of incoming signals to obtain the high-residency, membrane-bound state^42^.

In our studies, we used comprehensive MD simulations to collect activation-tending conformational ensembles of PI3Kα variants with single and double mutations and compiled their conformational profiles (**Fig. 9**). Each p110α mutation projects a spectrum of PI3Kα conformations with low conformational free energy barriers, capable of overcoming the inhibitory state. The hotspot E545K is primarily responsible for nSH2 release. Previous computational studies have also observed spontaneous dissociation of nSH2 by E545K^43^. This helical mutation exposes the pYXXM binding surface of nSH2, facilitating RTK recruitment, shifting the population toward the open state and membrane attachment^44^. nSH2 release also affects iSH2 dynamics and A-loop protrusion. Mutations in the ABD or C2 domain that affect the iSH2 interfaces have a similar effect, but with different strengths. Both R88Q and R93W exert fluctuations in the helix-turn-helix region of iSH2. E453K induces an iSH2 shift from the C2 domain. The conformational effects of R93W and E453K are weak, while those of R88Q are relatively strong. This suggests that R93W and E453K are weak mutations, while R88Q is a moderate mutation. iSH2 dynamics relieve the constraint on the collapsed A-loop conformation, causing it to protrude from the kinase domain surface. Similarly, kinase domain mutations M1043I and H1047R facilitate A-loop protrusion. The hotspot mutation H1047R promotes membrane interaction^11^, as does E726K; however, they do so by different mechanisms and thus with different strengths. The E726K and H1047R mutations replace Ras actions by increasing positive charge at the membrane-interacting surface of the N-lobe and C-lobe kinase domain, respectively, providing anchor points for PI3Kα in the membrane. In our studies, the PI3Kα pose at the lipid bilayer and the membrane-interacting domain are consistent with experimental results^12,39^, particularly the increased membrane binding caused by the reorientation of the WIF motif (W1057, I1058, and F1059) toward the membrane surface, which causes the A-loop to adopt a catalytically competent conformation.

**Fig. 9.**
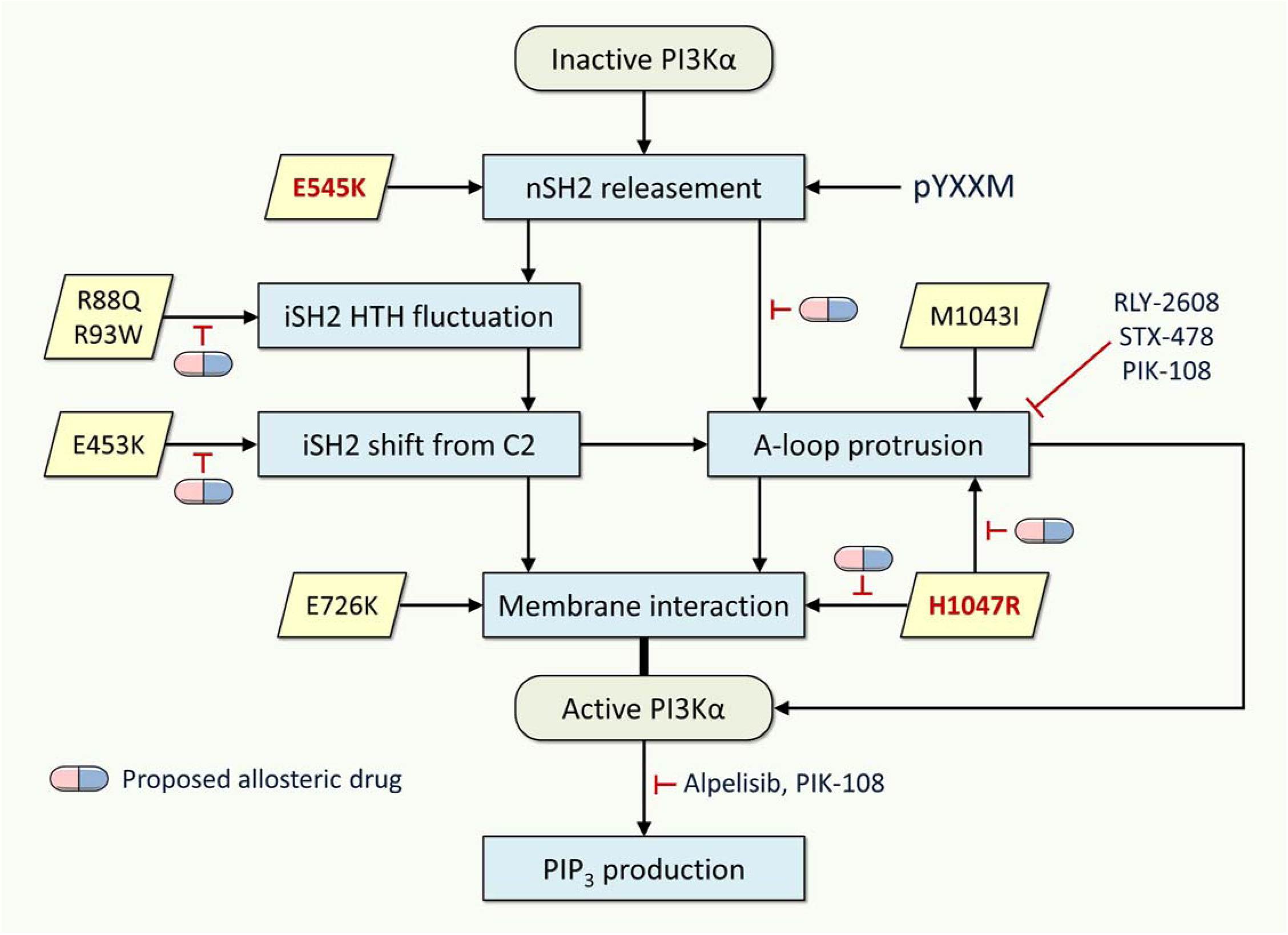
Activation mechanism of PI3K*α* variants and proposed allosteric drugs. The potential roles of PI3Kα mutations, as outlined in this study, provide a valuable framework for understanding the mechanisms involved. Mutations in the helical domain facilitate RTK recruitment by exposing the pYXXM binding surface of SH2, shifting the population toward the open state and membrane attachment. Mutations in the ABD or C2 domain affect the iSH2 dynamics. Mutations in the kinase domain promote membrane interaction. These mechanisms also suggest the design of allosteric drugs targeting PI3Kα variants, which could delay the uncontrolled activation of proteins. Our proposed allosteric drugs can target the following mutations: R88Q and R93W to reduce fluctuations of the helix-turn-helix motif of iSH2; E453K to suppress the shift of iSH2 from the C2 domain; E545K to prevent A-loop protrusion; and H1047R to prevent both A-loop protrusion and membrane interaction. A combination strategy of mutant-specific allosteric drugs is highly efficient at inhibiting the activity of PI3Kα variants with double mutations. The currently available orthosteric drugs that target the active site are alpelisib (Piqray) and PIK-108. Allosteric drugs that are currently available include RLY-2608, STX-478, and PIK-108, which constrain the A-loop allosterically. PIK-108 acts as both an orthosteric and an allosteric drug.

The presence of a second mutation can lead to new structural variants and change the relative stabilities of the ensembles, thereby the population. Double mutations can increase PI3Kα activity synergistically through allosteric communication. Vasan et al.^27^ reported that a double mutation in the *PIK3CA* gene results in increased PI3Kα activity, enhancing PI3Kα signaling and promoting cell proliferation and tumor growth. These mutations in *cis* occur on the same allele, making *PIK3CA* a hyperactive oncogene. This finding has led to clinical trials for cancers with PI3Kα mutations, including breast cancer. The potent E545K hotspot in the helical domain disengages nSH2 from p110α, and combined with this hotspot, the low frequency E453K mutation in the C2 domain concurrently induces iSH2 rotation. Similarly, the R93W mutation in the ABD concurrently induces fluctuations in the helix-turn-helix region of iSH2. These double mutations synergistically promote A-loop protrusion for substrate recruitment. The different locations of the mutations and the different types of residues involved may result in graded mutational outcomes, whether the mutations act through a complementary, e.g., nSH2 disengagement and enhanced membrane interaction, or a similar mechanism. With different single mutations harnessing variant mechanisms, their combinations suggest graded clinical phenotypic spectrum and pharmacological outcomes^12,27^.

Gaining insight into features that differentiate PI3Kα variants can help identify individuals at risk for variant-related diseases, guiding pharmacology. We observe graded strengthening of PI3Kα conformational profiles in solution and in the membrane, which mimic the functional protein environment. The activation of *PIK3CA* by strong or weak driver mutations, combined with other weak or strong latent driver mutations, may result in an increased fraction of active conformations in certain tumors. Suppressing PI3Kα variants harboring a combination of mutations that synergistically increase its activity is thus a coveted aim. Current FDA-approved PI3K inhibitors are primarily orthosteric. They target the ATP-binding site, exerting both orthosteric inhibition and allosteric regulation^45^. However, they often lack selectivity, resulting in off-target effects and undesirable side effects, such as insulin-resistant hyperglycemia^46^. Allosteric drugs, such as the RLY-2608^47,48^, and recently BBO-10203^49^, can overcome these side effects by binding to specific allosteric pockets designed for a particular mutant protein. It is common to combine an allosteric inhibitor with an orthosteric inhibitor^50,51^. For example, combining a mutant-selective allosteric EGFR inhibitor with osimertinib (an ATP-competitive, covalent inhibitor) resulted in increased apoptosis and more effective inhibition of cellular growth in both *in vitro* and *in vivo* models^52^. Similarly, clinical evidence for the combined treatment with two allosteric drugs is emerging. In oncology, targeting SHP2 phosphatase with two allosteric inhibitors at distinct binding sites offers an advantage in developing compounds with favorable pharmacokinetics and cell activity^53^. For central nervous system (CNS) disorders, combination of allosteric modulators has been proposed, and clinical trials targeting GPCR for schizophrenia and other CNS disorders are currently progressing^54,55^. These dual allosteric strategies leverage the advantages of allosteric modulators, such as enhanced selectivity and the capacity to circumvent drug resistance. These strategies enhance therapeutic outcomes compared to single-agent treatments. A two-drug approach targeting the same protein is a drug combination therapy that enhances efficacy, overcomes drug resistance, increases drug residence time^50^, and reduces toxicity. Depending on their mechanisms of action how they interact with the target protein, the two drugs may act synergistically, additively, or even antagonistically. This approach is particularly valuable for treating complex diseases, such as cancer and infectious diseases. In this study, we propose a two-drug approach based on the current drug combination strategy^56^.

The molecular mechanism by which allosteric drugs target PI3Kα mutants with double mutations has been unclear. We discovered cryptic allosteric pockets while observing the dynamics of the variants. These pockets are highly mutation-specific and unstable; thus, they exist transiently and are unobserved in wild-type proteins. A cryptic pocket is a transient allosteric drug binding site that may not be visible in the unbound state of proteins and emerges during protein dynamics. Cryptic pockets are valuable for drug discovery because they can provide novel targets for “undruggable” proteins, such as K-Ras4B with the G12C mutation^57^. The pocket formed beneath the Switch II region is targeted by covalent allosteric inhibitors, such as sotorasib and adagrasib. These FDA-approved allosteric drugs are highly effective against Ras proteins. However, since PI3Kα is much larger than K-Ras4B, a single agent may be ineffective. Thus, for PI3Kα variants, we propose using a combination of orthosteric and allosteric drugs to enhance efficacy, overcoming drug resistance^58^, and improving therapeutic outcomes. The orthosteric drug targets the active site, while the allosteric drug targets the cryptic pocket that emerged due to the mutation, cooperatively enhancing the orthosteric drug action^50^. For double mutations, we suggest considering two mutation-selective allosteric drugs to delay the activation process of the protein. Here we identified tentative cryptic pockets for allosteric drug binding mutational variants. These pockets can facilitate the design of allosteric drugs that precisely target the oncogenic variants. As illustrated in **Fig. 9**, *we suggest that graded conformational patterns of PI3Kα can be targeted by mutation-specific allosteric drugs. For the double mutations, a combination strategy of mutant-specific allosteric drugs could more effectively suppress PI3Kα variants*.

In summary, proteins are not rigid structural snapshots. Large proteins, such as PI3Kα, exhibit multiple conformational ensembles in different environments. PI3Kα variants expand distinct conformational spaces depending on their mutational burdens. PI3Kα is the second most mutated oncogene in cancer, indicating that targeting its variants with a single drug may not be sufficient to suppress its oncogenic function. We found that *each mutation has its own structural features and yields its own conformationally favored drug-binding pocket*. Incorporating a combination of allosteric drugs tailored to specific mutations can be more effective than using a single drug when dealing with variants exhibiting diverse conformational spectra, especially in cases of strong double mutants.

## Materials and methods

### Construction of PI3Kα with mutations

The initial coordinates of the inactive and active forms of wild-type PI3Kα obtained from a previous study^8^ were used to construct the initial configurations for the current simulations of the wild-type and mutant systems. The inactive and active conformations were distinguished by the presence and absence of nSH2, respectively, which can mimic the state of the protein before and after recruitment by RTK. These structures were originally derived from the crystal structure of the inactive form of PI3Kα (PDB ID: 4OVV). The coordinates of ATP and two Mg^2+^, which were missing from the crystal structure, were also adopted from the aforementioned study. We considered seven missense mutations in p110α and six combinations of these mutations (**Supplementary Table 10**). Different PI3Kα mutant systems were generated by modifying the wild-type sequence. The inactive PI3Kα systems, including PI3Kα^R88Q^, PI3Kα^R93W^, PI3Kα^E453K^, PI3Kα^E545K^, PI3Kα^M1043I^, PI3Kα^H1047R^, PI3Kα^R93W/E545K^, PI3Kα^E453K/E545K^, PI3Kα^E545K/M1043I^, PI3Kα^R93W/H1047R^, PI3Kα^E453K/H1047R^, were subjected to simulations in solution. The active PI3Kα systems, including PI3Kα^E726K^, PI3Kα^H1047R^, and PI3Kα^E726K/H1047R^, underwent simulations in a membrane environment. Explicit membrane simulations were performed with the proteins on an anionic lipid bilayer composed of DOPC:DOPS:PIP_2_ (28:6:1 molar ratio). For comparison purposes, both the inactive and active forms of PI3Kα^WT^ were simulated in solution and at the membrane. A total of 16 simulations were performed for different PI3Kα systems in different environments.

### Atomistic molecular dynamics simulations

We performed MD simulations on PI3Kα systems using the updated CHARMM program with the modified all-atom force field (version 36m)^59–61^. Our computational studies closely followed the same protocol as in our previous works^62–77^. The solution simulations were performed using the modified TIP3P water model (three-point model)^78,79^, which constitutes the isometric unit cell containing the protein. To neutralize the system and achieve a total ion concentration of approximately 100 mM, sodium (Na^+^) and chloride (Cl^‒^) ions were added. The cubic box containing PI3Kα and solvent has dimensions of 160 ×160 ×160 Å^3^ and contains nearly 420,000 atoms. The membrane simulations were performed using an anionic lipid bilayer generated by a bilayer-building protocol that involved pseudosphere interactions through a van der Waals (vdW) force field^80,81^. An example of this protocol can be found in the CHARMM program^59^. The lateral dimension of the unit cell is 153.5 ×153.5 Å^2^, constituting the bilayer with a total of 700 lipids (560 DOPC, 120 DOPS and 20 PIP_2_). The anionic bilayer system was formed by randomly distributing DOPC and DOPS across the bilayer plane. Specifically, PIP_2_ was positioned closer to the active site. TIP3P water molecules were added on both sides, with a lipid-to-water ratio of ∼1:200, and Na^+^ and Cl^‒^ were added to generate a final ionic strength of 100 mM and neutralize the system. The final bilayer system contains almost 570,000 atoms.

Following initial construction, both the solution and membrane systems underwent a series of minimizations and dynamics for the solvents, including ions and lipids, with a harmonically restrained protein backbone, until the solvent temperature reached 310 K. Then, preequilibrium simulations involving dynamic cycles were performed as the harmonic restraints on the PI3Kα backbones were gradually released. The final dynamics cycle occurred without the harmonic restraints and completed the preequilibrium stage, generating the starting point for the production run. In the production runs, we employed the particle mesh Ewald (PME) method to calculate long-range electrostatic interactions and vdW interactions to calculate short-range interactions between atoms, using switching functions with the twin range cutoffs at 12 Å and 14 Å. We used the Nosé-Hoover Langevin piston control algorithm to maintain a pressure of 1 atm and the Langevin thermostat method to keep the temperature at 310 K. We applied the SHAKE algorithm to constrain the motion of bonds involving hydrogen atoms. Sixteen different systems were simulated for 1 µs each, and two additional simulations were performed for each system to check reproducibility. For PI3Kα variants in solution, the root-mean-squared-deviation (RMSD) revealed that the domains in p85α slightly deviated during each replica simulation while the domains in p110α mainly retained their initial secondary structures (**Supplementary Fig. 17**). The unstructured interdomain regions in p110α contributed to the deviation in RMSD. Each replica simulation of each PI3Kα variant in membrane yielded similar membrane deviation profiles for the _941_KKKK_944_ motif in the A-loop, ATP, and PIP_2_ (**Supplementary Fig. 18**). However, these deviations differ among the variants. Production runs were performed using the NAMD parallel-computing code^82^ (https://www.ks.uiuc.edu/Research/namd/) on a Biowulf cluster at the National Institutes of Health (https://hpc.nih.gov/). Results were analyzed using the CHARMM program^59^. We used Chimera^83^ (https://www.cgl.ucsf.edu/chimera/) to implement ensemble clustering to obtain conformational representatives and determine the most populated conformation. The PISA program (https://www.ebi.ac.uk/pdbe/pisa/) was used to calculate the surface area of the interface between domains. To identify allosteric signal propagation pathways through the protein, we used the weighted implementation of suboptimal path (WISP)^28^ algorithm (https://github.com/durrantlab/wisp). To discover possible cryptic allosteric pockets that occurred during protein dynamics, we applied the PockDrug program^37^ (https://pockdrug.rpbs.univ-paris-diderot.fr/cgi-bin/index.py?page=Home) to the most populated protein conformation.

### Statistics of PI3Kα double mutations

In our recent work, we aimed to identify latent driver mutations by determining double mutations among pan-cancer datasets ^84^. To this end, we used publicly available TCGA (The Cancer Genome Atlas, https://www.genome.gov/Funded-Programs-Projects/Cancer-Genome-Atlas) and AACR GENIE (Genomics Evidence Neoplasia Information Exchange, https://www.aacr.org/professionals/research/aacr-project-genie/) datasets to check the occurrence of double mutations in several genes mutated in this cohort. We tested the significance of doublets among 62,567 samples after prefiltering with respect to the variant allele frequency (VAF) and the presence of nonsense mutations among double mutant tumors. We labeled constituents of the significant doublets as driver mutation if it is among validated cancer driver mutations deposited in CGI (Cancer Genome Interpreter, https://www.cancergenomeinterpreter.org/home); otherwise, it is a latent driver mutation. We further classify a driver mutation as a strong driver if it is mutated in at least 10% of gene-mutant tumors and as a weak driver otherwise. Similarly, we categorize a latent driver mutation as a strong latent driver if it is mutated in at least 1% of the gene-mutant tumors and as a weak latent driver otherwise. This analysis yielded 155 double mutations accumulated in 53 genes and amongst this cohort, *PIK3CA* harbored 23 double mutations.

## Acknowledgements

This Research was supported by the Cancer Innovation Laboratory, Center for Cancer Research, National Cancer Institute, National Institutes of Health Intramural Research Program project number ZIA BC 010441 and federal funds from the National Cancer Institute, National Institutes of Health, under contract HHSN261201500003I. The contributions of the NIH authors were made as part of their official duties as NIH federal employees, are in compliance with agency policy requirements, and are considered Works of the United States Government. However, the findings and conclusions presented in this paper are those of the authors and do not necessarily reflect the views of the NIH or the U.S. Department of Health and Human Services. All simulations had been performed using the high-performance computational facilities of the Biowulf PC/Linux cluster at the National Institutes of Health, Bethesda, MD (https://hpc.nih.gov/).

## Author contributions

H.J. and M.Z. built models and ran/analyzed molecular dynamics simulations. B.R.Y. collected genomic data. H.J. wrote the initial draft, and B.R.Y., M.Z., Y.L., and R.N. edited the manuscript. R.N. supervised the project.

## Data availability

Representative structures of PI3Kα variants in solution and their starting points in the membrane, along with PDB trajectories, are available in the GitHub repository: https://github.com/hbj-md/PI3K_variants. The repository also contains the CHARMM topology and parameter of PIP_2_ for the standard MD simulations performed in this study.

## References

1. Wu, S., Zhu, W., Thompson, P. & Hannun, Y.A. Evaluating intrinsic and non-intrinsic cancer risk factors. Nat Commun 9, 3490 (2018).

2. Nussinov, R., Tsai, C.J. & Jang, H. A New View of Activating Mutations in Cancer. Cancer Res 82, 4114–4123 (2022).

3. Fruman, D.A. et al. The PI3K Pathway in Human Disease. Cell 170, 605–635 (2017).

4. Madsen, R.R. & Toker, A. PI3K signaling through a biochemical systems lens. J Biol Chem 299, 105224 (2023).

5. Burke, J.E. & Williams, R.L. Synergy in activating class I PI3Ks. Trends Biochem Sci 40, 88–100 (2015).

6. Burke, J.E., Perisic, O., Masson, G.R., Vadas, O. & Williams, R.L. Oncogenic mutations mimic and enhance dynamic events in the natural activation of phosphoinositide 3-kinase p110alpha (PIK3CA). Proc Natl Acad Sci U S A 109, 15259–64 (2012).

7. Miled, N. et al. Mechanism of two classes of cancer mutations in the phosphoinositide 3-kinase catalytic subunit. Science 317, 239–42 (2007).

8. Zhang, M., Jang, H. & Nussinov, R. The mechanism of PI3Kα activation at the atomic level. Chem Sci 10, 3671–3680 (2019).

9. Liu, X. et al. Cryo-EM structures of PI3Kalpha reveal conformational changes during inhibition and activation. Proc Natl Acad Sci U S A 118, e2109327118 (2021).

10. Liu, X. et al. Cryo-EM structures of cancer-specific helical and kinase domain mutations of PI3Kalpha. Proc Natl Acad Sci U S A 119, e2215621119 (2022).

11. Madsen, R.R. et al. Oncogenic PIK3CA corrupts growth factor signaling specificity. Mol Syst Biol 21, 126–157 (2025).

12. Jenkins, M.L. et al. Oncogenic mutations of PIK3CA lead to increased membrane recruitment driven by reorientation of the ABD, p85 and C-terminus. Nat Commun 14, 181 (2023).

13. Chen, Z. et al. Molecular Features of Phosphatase and Tensin Homolog (PTEN) Regulation by C-terminal Phosphorylation. J Biol Chem 291, 14160–14169 (2016).

14. Nussinov, R. & Tsai, C.J. ’Latent drivers’ expand the cancer mutational landscape. Curr Opin Struct Biol 32, 25–32 (2015).

15. Nussinov, R., Jang, H., Tsai, C.J. & Cheng, F. Precision medicine review: rare driver mutations and their biophysical classification. Biophys Rev 11, 5–19 (2019).

16. Li, R., He, X., Wu, C., Li, M. & Zhang, J. Advances in structure-based allosteric drug design. Curr Opin Struct Biol 90, 102974 (2025).

17. Astl, L., Tse, A. & Verkhivker, G.M. Interrogating Regulatory Mechanisms in Signaling Proteins by Allosteric Inhibitors and Activators: A Dynamic View Through the Lens of Residue Interaction Networks. Adv Exp Med Biol 1163, 187–223 (2019).

18. Deng, J., Yuan, Y. & Cui, Q. Modulation of Allostery with Multiple Mechanisms by Hotspot Mutations in TetR. J Am Chem Soc 146, 2757–2768 (2024).

19. Keppler-Noreuil, K.M. et al. Clinical delineation and natural history of the PIK3CA-related overgrowth spectrum. Am J Med Genet A 164A, 1713–33 (2014).

20. Mirzaa, G., et al. PIK3CA-associated developmental disorders exhibit distinct classes of mutations with variable expression and tissue distribution. JCI Insight 1, e87623 (2016).

21. Kuentz, P. et al. Molecular diagnosis of PIK3CA-related overgrowth spectrum (PROS) in 162 patients and recommendations for genetic testing. Genet Med 19, 989–997 (2017).

22. Dogruluk, T. et al. Identification of Variant-Specific Functions of PIK3CA by Rapid Phenotyping of Rare Mutations. Cancer Res 75, 5341–54 (2015).

23. Bukowska, B., Gajek, A. & Marczak, A. Two drugs are better than one. A short history of combined therapy of ovarian cancer. Contemp Oncol (Pozn) 19, 350–3 (2015).

24. Nussinov, R., Yavuz, B.R. & Jang, H. Allostery in Disease: Anticancer Drugs, Pockets, and the Tumor Heterogeneity Challenge. J Mol Biol, 169050 (2025).

25. Campbell, I.G. et al. Mutation of the PIK3CA gene in ovarian and breast cancer. Cancer Res 64, 7678–81 (2004).

26. Tharin, Z. et al. PIK3CA and PIK3R1 tumor mutational landscape in a pan-cancer patient cohort and its association with pathway activation and treatment efficacy. Sci Rep 13, 4467 (2023).

27. Vasan, N. et al. Double PIK3CA mutations in cis increase oncogenicity and sensitivity to PI3Kalpha inhibitors. Science 366, 714–723 (2019).

28. Van Wart, A.T., Durrant, J., Votapka, L. & Amaro, R.E. Weighted Implementation of Suboptimal Paths (WISP): An Optimized Algorithm and Tool for Dynamical Network Analysis. J Chem Theory Comput 10, 511–517 (2014).

29. Zhang, M., Jang, H. & Nussinov, R. Structural Features that Distinguish Inactive and Active PI3K Lipid Kinases. J Mol Biol 432, 5849–5859 (2020).

30. Zhao, L. & Vogt, P.K. Helical domain and kinase domain mutations in p110alpha of phosphatidylinositol 3-kinase induce gain of function by different mechanisms. Proc Natl Acad Sci U S A 105, 2652–7 (2008).

31. Duewell, B.R., Wilson, N.E., Bailey, G.M., Peabody, S.E. & Hansen, S.D. Molecular dissection of PI3Kβ synergistic activation by receptor tyrosine kinases, GβGγ, and Rho-family GTPases. Elife 12, doi: 10.7554/eLife.88991.3 (2024).

32. Buckles, T.C., Ziemba, B.P., Masson, G.R., Williams, R.L. & Falke, J.J. Single-Molecule Study Reveals How Receptor and Ras Synergistically Activate PI3Kalpha and PIP(3) Signaling. Biophys J 113, 2396–2405 (2017).

33. Kotzampasi, D.M., Papadourakis, M., Burke, J.E. & Cournia, Z. Free energy landscape of the PI3Kalpha C-terminal activation. Comput Struct Biotechnol J 23, 3118–3131 (2024).

34. Miller, M.S. et al. Structural basis of nSH2 regulation and lipid binding in PI3Kalpha. Oncotarget 5, 5198–208 (2014).

35. Saito, Y. et al. Landscape and function of multiple mutations within individual oncogenes. Nature 582, 95–99 (2020).

36. Saito, Y., Koya, J. & Kataoka, K. Multiple mutations within individual oncogenes. Cancer Sci 112, 483–489 (2021).

37. Hussein, H.A. et al. PockDrug-Server: a new web server for predicting pocket druggability on holo and apo proteins. Nucleic Acids Res 43, W436–42 (2015).

38. Miller, M.S. et al. Identification of allosteric binding sites for PI3Kalpha oncogenic mutant specific inhibitor design. Bioorg Med Chem 25, 1481–1486 (2017).

39. Hon, W.C., Berndt, A. & Williams, R.L. Regulation of lipid binding underlies the activation mechanism of class IA PI3-kinases. Oncogene 31, 3655–66 (2012).

40. Gkeka, P., Papafotika, A., Christoforidis, S. & Cournia, Z. Exploring a non-ATP pocket for potential allosteric modulation of PI3Kalpha. J Phys Chem B 119, 1002–16 (2015).

41. Re, S., Oshima, H., Kasahara, K., Kamiya, M. & Sugita, Y. Encounter complexes and hidden poses of kinase-inhibitor binding on the free-energy landscape. Proc Natl Acad Sci U S A 116, 18404–18409 (2019).

42. Vanhaesebroeck, B., Perry, M.W.D., Brown, J.R., Andre, F. & Okkenhaug, K. PI3K inhibitors are finally coming of age. Nat Rev Drug Discov 20, 741–769 (2021).

43. Leontiadou, H., Galdadas, I., Athanasiou, C. & Cournia, Z. Insights into the mechanism of the PIK3CA E545K activating mutation using MD simulations. Sci Rep 8, 15544 (2018).

44. Swaney, D.L. et al. A protein network map of head and neck cancer reveals PIK3CA mutant drug sensitivity. Science 374, eabf2911 (2021).

45. Li, M., Rehman, A.U., Liu, Y., Chen, K. & Lu, S. Dual roles of ATP-binding site in protein kinases: Orthosteric inhibition and allosteric regulation. Adv Protein Chem Struct Biol 124, 87–119 (2021).

46. Loke, M., Sehgal, V. & Gupta, N. Alpelisib-Induced Diabetic Ketoacidosis and Insulin-Resistant Hyperglycemia. AACE Clin Case Rep 11, 40–44 (2025).

47. Varkaris, A. et al. Discovery and Clinical Proof-of-Concept of RLY-2608, a First-in-Class Mutant-Selective Allosteric PI3Kalpha Inhibitor That Decouples Antitumor Activity from Hyperinsulinemia. Cancer Discov 14, 240–257 (2024).

48. Gong, G.Q. & Vanhaesebroeck, B. Precision Targeting of Mutant PI3Kalpha. Cancer Discov 14, 204–207 (2024).

49. Simanshu, D.K. et al. BBO-10203 inhibits tumor growth without inducing hyperglycemia by blocking RAS-PI3Kalpha interaction. Science, eadq2004 (2025).

50. Nussinov, R. & Jang, H. How residence time works in allosteric drugs. Curr Opin Struct Biol 94, 103149 (2025).

51. Nussinov, R. & Jang, H. The value of protein allostery in rational anticancer drug design: an update. Expert Opin Drug Discov 19, 1071–1085 (2024).

52. To, C. et al. Single and Dual Targeting of Mutant EGFR with an Allosteric Inhibitor. Cancer Discov 9, 926–943 (2019).

53. Sarver, P. et al. 6-Amino-3-methylpyrimidinones as Potent, Selective, and Orally Efficacious SHP2 Inhibitors. J Med Chem 62, 1793–1802 (2019).

54. Wild, C., Cunningham, K.A. & Zhou, J. Allosteric Modulation of G Protein-Coupled Receptors: An Emerging Approach of Drug Discovery. Austin J Pharmacol Ther 2(2014).

55. Conn, P.J., Kuduk, S.D. & Doller, D. Drug Design Strategies for GPCR Allosteric Modulators. Annu Rep Med Chem 47, 441–457 (2012).

56. Nussinov, R., Yavuz, B.R. & Jang, H. Anticancer drugs: How to select small molecule combinations? Trends Pharmacol Sci 45, 503–519 (2024).

57. Ostrem, J.M., Peters, U., Sos, M.L., Wells, J.A. & Shokat, K.M. K-Ras(G12C) inhibitors allosterically control GTP affinity and effector interactions. Nature 503, 548–51 (2013).

58. Gremke, N. et al. Targeting PI3K inhibitor resistance in breast cancer with metabolic drugs. Signal Transduct Target Ther 10, 92 (2025).

59. Brooks, B.R. et al. CHARMM: the biomolecular simulation program. J Comput Chem 30, 1545–614 (2009).

60. Klauda, J.B. et al. Update of the CHARMM all-atom additive force field for lipids: validation on six lipid types. J Phys Chem B 114, 7830–43 (2010).

61. Huang, J. et al. CHARMM36m: an improved force field for folded and intrinsically disordered proteins. Nat Methods 14, 71–73 (2017).

62. Liu, Y., Zhang, W., Jang, H. & Nussinov, R. mTOR Variants Activation Discovers PI3K-like Cryptic Pocket, Expanding Allosteric, Mutant-Selective Inhibitor Designs. J Chem Inf Model 65, 966–980 (2025).

63. Xu, L., Jang, H. & Nussinov, R. Allosteric modulation of NF1 GAP: Differential distributions of catalytically competent populations in loss-of-function and gain-of-function mutants. Protein Sci 34, e70042 (2025).

64. Xu, L., Jang, H. & Nussinov, R. Capturing Autoinhibited PDK1 Reveals the Linker’s Regulatory Role, Informing Innovative Inhibitor Design. J Chem Inf Model 64, 7709–7724 (2024).

65. Zhang, W., Liu, Y., Jang, H. & Nussinov, R. Slower CDK4 and faster CDK2 activation in the cell cycle. Structure 32, 1269–1280 e2 (2024).

66. Zhang, W., Liu, Y., Jang, H. & Nussinov, R. CDK2 and CDK4: Cell Cycle Functions Evolve Distinct, Catalysis-Competent Conformations, Offering Drug Targets. JACS Au 4, 1911–1927 (2024).

67. Liu, Y., Zhang, M., Jang, H. & Nussinov, R. The allosteric mechanism of mTOR activation can inform bitopic inhibitor optimization. Chem Sci 15, 1003–1017 (2024).

68. Liu, Y., Zhang, W., Jang, H. & Nussinov, R. SHP2 clinical phenotype, cancer, or RASopathies, can be predicted by mutant conformational propensities. Cell Mol Life Sci 81, 5 (2024).

69. Jang, H., Chen, J., Iakoucheva, L.M. & Nussinov, R. Cancer and Autism: How PTEN Mutations Degrade Function at the Membrane and Isoform Expression in the Human Brain. J Mol Biol 435, 168354 (2023).

70. Jang, H., Smith, I.N., Eng, C. & Nussinov, R. The mechanism of full activation of tumor suppressor PTEN at the phosphoinositide-enriched membrane. iScience 24, 102438 (2021).

71. Zhang, M., Maloney, R., Jang, H. & Nussinov, R. The mechanism of Raf activation through dimerization. Chem Sci 12, 15609–15619 (2021).

72. Zhang, M., Jang, H., Li, Z., Sacks, D.B. & Nussinov, R. B-Raf autoinhibition in the presence and absence of 14-3-3. Structure 29, 768–777 e2 (2021).

73. Maloney, R.C., Zhang, M., Jang, H. & Nussinov, R. The mechanism of activation of monomeric B-Raf V600E. Comput Struct Biotechnol J 19, 3349–3363 (2021).

74. Jang, H., Zhang, M. & Nussinov, R. The quaternary assembly of KRas4B with Raf-1 at the membrane. Comput Struct Biotechnol J 18, 737–748 (2020).

75. Zhang, M., Jang, H. & Nussinov, R. The structural basis for Ras activation of PI3Kalpha lipid kinase. Phys Chem Chem Phys 21, 12021–12028 (2019).

76. Jang, H., Muratcioglu, S., Gursoy, A., Keskin, O. & Nussinov, R. Membrane-associated Ras dimers are isoform-specific: K-Ras dimers differ from H-Ras dimers. Biochem J 473, 1719–32 (2016).

77. Jang, H. et al. The higher level of complexity of K-Ras4B activation at the membrane. FASEB J 30, 1643–55 (2016).

78. Durell, S.R., Brooks, B.R. & Bennaim, A. Solvent-Induced Forces between 2 Hydrophilic Groups. Journal of Physical Chemistry 98, 2198–2202 (1994).

79. Price, D.J. & Brooks, C.L., 3rd. A modified TIP3P water potential for simulation with Ewald summation. J Chem Phys 121, 10096–103 (2004).

80. Woolf, T.B. & Roux, B. Molecular dynamics simulation of the gramicidin channel in a phospholipid bilayer. Proc Natl Acad Sci U S A 91, 11631–5 (1994).

81. Woolf, T.B. & Roux, B. Structure, energetics, and dynamics of lipid-protein interactions: A molecular dynamics study of the gramicidin A channel in a DMPC bilayer. Proteins 24, 92–114 (1996).

82. Phillips, J.C. et al. Scalable molecular dynamics with NAMD. J Comput Chem 26, 1781–802 (2005).

83. Pettersen, E.F. et al. UCSF Chimera--a visualization system for exploratory research and analysis. J Comput Chem 25, 1605–12 (2004).

84. Yavuz, B.R., Tsai, C.J., Nussinov, R. & Tuncbag, N. Pan-cancer clinical impact of latent drivers from double mutations. Commun Biol 6, 202 (2023).

