## Supplemental figures and tables for "Oncogenic PI3Kα variants reveal graded conformational spectrum with mutation-specific cryptic pockets"

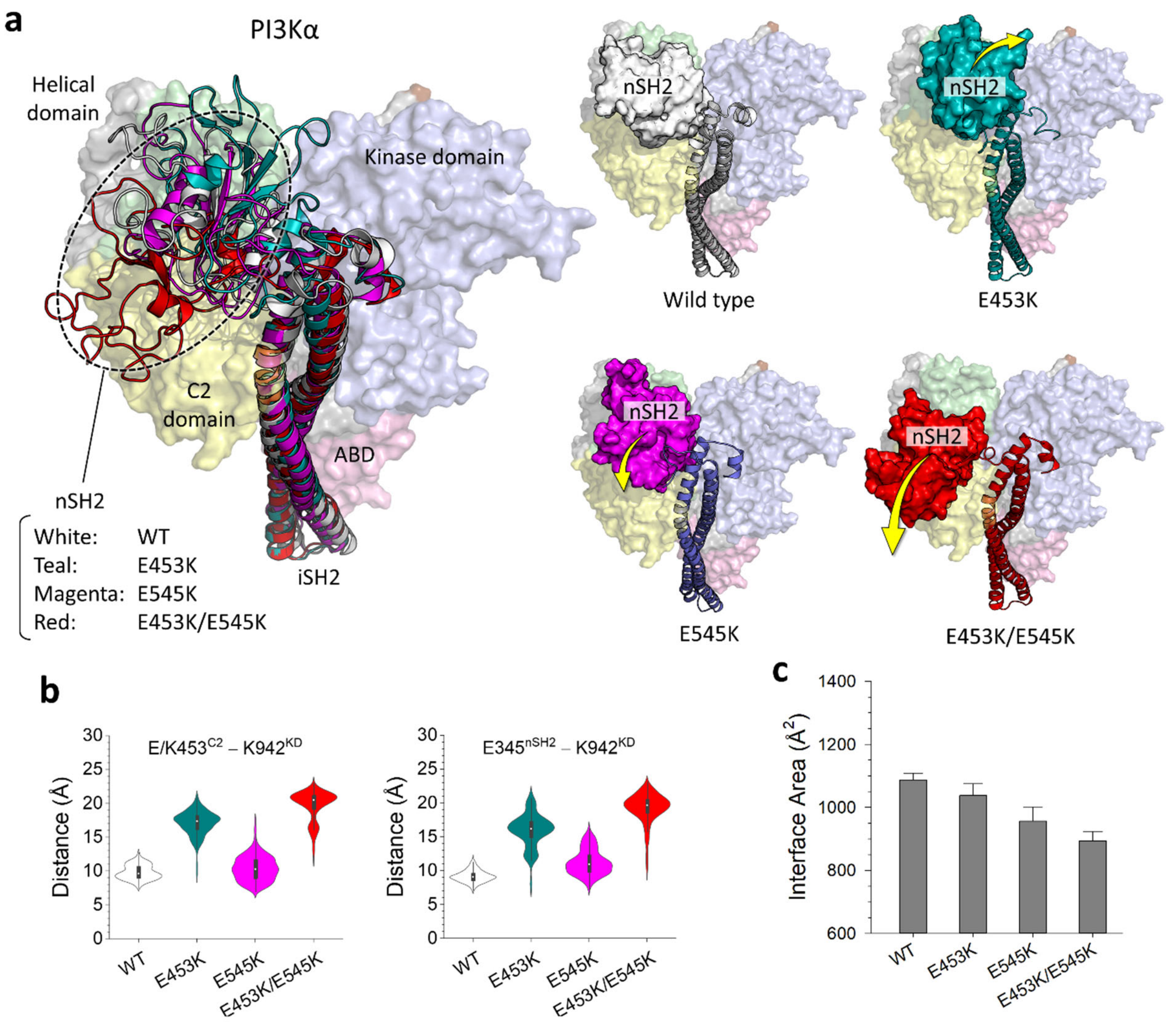


**Supplementary Fig. 1 | The best representative conformations of PI3Kα variants compared with the wild type.**

**a**, Superimposition of the best representative conformations of PI3Kα$^{WT}$, PI3Kα$^{E453K}$, PI3Kα$^{E545K}$, and PI3Kα$^{E453K/E545K}$ (*left*), along with their individual snapshots (*right*). Yellow arrows indicate the nSH2 movement relative to the wild type. The size of arrows refers to the strength of the movement. **b**, Violin plots representing the atomic pair distances of E/K453$^{C2}$–K942$^{KD}$ and E345$^{nSH2}$–K942$^{KD}$. **c**, The surface area of the interface between nSH2 and the p110α subunit calculated using the PISA program (https://www.ebi.ac.uk/pdbe/pisa/).

.

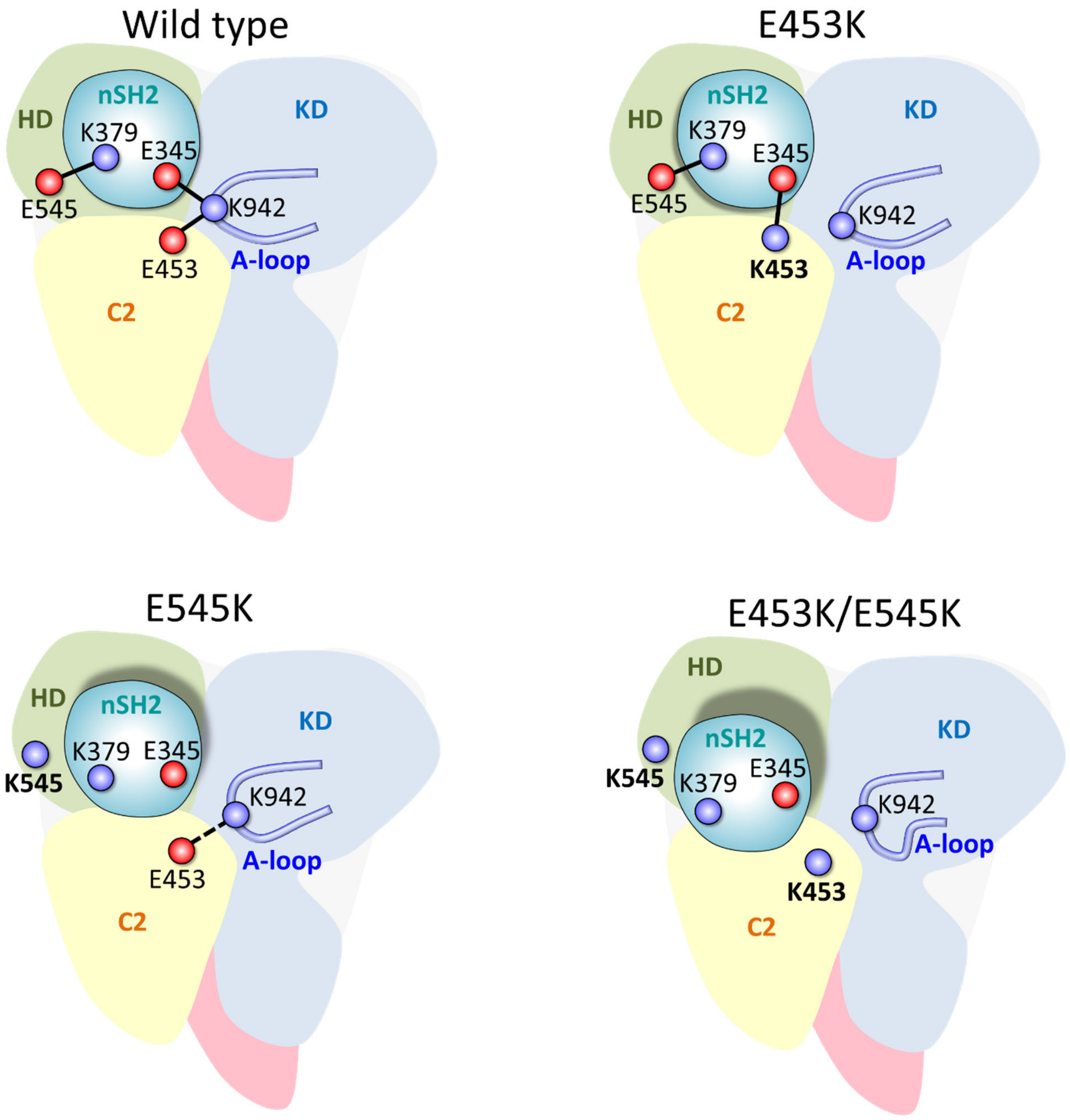


**Supplementary Fig. 2 | The nSH2 release mechanism.**
Schematic diagrams represent how the mutations in PI3Kα disrupt the salt bridge network at the nSH2/helical/C2/kinase domain interface. A significant conformational change in the A-loop can be observed upon nSH2 release.

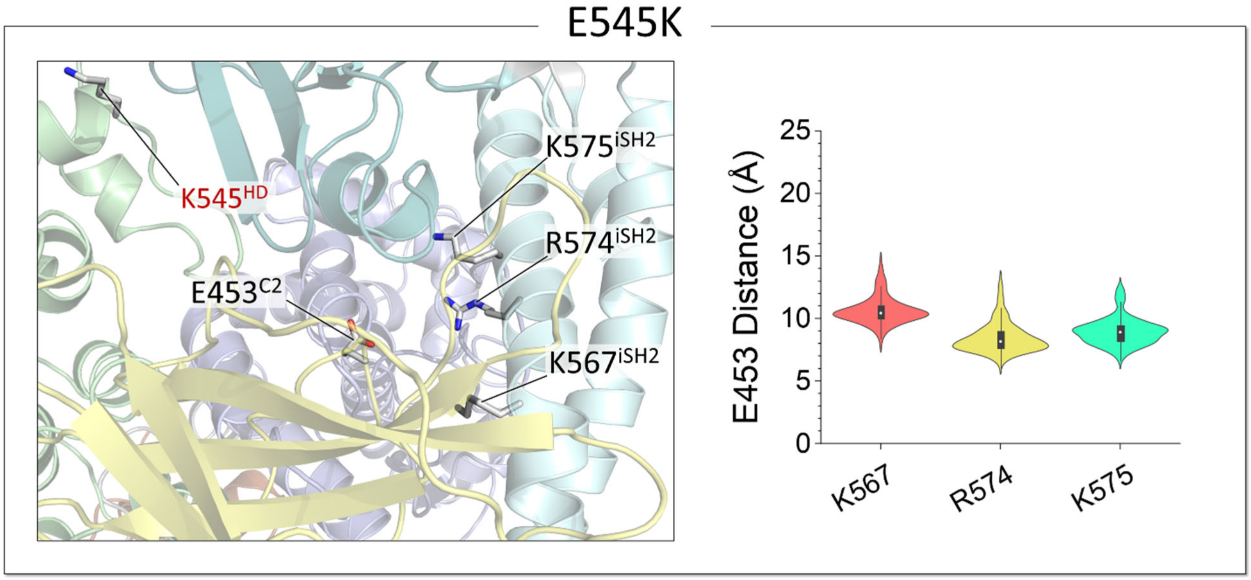


**Supplementary Fig. 3 | The hotspot E545K mutation alone is less effective on the iSH2 movement.**
Highlights of the salt bridge interactions at the iSH2/C2 domain interfaces for PI3Kα$^{E545K}$ (*left panel*), and violin plots representing the atomic pair distances of E453$^{C2}$–K567$^{iSH2}$, E453$^{C2}$–R574$^{iSH2}$, and E453$^{C2}$– K575$^{iSH2}$ (*right panel*). The protein structure is depicted as a cartoon with different colors: the C2 domain is yellow, the helix domain is green, the kinase domain is blue, and iSH2 is cyan.

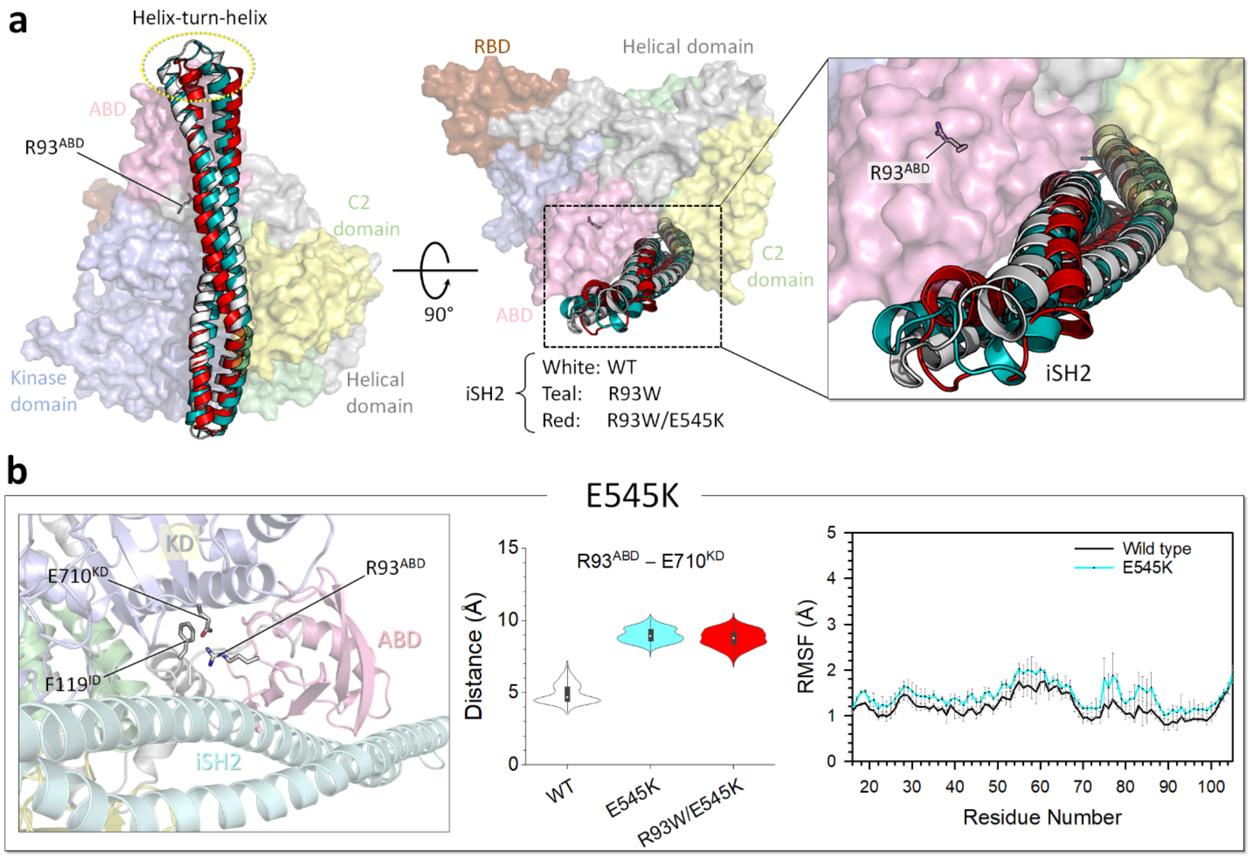


**Supplementary Fig. 4 | The R93W mutation disrupts the ABD/kinase domain interface.**
**a,** Superimposition of the best representative conformations of PI3Kα$^{WT}$, PI3Kα$^{R93W}$, and PI3Kα$^{R93W/E545K}$ highlighting the movement of the helix-turn-helix region of iSH2. The p110α is shown in surface representation with different color codes: The ABD is pink, the RBD is brown, the C2 domain is yellow, the helical domain is green, and the kinase domain is blue. The iSH2 structure is depicted as a cartoon, showing the wild type in white, the single mutation R93W in teal, and the double mutation R93W/E545K in red. **b**, The best representative conformation from the ensemble clusters for PI3Kα$^{E545K}$ is shown, with highlights of the residue interactions at the ABD/kinase domain interfaces (*left*). Violin plot representing the atomic pair distance of R93$^{ABD}$–E710$^{KD}$ compared with the wild type and the double mutation R93W/E545K (*middle*). The root-mean-squared-fluctuation (RMSF) of the ABD for PI3Kα$^{E545K}$ (cyan) compared with the wild type (black) (*right*).

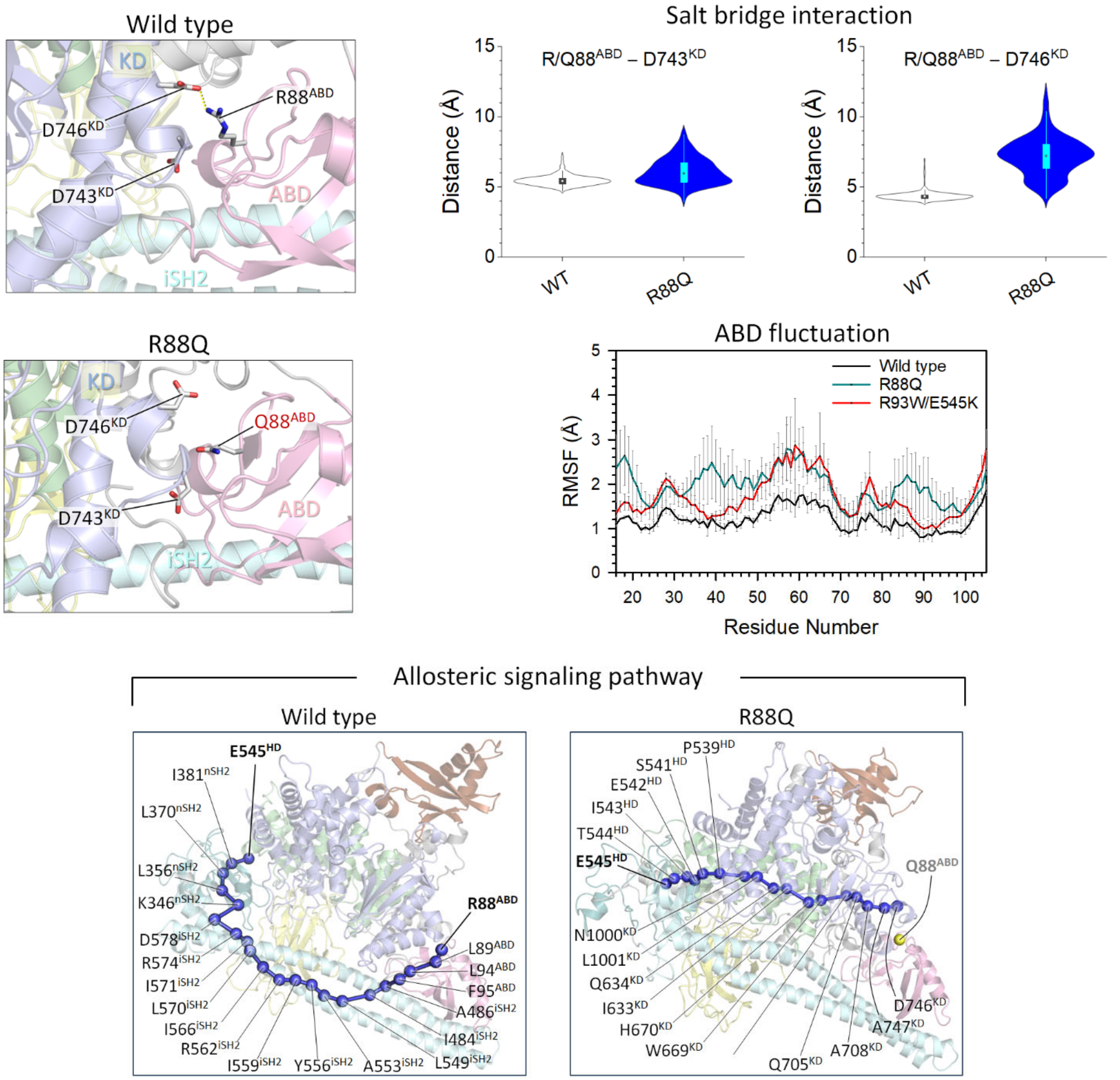


**Supplementary Fig. 5 | The R88Q mutation also disrupts the ABD/kinase domain interface.** The best representative conformations from the ensemble clusters for PI3Kα$^{WT}$ and PI3Kα$^{R88Q}$ are shown, with highlights of the residue interactions at the ABD/kinase domain interfaces (*left*). The protein structure is depicted as a cartoon with different colors: the ABD is pink, the helix domain is green, the kinase domain is blue, and iSH2 is cyan. Violin plots representing the atomic pair distances of R/Q88$^{ABD}$–D743$^{KD}$ and R/Q88$^{ABD}$–D746$^{KD}$ (*right*). The root-mean-squared-fluctuation (RMSF) of the ABD for PI3Kα$^{R88Q}$ (teal) compared with that for PI3Kα$^{WT}$ (black) and PI3Kα$^{R93W/E545K}$ (red) (*middle*). The allosteric signaling pathways between the source residue E545$^{HD}$ and the sink residue R/Q88$^{ABD}$. Blue beads denote the allosteric signal nodes, and yellow bead for Q88$^{ABD}$ in the R88Q mutation indicates the broken allosteric signal.

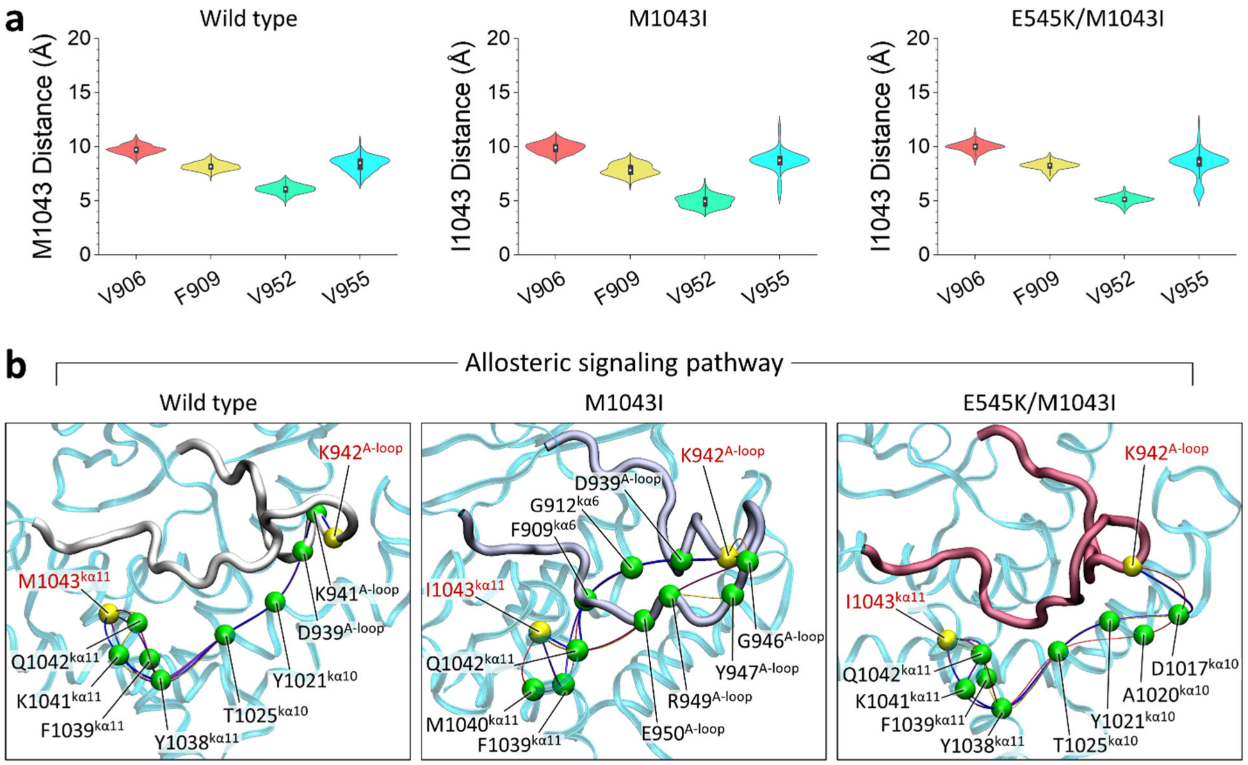


**Supplementary Fig. 6 | The M1043I mutation affects the A-loop conformation.**
**a**, Violin plots representing the atomic pair distances of M/I1043$^{k\alpha 11}$–V906$^{k\alpha 6}$, M/I1043$^{k\alpha 11}$–F909$^{k\alpha 6}$, M/I1043$^{k\alpha 11}$–V952$^{A\text{-}loop}$, and M/I1043$^{k\alpha 11}$–V955$^{A\text{-}loop}$ for the wild type, the single mutation M1043I, and the double mutation E545K/M1043I. Both mutations preserve the hydrophobic cluster, similar to the wild type. **b**, The allosteric signaling pathways between the source residue M/I1043$^{k\alpha 11}$ and the sink residue K942$^{A\text{-}loop}$. Yellow beads represent the source and sink residues, while green beads denote the allosteric signal nodes.

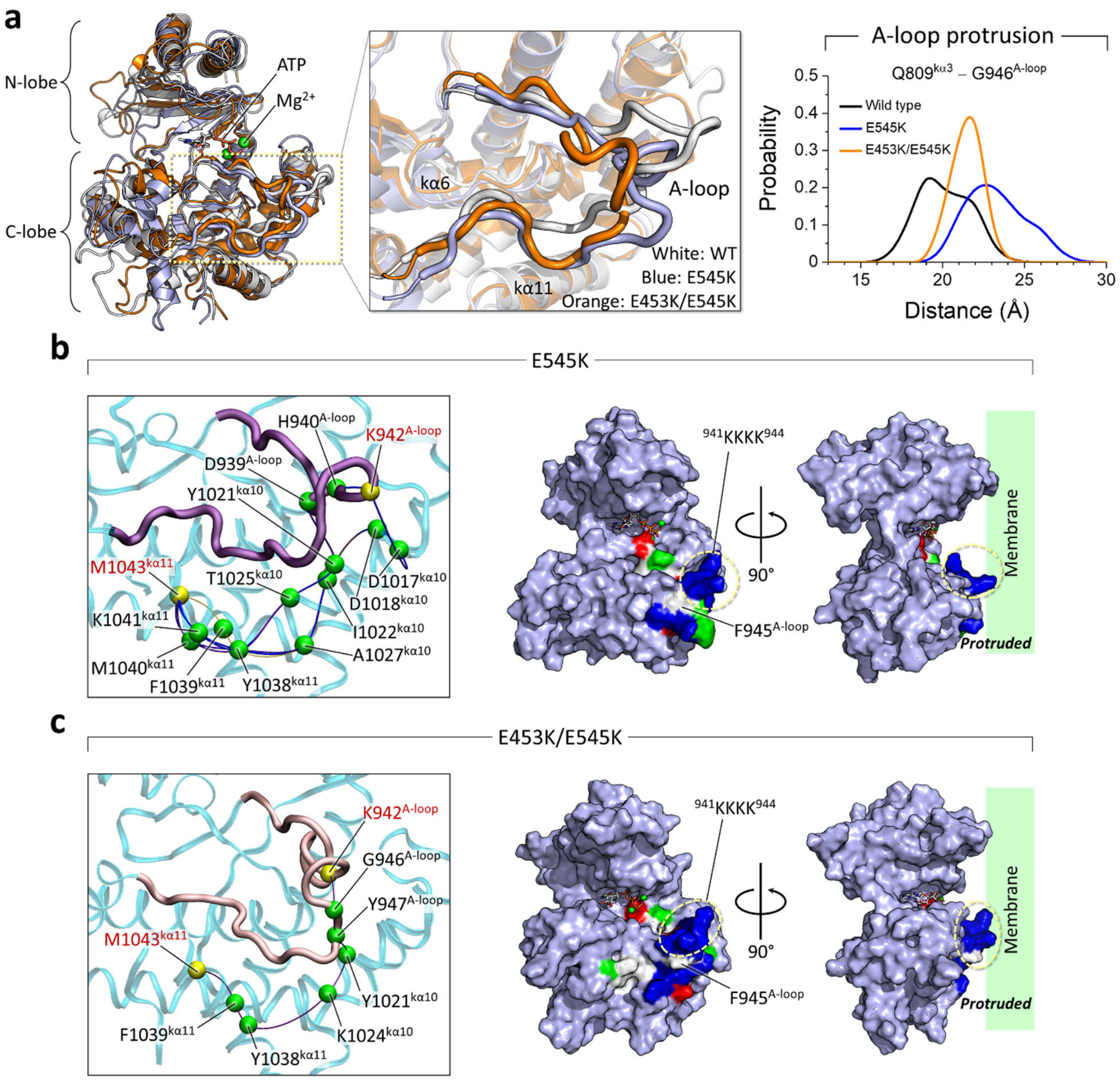


**Supplementary Fig. 7 | Both E545K and E453K/E545K mutations also cause the A-loop to protrude.**

**a**, Superimposition of kinase domains from the best representative conformations highlights A-loop conformations. Wild-type is shown in white, the single mutation E545K is shown in blue, and the double mutation E453K/E545K is shown in orange (*left and middle*). The probability distribution of the distance between $Q809^{k\alpha3}$ and $G946^{\text{A-loop}}$ representing the A-loop protrusion (*right*). **b**, The allosteric signaling pathway between the source residue $M1043^{k\alpha11}$ and the sink residue $K942^{\text{A-loop}}$, and the kinase domain structure highlighting the A-loop for $PI3K\alpha^{E545K}$. **c**, The same for $PI3K\alpha^{E453K/E545K}$. In the pathways, yellow beads represent the source and sink residues, while green beads denote the allosteric signal nodes. In the A-loop structure, residues are colored white, green, blue, and red to indicate their hydrophobic, polar, positively charged, and negatively charged properties, respectively. The protruding region of the A-loop toward the pseudo-membrane is marked.

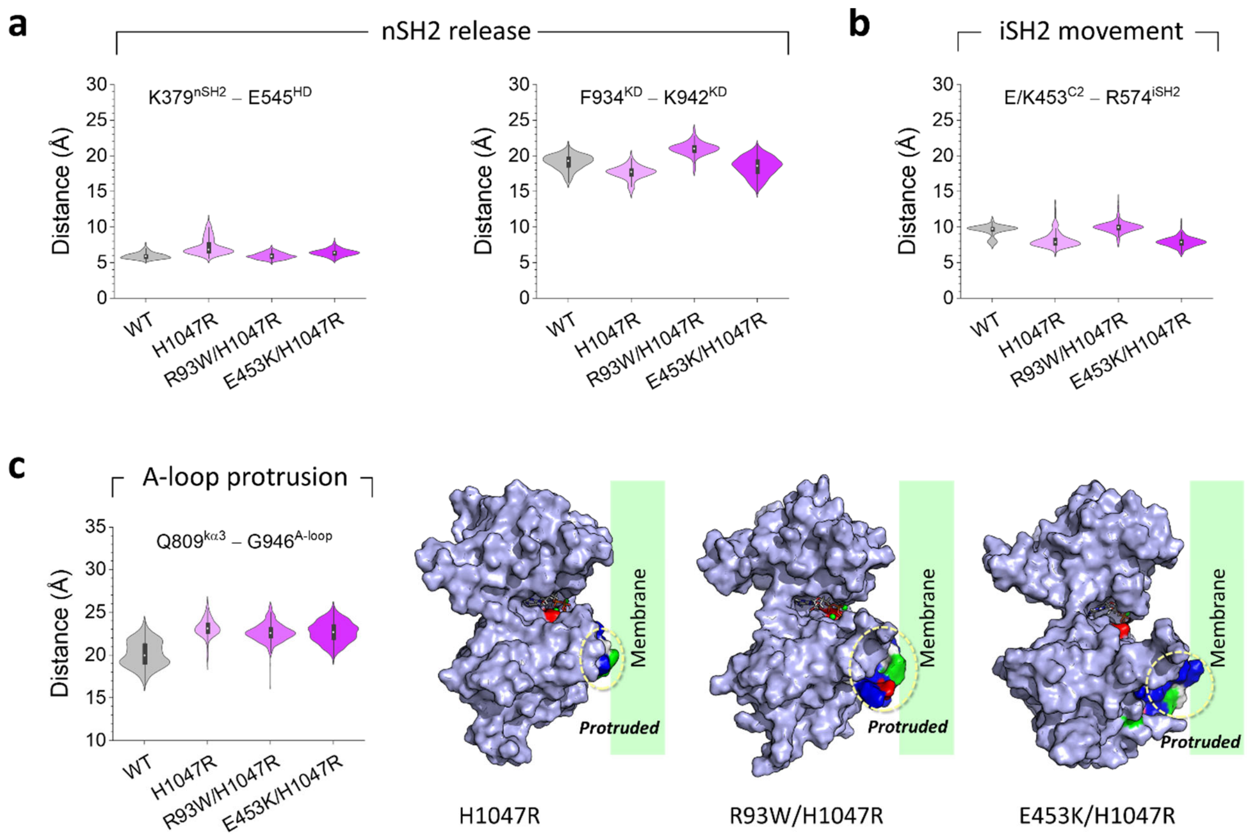


**Supplementary Fig. 8 | The H1047R mutation slightly affects the A-loop dynamics.**
**a**, Violin plots representing the atomic pair distances of K379$^{nSH2}$–E545$^{HD}$ and F934$^{KD}$–K942$^{KD}$ for PI3Kα$^{WT}$, PI3Kα$^{H1047R}$, PI3Kα$^{R93W/H1047R}$, and PI3Kα$^{E453K/H1047R}$, indicating the nSH2 release. HD and KD denote the helical and kinase domains, respectively. **b**, Violin plot representing the atomic pair distance of E/K453$^{C2}$–R574$^{iSH2}$, indicating the iSH2 movement. **c**, Violin plot representing the atomic pair distance of Q809$^{k\alpha3}$–G946$^{A\text{-}loop}$, indicating the A-loop protrusion, and the kinase domain structures highlighting the A-loop for the mutations. The kinase domain is colored cyan. In the A-loop structure, residues are colored white, green, blue, and red to indicate their hydrophobic, polar, positively charged, and negatively charged properties, respectively. The protruding region of the A-loop toward the pseudo-membrane is marked.

.

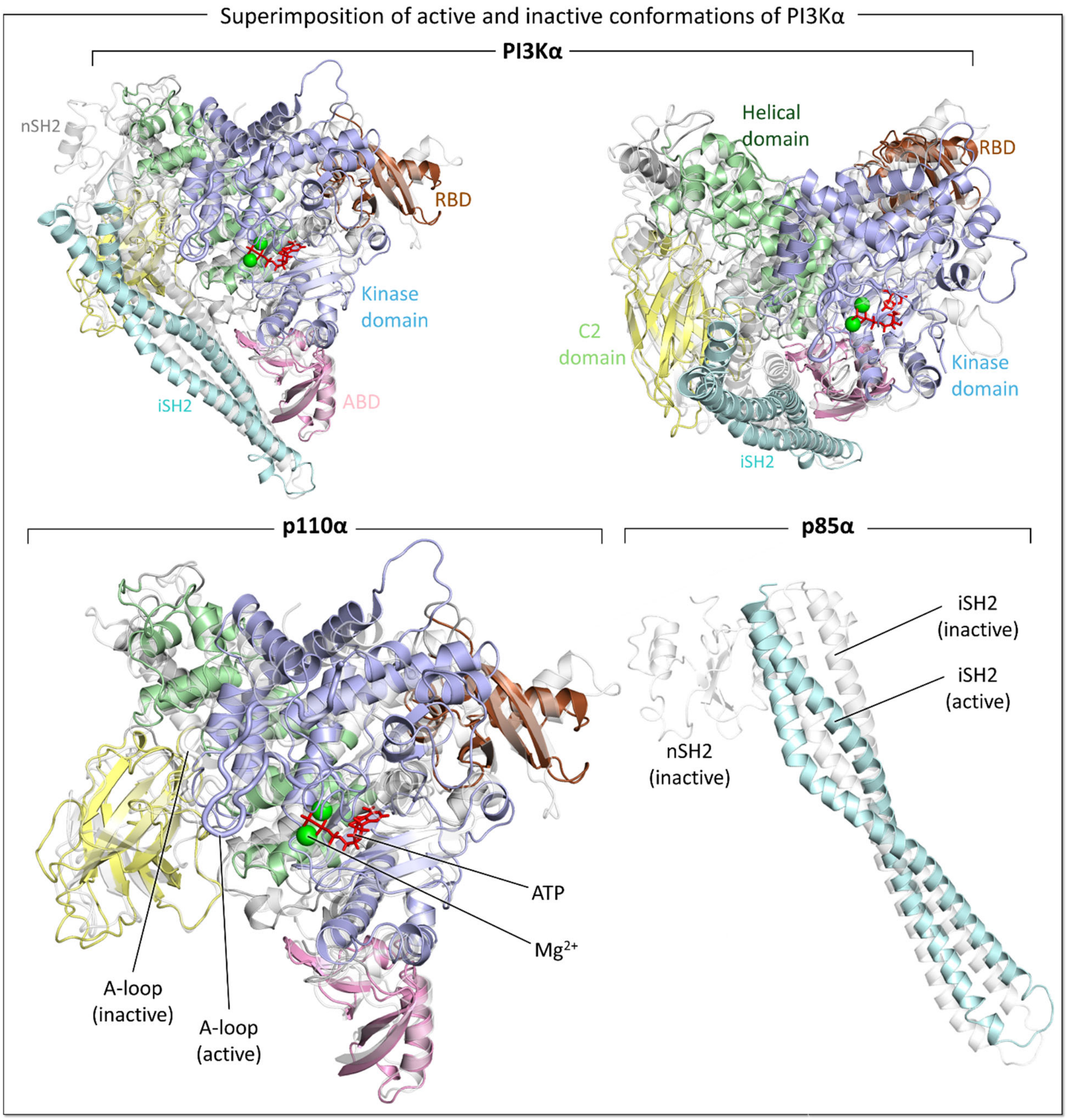


**Supplementary Fig. 9 | Active and inactive conformations of PI3Kα differ.**
Superimposition of the active and inactive conformations of $PI3K\alpha^{WT}$. Blow, the superimposition of the p110α and p85α subunits is highlighted separately. The colored cartoon represents the active form, and the while white, transparent cartoon represents the inactive form.

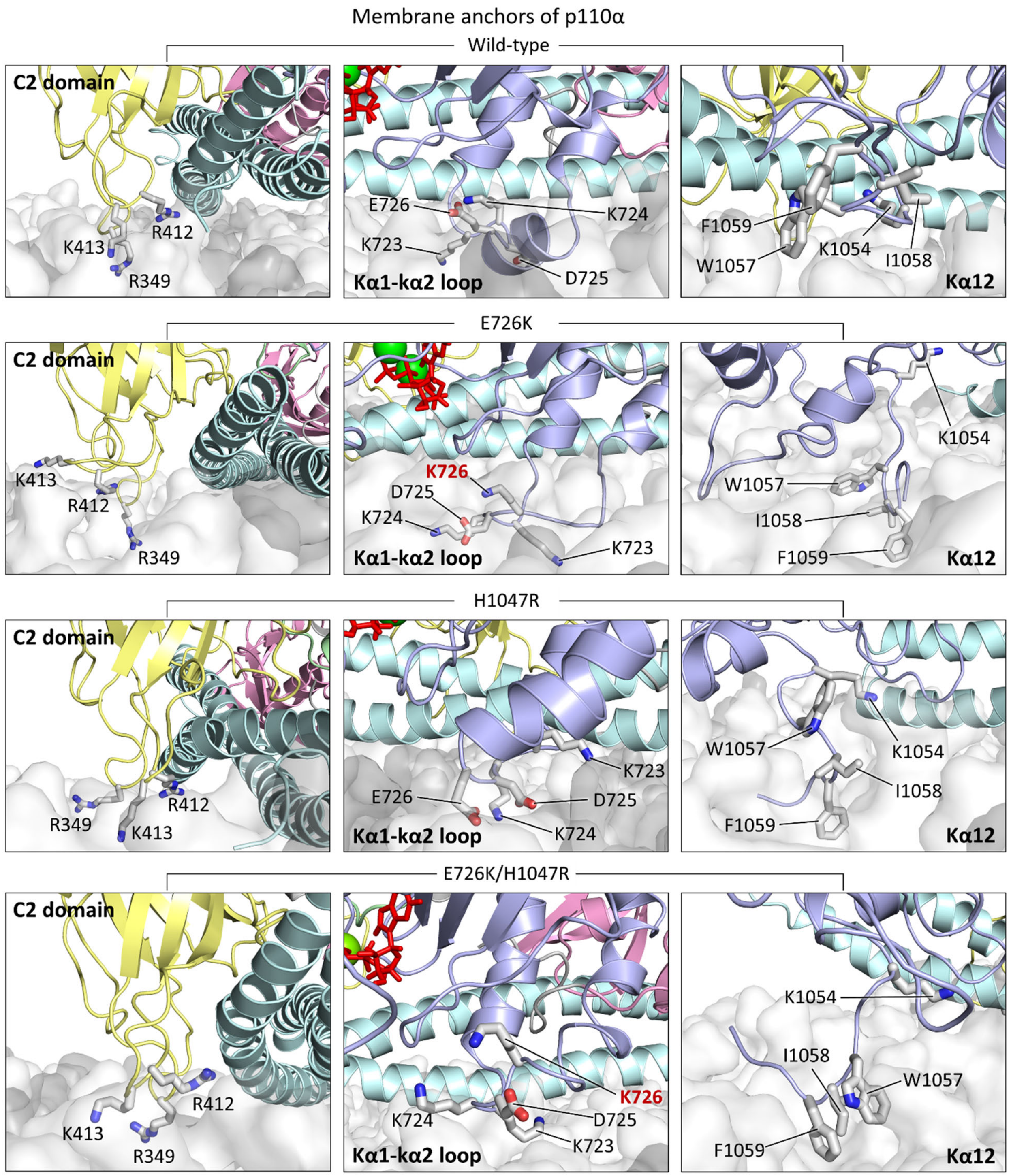


**Supplementary Fig. 10 | Membrane interactions of the three anchor points for active PI3Kα.** The best representative conformations from the ensemble clusters for PI3Kα$^{WT}$, PI3Kα$^{E726K}$, PI3Kα$^{H1047R}$, and PI3Kα$^{E726K/H1047R}$ are shown, with highlights of the residues interacting with the membrane at the three anchor points located in the C2 domain, the kα1-kα2 loop, and the kα12 helix.

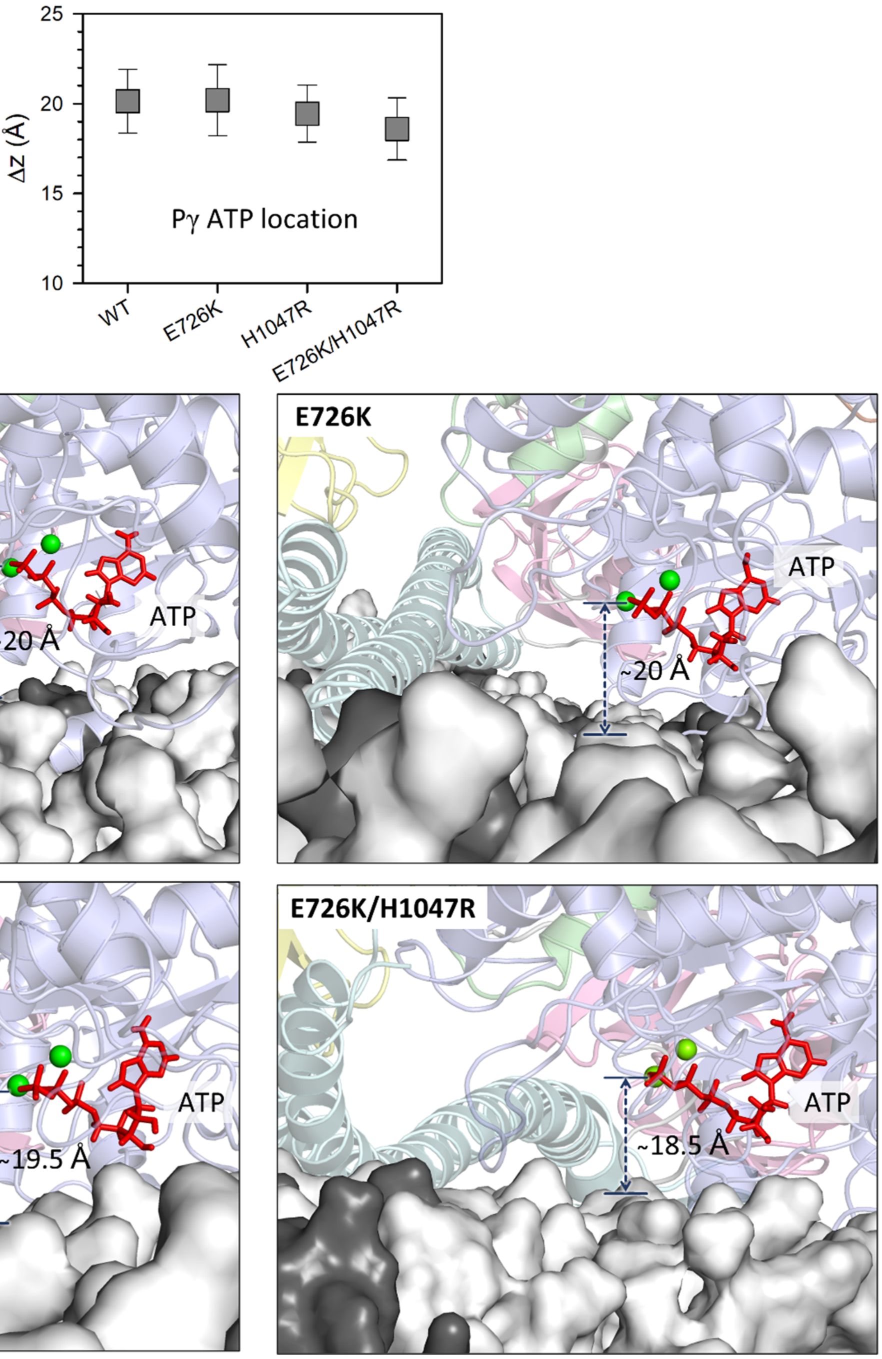


**Supplementary Fig. 11 | ATP location from the bilayer surface.**
Deviation of Pγ of ATP from the averaged position of the phosphate atoms of DOPC and DOPS for $PI3K\alpha^{WT}$, $PI3K\alpha^{E726K}$, $PI3K\alpha^{H1047R}$, and $PI3K\alpha^{E726K/H1047R}$. Snapshots highlighting ATP with deviations of Pγ of ATP marked on each system.

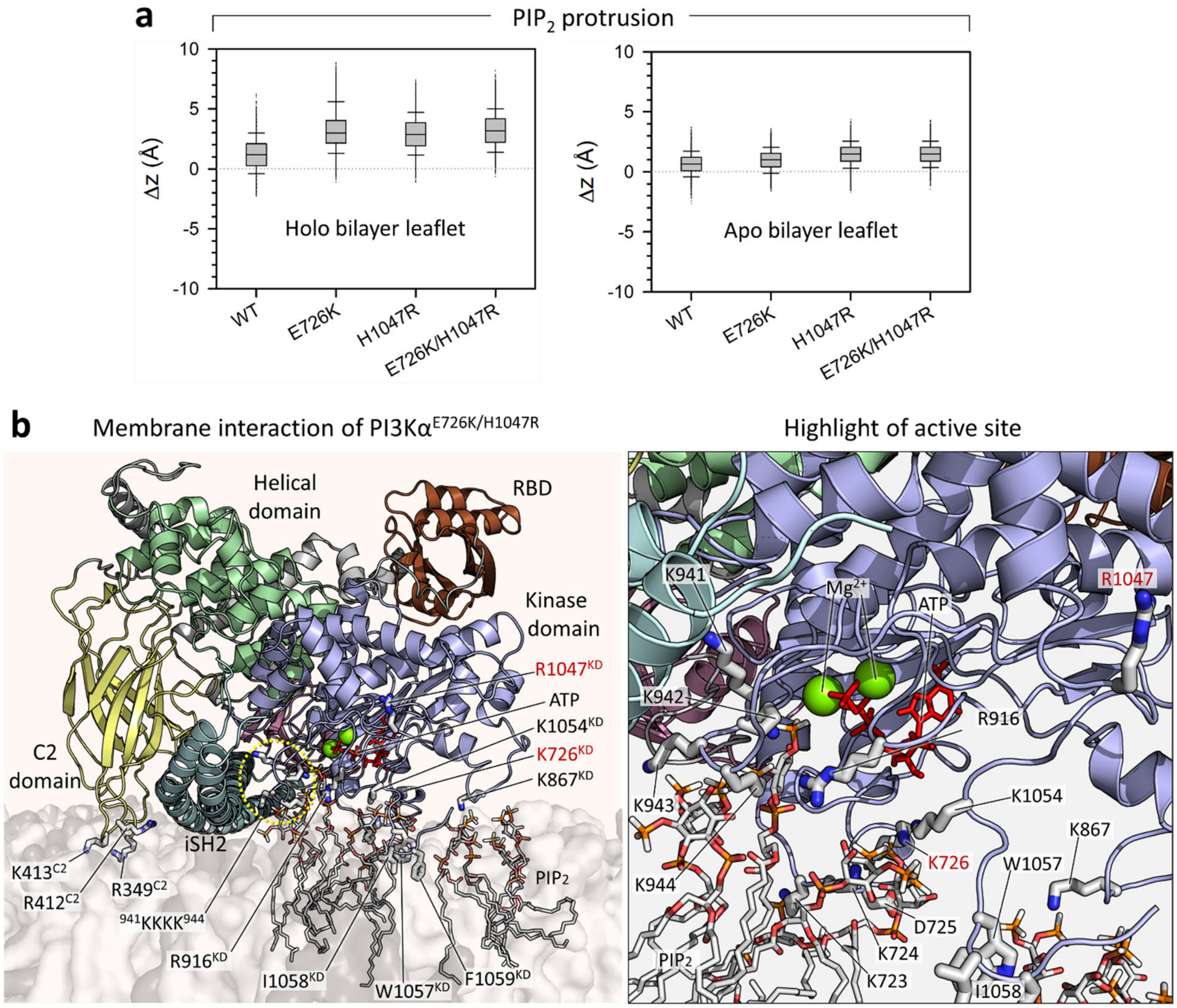


**Supplementary Fig. 12 | PIP₂ protrusion.**

**a**, Deviations of the phosphate atoms of $PIP_2$ from the averaged position of the phosphate atoms of DOPC and DOPS for the *holo* and *apo* bilayer leaflets for $PI3K\alpha^{WT}$, $PI3K\alpha^{E726K}$, $PI3K\alpha^{H1047R}$, and $PI3K\alpha^{E726K/H1047R}$. **b**, Snapshots showing membrane interaction of $PI3K\alpha^{E726K/H1047R}$ and highlighting the active site in the kinase domain. $PIP_2$ is protruding from the bilayer surface toward the active site. The mutant residues K726 and R1047 are marked in red. The protein structure is depicted as a cartoon with different color codes: The ABD is pink, the RBD is brown, the C2 domain is yellow, the helical domain is green, the kinase domain is blue, and iSH2 is teal. $PIP_2$ is shown as sticks. The white and gray transparent surfaces represent DOPC and DOPS, respectively. ATP is shown as red sticks, and $Mg^{2+}$ is shown as a green sphere.

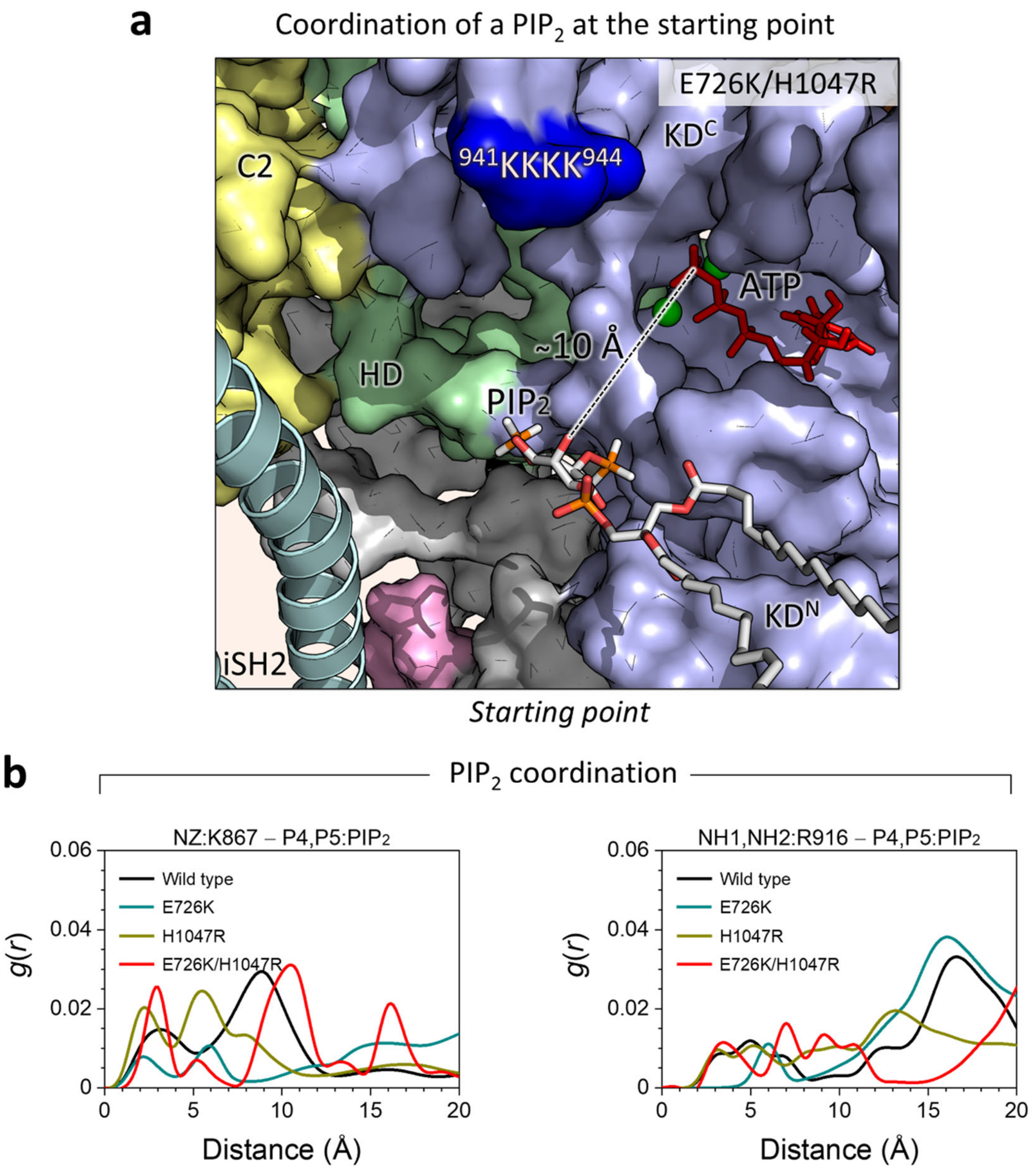


**Supplementary Fig. 13 | $PIP_2$ coordination.**

**a**, Snapshot highlighting the active site of PI3Kα$^{E726K/H1047R}$ at the starting point. A particular $PIP_2$ is initially placed at the active site with a coordination distance of ~10 Å. The coordination distance is defined as the distance between the hydroxyl group at the 3rd position of the inositol ring of $PIP_2$ and the Pγ of ATP. The p110α structure is shown in surface representation with different color codes: The C2 domain is yellow, the helical domain is green, and the kinase domain is blue. The p85α structure is depicted as a cartoon. $PIP_2$ is shown as sticks. ATP is shown as red sticks, and $Mg^{2+}$ is shown as a green sphere. C2 and HD denote the C2 and helical domains, respectively. $KD^N$ and $KD^C$ represent the N-lobe and C-lobe kinase domain, respectively. The dark blue surface highlights the $^{941}$KKKK$^{944}$ motif in the A-loop. **b**, The radial distribution function, $g(r)$, of selected atom pairs for PI3Kα$^{WT}$, PI3Kα$^{E726K}$, PI3Kα$^{H1047R}$, and PI3Kα$^{E726K/H1047R}$.

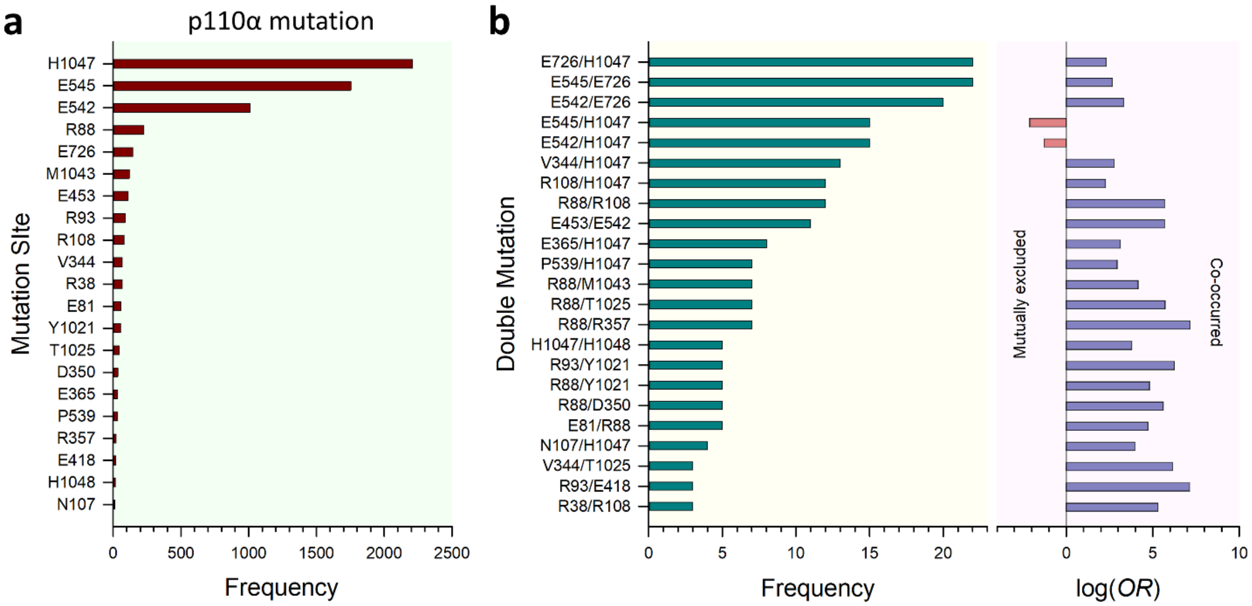


**Supplementary Fig. 14 | *PIK3CA* mutations and their frequencies.**

**a**, Statistical analyses based on the frequencies of single mutations in p110α. **b**, Statistical analyses based on the frequencies of double mutations. The red bars on the left indicate mutually exclusive doublets for E545/H1047 and E542/H1047.

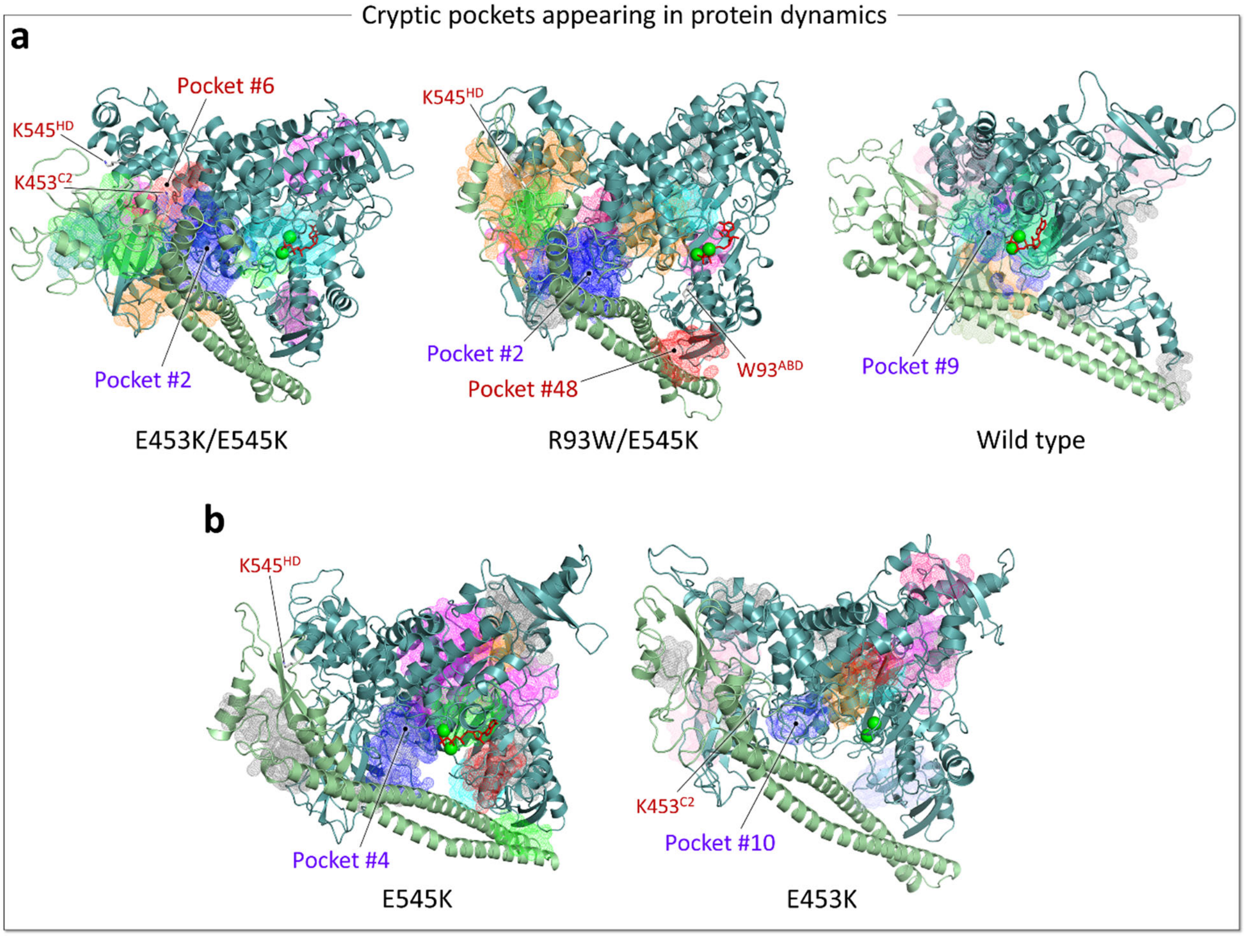


**Supplementary Fig. 15 | Discovery of cryptic allosteric pockets in PI3Kα**

**a**, Predicted cryptic allosteric pockets for PI3Kα$^{E453K/E545K}$, PI3Kα$^{R93W/E545K}$, and PI3Kα$^{WT}$. **b**, Predicted cryptic allosteric pockets for PI3Kα$^{E545K}$ and PI3Kα$^{E453K}$. The blue mesh represents the pocket with the highest probability, and the red mesh represents the next highest probability pocket. Other possible pocket locations are shown in different colors on a mesh background.

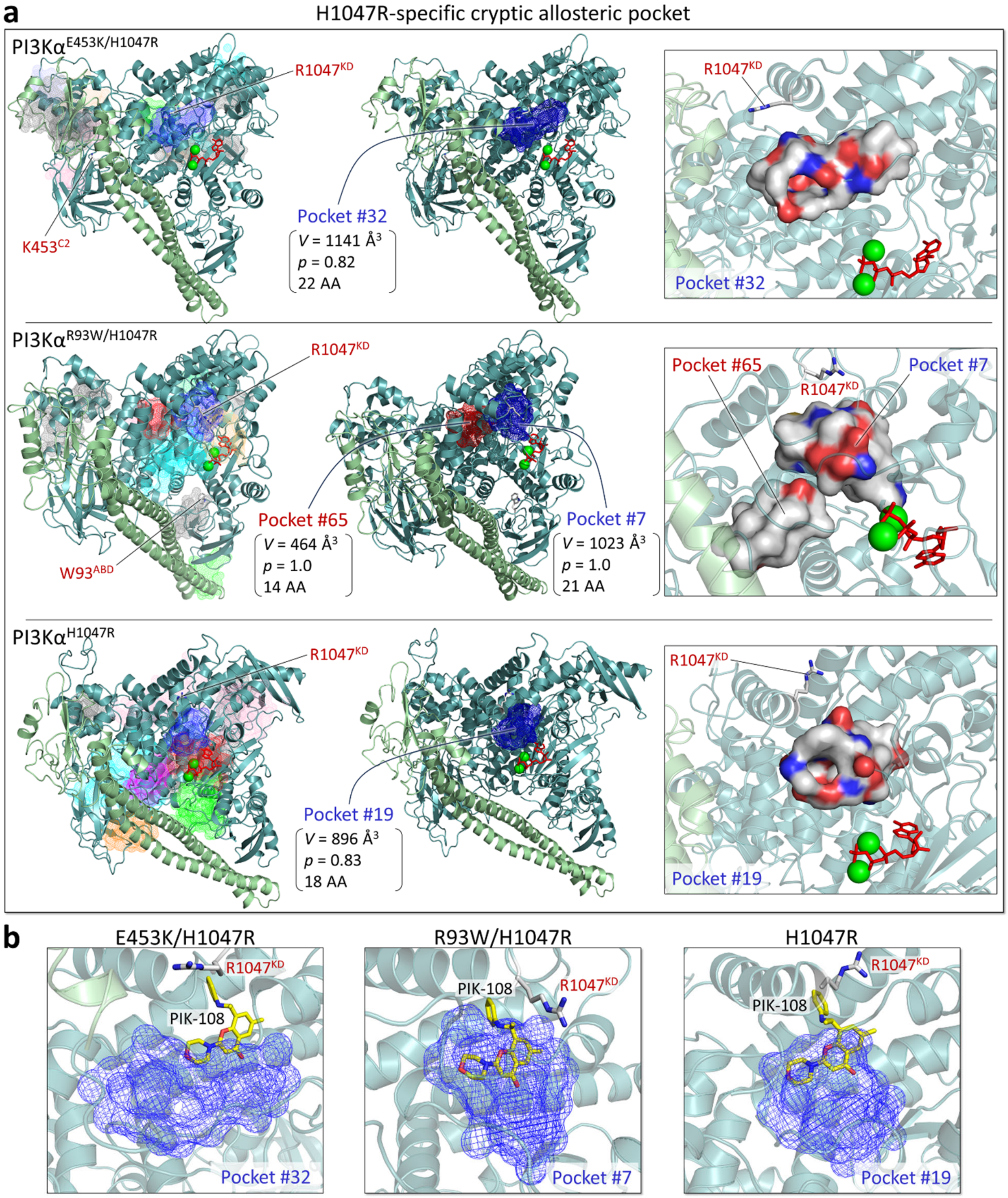


**Supplementary Fig. 16 | H1047R-specific cryptic allosteric pockets.**
**a**, Predicted cryptic allosteric pockets for PI3Kα$^{E453K/H1047R}$, PI3Kα$^{R93W/H1047R}$, and PI3Kα$^{H1047R}$ (*left*). The blue mesh represents the pocket with the highest probability, and the red mesh represents the next highest probability pocket. The pocket volume, probability, and number of amino acids involved in the pocket are indicated (*middle*). Highlights of these pockets are shown in the surface

representation (*right*). In the pocket structure, residues are colored white, green, blue, and red to indicate their hydrophobic, polar, positively charged, and negatively charged properties, respectively. **b**, Overlays of the allosteric drug PIK-108 in the pockets predicted for the E453K/H1047R (*left*), R93W/H1047R (*middle*), and H1047R (*right*) mutations.

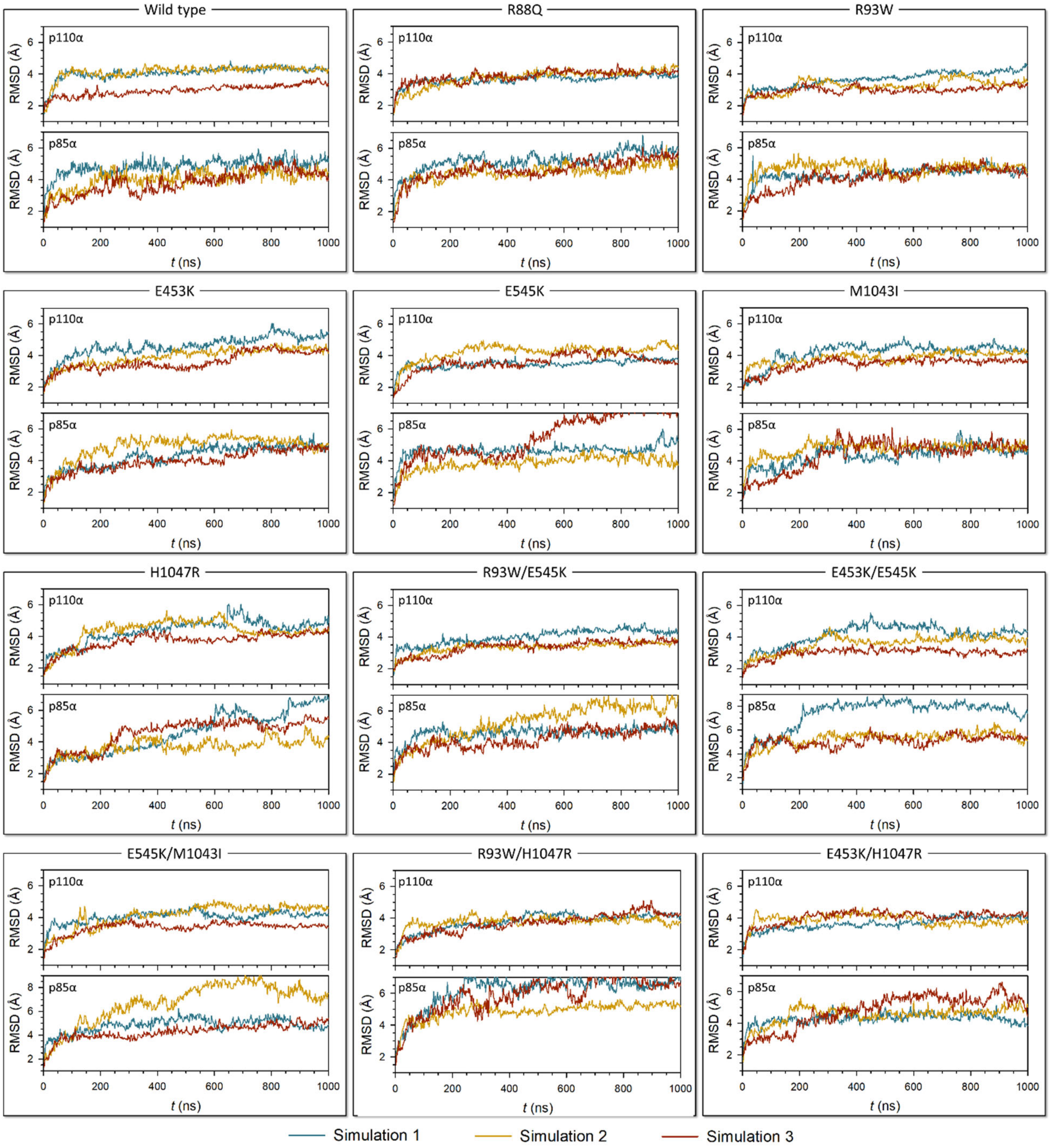


**Supplementary Fig. 17 | RMSD of p110α and p85α subunits.**
Time series showing the root-mean-squared-deviations (RMSDs) of the p110α and p85α subunits for three replica simulations of each system in solution.

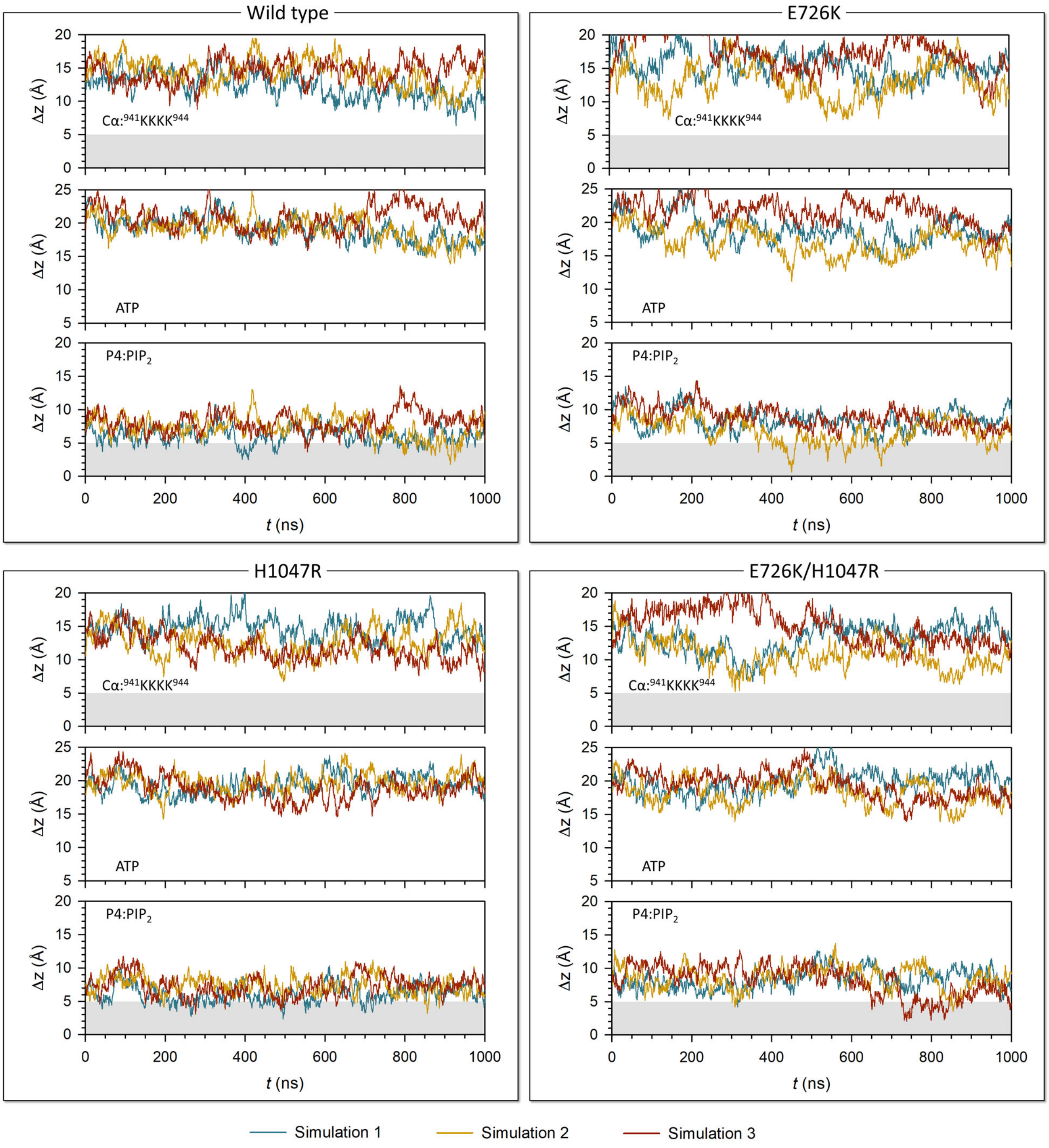


**Supplementary Fig. 18 | Location of the $_{941}$KKKK$_{944}$ motif in the A-loop, ATP, and $PIP_2$ at the membrane.**

Time series showing the deviations of the Cα atoms in the $_{941}$KKKK$_{944}$ motif in the A-loop, the center of mass of ATP, and the phosphate at the 4th position of the inositol ring of $PIP_2$ from the bilayer surface for three replica simulations of each system.

**Supplementary Table 1.** *PIK3CA* mutations and their functions, and diseases associated with them. Oncogenic hotspot mutations are in red.

| p110α mutation Interpretation (germline) [somatic] | CDS mutation Genomic location (GRCh38) [GRCh37] | Function | Disease | Reference (PMID) |
|---|---|---|---|---|
| **R88Q**<br><br>(Pathogenic)<br>[Oncogenic] | c.263G>A<br><br>(3: 179199088)<br>[3: 178916876] | Gain of function mutation, increased AKT phosphorylation, downstream target rpS6, cell proliferation, migration, and transformation ability | Ovarian neoplasm, Cowden syndrome, megalencephaly-capillary malformation-polymicrogyria syndrome, PROS, abnormal cerebral morphology, CLOVES syndrome, neoplasm | 18829572, 22729224 22949682, 28941273, 29296277, 29533785, 32778138, 34779417, 39825153 |
| **R93W**<br><br>(Pathogenic/Likely pathogenic)<br>[Likely oncogenic] | c.277C>T<br><br>(3: 179199102)<br>[3: 178916890] | Gain of function mutation, increased AKT phosphorylation and transformation ability | Megalencephaly-capillary malformation-polymicrogyria syndrome, PROS, neoplasm | 21266528, 27191687, 27631024, 28191889, 29533785, 32733937, 34733958 |
| **E453K**<br><br>(Pathogenic)<br>[No somatic data] | c.1357G>A<br><br>(3: 179210291)<br>[3: 178928079] | Gain of function mutation, increased AKT and MEK1/2 phosphorylation, cell proliferation, migration, and transformation ability, and enhanced cell survival | Ovarian neoplasm, Cowden syndrome, megalencephaly-capillary malformation-polymicrogyria syndrome, CLOVES syndrome, abnormal cardiovascular system morphology, PROS | 22729224, 26627007, 27631024, 28151489, 29533785, 30063105, 34779417, 36474027 |
| **E545K**<br><br>(Pathogenic/Likely pathogenic)<br>[Oncogenic] | c.1633G>A<br><br>(3: 179218303)<br>[3: 178936091] | Gain of function mutation, increased AKT and MEK1/2 phosphorylation, growth factor-independent cell survival, and transformation ability | Epithelial ovarian cancer, breast adenocarcinoma, carcinoma of colon, seborrheic keratosis, ovarian neoplasm, non-small cell lung carcinoma, megalencephaly-capillary malformation-polymicrogyria syndrome, sarcoma, segmental undergrowth, gallbladder cancer, neoplasm | 23946963,25880439, 26627007, 27631024, 28151489, 28425981, 29533785, 29661094, 30376034, 30543347, 31536475, 37712948 |
| **E726K**<br><br>(Pathogenic)<br>[No somatic data] | c.2176G>A<br><br>(3: 179221146)<br>[3: 178938934] | Gain of function mutation, increased cell proliferation, migration, AKT phosphorylation, and transformation ability | Inborn genetic diseases, overgrowth syndrome and/or cerebral malformations, Cowden syndrome, megalencephaly-capillary malformation-polymicrogyria syndrome, PROS | 22729224, 23246288, 24497998, 26351730, 27631024, 28151489, 28566443, 28941273, 29533785, 29549527, 34779417 |
| **M1043I**<br><br>(Pathogenic)<br>[No somatic data] | c.3129G>T<br><br>(3: 179234286)<br>[3: 178952074] | Gain of function mutation, increased AKT phosphorylation, activation of downstream signaling, and transformation | Cowden syndrome, PROS, | 15930273, 17376864, 22120714, 22729224, 29533785 |
| **H1047R**<br><br>(Pathogenic)<br>[Oncogenic] | c.3140A>G<br><br>(3: 179234297)<br>[3: 178952085] | Gain of function mutation, increased AKT and MEK1/2 phosphorylation, growth factor-independent cell survival, and transformation ability | Epithelial ovarian cancer, breast adenocarcinoma, carcinoma of colon, hepatocellular carcinoma, non-small cell lung carcinoma, seborrheic keratosis, CLOVES syndrome, macrodactyly, ovarian neoplasm, neoplasm | 23946963, 26266985, 26627007, 27126994, 29533785, 29988677, 31371346, 31536475, 34008892, 34496175, 39825153 |

*Data were imported from ClinVar (https://www.ncbi.nlm.nih.gov/clinvar) and the Jackson Laboratory Clinical Knowledgebase (CKB, https://ckb.jax.org/gene/grid).

**Supplementary Table 2.** Properties of the predicted cryptic pockets for PI3Kα$^{E453K/E545K}$. Here, $P_{Hydro}$, $P_{Polar}$, $P_{Aroma}$, and $P_{Alipha}$ correspond to the proportions of hydrophobic, polar, aromatic, and aliphatic residues in the pockets, respectively. GRAVY denotes the grand average of hydropathy (doi: 10.1016/0022-2836(82)90515-0). This is related to Fig. 8a and Supplementary Fig. 15a.

| Pocket | $P_{Hydro}$ | $P_{Polar}$ | $P_{Aroma}$ | $P_{Alipha}$ | Volume (Å$^3$) | Surface (Å$^2$) | Diameter (Å) | GRAVY | $N_{Res}$ | *p* |
|---|---|---|---|---|---|---|---|---|---|---|
| **#2** | **0.59** | **0.62** | **0.18** | **0.18** | **11603** | **2936** | **54** | **-0.90** | **71** | **0.75** |
| **#6** | **0.55** | **0.52** | **0.15** | **0.23** | **3710** | **1157** | **24** | **-0.58** | **40** | **0.63** |
| #38 | 0.88 | 0.25 | 0.31 | 0.44 | 632 | 418 | 18 | 1.83 | 16 | 0.99 |
| #50 | 0.80 | 0.13 | 0.13 | 0.60 | 575 | 364 | 14 | 2.19 | 15 | 0.99 |
| #35 | 0.71 | 0.43 | 0.29 | 0.33 | 1082 | 574 | 20 | 0.74 | 21 | 0.98 |
| #4 | 0.71 | 0.38 | 0.15 | 0.35 | 2645 | 1070 | 25 | 0.91 | 34 | 0.98 |
| #13 | 0.63 | 0.51 | 0.14 | 0.29 | 2221 | 967 | 29 | 0.15 | 35 | 0.90 |
| #21 | 0.71 | 0.52 | 0.23 | 0.23 | 2806 | 1091 | 26 | -0.18 | 31 | 0.86 |
| #53 | 0.73 | 0.64 | 0.36 | 0.14 | 1839 | 689 | 21 | -0.64 | 22 | 0.82 |
| #1 | 0.65 | 0.54 | 0.11 | 0.28 | 5900 | 1788 | 32 | -0.27 | 57 | 0.80 |
| #23 | 0.65 | 0.41 | 0.06 | 0.24 | 709 | 425 | 15 | 0.18 | 17 | 0.78 |
| #7 | 0.64 | 0.61 | 0.21 | 0.14 | 2540 | 999 | 24 | -0.82 | 28 | 0.53 |

**Supplementary Table 3.** Properties of the predicted cryptic pockets for PI3Kα$^{R93W/E545K}$. See Supplementary Table 2 for descriptions of the parameters used in the table. This is related to Fig. 8b and Supplementary Fig. 15a.

| Pocket | $P_{Hydro}$ | $P_{Polar}$ | $P_{Aroma}$ | $P_{Alipha}$ | Volume (Å$^3$) | Surface (Å$^2$) | Diameter (Å) | GRAVY | $N_{Res}$ | *p* |
|---|---|---|---|---|---|---|---|---|---|---|
| **#2** | **0.61** | **0.58** | **0.18** | **0.17** | **9043** | **2359** | **39** | **-0.92** | **71** | **0.65** |
| **#48** | **0.58** | **0.58** | **0.21** | **0.21** | **1638** | **811** | **25** | **-0.35** | **19** | **0.75** |
| #12 | 0.69 | 0.38 | 0.19 | 0.44 | 805 | 463 | 17 | 1.58 | 16 | 0.99 |
| #39 | 0.81 | 0.25 | 0.31 | 0.50 | 713 | 441 | 18 | 1.85 | 16 | 0.99 |
| #65 | 0.88 | 0.13 | 0.19 | 0.56 | 623 | 394 | 14 | 2.5 | 16 | 0.99 |
| #46 | 0.80 | 0.40 | 0.27 | 0.27 | 607 | 389 | 16 | 0.63 | 15 | 0.98 |
| #42 | 0.79 | 0.50 | 0.36 | 0.21 | 751 | 479 | 18 | -0.21 | 14 | 0.88 |
| #53 | 0.50 | 0.50 | 0 | 0.36 | 459 | 339 | 16 | 0.59 | 14 | 0.88 |
| #7 | 0.59 | 0.47 | 0.24 | 0.24 | 855 | 477 | 17 | 0.08 | 17 | 0.87 |
| #18 | 0.76 | 0.53 | 0.24 | 0.24 | 770 | 457 | 16 | 0.04 | 17 | 0.86 |
| #29 | 0.60 | 0.53 | 0.10 | 0.23 | 2432 | 974 | 25 | -0.14 | 30 | 0.78 |
| #49 | 0.57 | 0.50 | 0.29 | 0.21 | 666 | 412 | 17 | -0.43 | 14 | 0.77 |
| #1 | 0.59 | 0.60 | 0.12 | 0.21 | 13630 | 3078 | 42 | -0.68 | 91 | 0.74 |
| #9 | 0.57 | 0.57 | 0.07 | 0.29 | 510 | 335 | 14 | -0.36 | 14 | 0.50 |

**Supplementary Table 4.** Properties of the predicted cryptic pockets for PI3Kα$^{E545K}$. See Supplementary Table 2 for descriptions of the parameters used in the table. This is related to Supplementary Fig. 15b.

| Pocket | $P_{Hydro}$ | $P_{Polar}$ | $P_{Aroma}$ | $P_{Alipha}$ | Volume (Å$^3$) | Surface (Å$^2$) | Diameter (Å) | GRAVY | $N_{Res}$ | *p* |
|---|---|---|---|---|---|---|---|---|---|---|
| **#4** | **0.63** | **0.60** | **0.23** | **0.15** | **3862** | **1297** | **25** | **-0.88** | **40** | **0.60** |
| #62 | 0.86 | 0.14 | 0.21 | 0.50 | 435 | 312 | 15 | 2.39 | 14 | 0.99 |
| #64 | 0.78 | 0.39 | 0.11 | 0.39 | 1043 | 591 | 22 | 0.98 | 18 | 0.98 |
| #9 | 0.67 | 0.48 | 0.14 | 0.38 | 1184 | 604 | 20 | 0.73 | 21 | 0.97 |
| #12 | 0.68 | 0.39 | 0.18 | 0.36 | 1941 | 855 | 24 | 0.50 | 28 | 0.95 |
| #27 | 0.75 | 0.56 | 0.38 | 0.13 | 658 | 419 | 18 | -0.23 | 16 | 0.93 |
| #41 | 0.67 | 0.47 | 0.27 | 0.33 | 688 | 421 | 15 | -0.01 | 15 | 0.90 |
| #20 | 0.61 | 0.56 | 0.17 | 0.22 | 688 | 422 | 16 | -0.16 | 18 | 0.80 |
| #51 | 0.71 | 0.57 | 0.29 | 0.21 | 642 | 428 | 19 | -0.39 | 14 | 0.80 |
| #22 | 0.63 | 0.63 | 0.25 | 0.25 | 805 | 466 | 16 | -0.23 | 16 | 0.78 |
| #32 | 0.71 | 0.47 | 0.06 | 0.35 | 918 | 521 | 19 | 0.11 | 17 | 0.78 |
| #8 | 0.58 | 0.58 | 0.17 | 0.25 | 1935 | 824 | 21 | -0.29 | 24 | 0.72 |
| #6 | 0.62 | 0.62 | 0.17 | 0.24 | 5280 | 1709 | 32 | -0.64 | 42 | 0.67 |
| #16 | 0.53 | 0.59 | 0.12 | 0.18 | 741 | 428 | 15 | -0.37 | 17 | 0.56 |

**Supplementary Table 5.** Properties of the predicted cryptic pockets for PI3Kα$^{E453K}$. See Supplementary Table 2 for descriptions of the parameters used in the table. This is related to Supplementary Fig. 15b.

| Pocket | $P_{Hydro}$ | $P_{Polar}$ | $P_{Aroma}$ | $P_{Alipha}$ | Volume (Å$^3$) | Surface (Å$^2$) | Diameter (Å) | GRAVY | $N_{Res}$ | *p* |
|---|---|---|---|---|---|---|---|---|---|---|
| **#10** | **0.69** | **0.56** | **0.19** | **0.19** | **876** | **494** | **17** | **-0.31** | **16** | **0.69** |
| #45 | 0.86 | 0.29 | 0.21 | 0.36 | 480 | 332 | 14 | 2.16 | 14 | 0.99 |
| #52 | 0.80 | 0.27 | 0.20 | 0.53 | 680 | 417 | 15 | 2.01 | 15 | 0.99 |
| #27 | 0.93 | 0.40 | 0.33 | 0.27 | 574 | 379 | 15 | 0.71 | 15 | 0.98 |
| #36 | 0.80 | 0.33 | 0 | 0.40 | 784 | 466 | 16 | 0.99 | 15 | 0.95 |
| #24 | 0.64 | 0.50 | 0.21 | 0.36 | 365 | 271 | 11 | 0.34 | 14 | 0.91 |
| #63 | 0.58 | 0.47 | 0.21 | 0.26 | 1123 | 595 | 18 | -0.19 | 19 | 0.84 |
| #3 | 0.63 | 0.57 | 0.14 | 0.29 | 5653 | 1810 | 37 | -0.32 | 49 | 0.82 |
| #7 | 0.64 | 0.64 | 0.29 | 0.21 | 879 | 516 | 18 | -0.30 | 14 | 0.78 |
| #33 | 0.64 | 0.57 | 0.21 | 0.21 | 544 | 359 | 14 | -0.35 | 14 | 0.74 |
| #6 | 0.61 | 0.67 | 0.17 | 0.17 | 920 | 523 | 17 | -0.36 | 18 | 0.70 |
| #54 | 0.57 | 0.50 | 0.07 | 0.36 | 598 | 384 | 15 | -0.14 | 14 | 0.68 |
| #12 | 0.61 | 0.56 | 0.28 | 0.17 | 910 | 504 | 17 | -0.52 | 18 | 0.67 |
| #41 | 0.53 | 0.47 | 0.12 | 0.29 | 1042 | 569 | 19 | -0.32 | 17 | 0.63 |

**Supplementary Table 6.** Properties of the predicted cryptic pockets for PI3Kα$^{WT}$. See Supplementary Table 2 for descriptions of the parameters used in the table. This is related to Fig. 8c and Supplementary Fig. 15a.

| Pocket | $P_{Hydro}$ | $P_{Polar}$ | $P_{Aroma}$ | $P_{Alipha}$ | Volume (Å$^3$) | Surface (Å$^2$) | Diameter (Å) | GRAVY | $N_{Res}$ | *p* |
|---|---|---|---|---|---|---|---|---|---|---|
| **#9** | **0.71** | **0.60** | **0.20** | **0.17** | **3475** | **1266** | **29** | **-0.81** | **35** | **0.58** |
| #42 | 0.87 | 0.20 | 0.20 | 0.67 | 576 | 366 | 13 | 2.5 | 15 | 0.99 |
| #17 | 0.60 | 0.43 | 0.20 | 0.33 | 2040 | 891 | 23 | 0.67 | 30 | 0.97 |
| #21 | 0.72 | 0.48 | 0.68 | 0.24 | 1860 | 846 | 25 | 0.68 | 25 | 0.96 |
| #16 | 0.68 | 0.58 | 0.26 | 0.26 | 2798 | 1097 | 25 | -0.09 | 31 | 0.91 |
| #11 | 0.72 | 0.50 | 0.19 | 0.25 | 3824 | 1337 | 29 | -0.08 | 36 | 0.91 |
| #2 | 0.69 | 0.51 | 0.15 | 0.26 | 6509 | 1912 | 36 | -0.10 | 61 | 0.90 |
| #14 | 0.70 | 0.48 | 0.15 | 0.27 | 2391 | 978 | 25 | 0.08 | 33 | 0.87 |
| #7 | 0.69 | 0.64 | 0.23 | 0.15 | 4050 | 1402 | 32 | -0.38 | 39 | 0.86 |
| #35 | 0.64 | 0.50 | 0.14 | 0.36 | 612 | 398 | 15 | -0.24 | 14 | 0.72 |
| #22 | 0.63 | 0.42 | 0.11 | 0.11 | 844 | 486 | 17 | -0.08 | 19 | 0.72 |
| #25 | 0.53 | 0.40 | 0.07 | 0.40 | 595 | 378 | 14 | 0.01 | 15 | 0.71 |
| #5 | 0.55 | 0.68 | 0.18 | 0.20 | 3919 | 1328 | 27 | -0.96 | 40 | 0.51 |

**Supplementary Table 7.** Properties of the predicted cryptic pockets for PI3Kα$^{E453K/H1047R}$. See Supplementary Table 2 for descriptions of the parameters used in the table. This is related to Supplementary Fig. 16a.

| Pocket | $P_{Hydro}$ | $P_{Polar}$ | $P_{Aroma}$ | $P_{Alipha}$ | Volume (Å$^3$) | Surface (Å$^2$) | Diameter (Å) | GRAVY | $N_{Res}$ | *p* |
|---|---|---|---|---|---|---|---|---|---|---|
| **#32** | **0.73** | **0.50** | **0.23** | **0.27** | **1141** | **618** | **23** | **-0.32** | **22** | **0.82** |
| #19 | 0.71 | 0.29 | 0.12 | 0.53 | 681 | 414 | 15 | 2.03 | 17 | 0.99 |
| #79 | 0.99 | 0.07 | 0.29 | 0.57 | 629 | 396 | 15 | 3.19 | 14 | 0.99 |
| #13 | 0.75 | 0.44 | 0.31 | 0.31 | 565 | 362 | 14 | 0.90 | 16 | 0.98 |
| #49 | 0.71 | 0.43 | 0.07 | 0.43 | 548 | 372 | 18 | 1.13 | 14 | 0.98 |
| #12 | 0.73 | 0.57 | 0.30 | 0.33 | 2763 | 1108 | 28 | 0.04 | 30 | 0.95 |
| #3 | 0.65 | 0.41 | 0.24 | 0.29 | 980 | 522 | 16 | 0.52 | 17 | 0.95 |
| #8 | 0.76 | 0.47 | 0.18 | 0.35 | 609 | 381 | 14 | 0.52 | 17 | 0.94 |
| #7 | 0.71 | 0.43 | 0.21 | 0.29 | 579 | 386 | 14 | 0 | 14 | 0.83 |
| #43 | 0.59 | 0.59 | 0.18 | 0.18 | 1502 | 723 | 22 | -0.40 | 22 | 0.74 |
| #76 | 0.59 | 0.53 | 0.18 | 0.24 | 853 | 480 | 18 | -0.22 | 17 | 0.73 |
| #26 | 0.53 | 0.53 | 0.12 | 0.29 | 669 | 422 | 16 | -0.37 | 17 | 0.57 |
| #18 | 0.64 | 0.50 | 0 | 0.29 | 901 | 507 | 18 | -0.22 | 14 | 0.53 |

**Supplementary Table 8.** Properties of the predicted cryptic pockets for PI3Kα$^{R93W/H1047R}$. See Supplementary Table 2 for descriptions of the parameters used in the table. This is related to Supplementary Fig. 16a.

| Pocket | $P_{Hydro}$ | $P_{Polar}$ | $P_{Aroma}$ | $P_{Alipha}$ | Volume (Å$^3$) | Surface (Å$^2$) | Diameter (Å) | GRAVY | $N_{Res}$ | *p* |
|---|---|---|---|---|---|---|---|---|---|---|
| **#7** | **0.90** | **0.38** | **0.33** | **0.29** | **1023** | **551** | **18** | **1.53** | **21** | **0.99** |
| **#65** | **0.99** | **0** | **0.21** | **0.57** | **464** | **339** | **14** | **3.46** | **14** | **0.99** |
| #35 | 0.87 | 0.40 | 0.07 | 0.47 | 974 | 547 | 18 | 1.03 | 15 | 0.97 |
| #4 | 0.65 | 0.45 | 0.25 | 0.30 | 1311 | 640 | 20 | 0.43 | 20 | 0.95 |
| #42 | 0.73 | 0.53 | 0.13 | 0.40 | 656 | 420 | 16 | 0.65 | 15 | 0.94 |
| #32 | 0.67 | 0.47 | 0.13 | 0.27 | 587 | 385 | 16 | 0.39 | 15 | 0.92 |
| #15 | 0.65 | 0.59 | 0.18 | 0.18 | 622 | 407 | 17 | -0.06 | 17 | 0.79 |
| #31 | 0.71 | 0.53 | 0.24 | 0.24 | 1168 | 603 | 20 | -0.34 | 17 | 0.78 |
| #16 | 0.58 | 0.58 | 0.16 | 0.26 | 1098 | 579 | 18 | -0.19 | 19 | 0.74 |
| #53 | 0.60 | 0.60 | 0.20 | 0.33 | 801 | 497 | 18 | -0.58 | 15 | 0.62 |

**Supplementary Table 9.** Properties of the predicted cryptic pockets for PI3Kα$^{H1047R}$. See Supplementary Table 2 for descriptions of the parameters used in the table. This is related to Supplementary Fig. 16a.

| Pocket | $P_{Hydro}$ | $P_{Polar}$ | $P_{Aroma}$ | $P_{Alipha}$ | Volume (Å$^3$) | Surface (Å$^2$) | Diameter (Å) | GRAVY | $N_{Res}$ | *p* |
|---|---|---|---|---|---|---|---|---|---|---|
| **#19** | **0.72** | **0.56** | **0.33** | **0.22** | **896** | **503** | **18** | **-0.24** | **18** | **0.83** |
| #53 | 0.80 | 0.40 | 0.27 | 0.33 | 758 | 470 | 19 | 0.57 | 15 | 0.97 |
| #6 | 0.74 | 0.48 | 0.39 | 0.21 | 1347 | 650 | 20 | 0.07 | 23 | 0.97 |
| #7 | 0.79 | 0.57 | 0.21 | 0.36 | 591 | 389 | 16 | 0.72 | 14 | 0.97 |
| #10 | 0.68 | 0.50 | 0.23 | 0.27 | 1015 | 548 | 18 | 0.22 | 22 | 0.94 |
| #11 | 0.69 | 0.46 | 0.23 | 0.31 | 1697 | 819 | 26 | 0.22 | 26 | 0.94 |
| #2 | 0.59 | 0.49 | 0.13 | 0.28 | 7442 | 2116 | 34 | -0.16 | 61 | 0.87 |
| #41 | 0.69 | 0.50 | 0.13 | 0.13 | 607 | 393 | 17 | 0.09 | 16 | 0.81 |
| #25 | 0.47 | 0.59 | 0.18 | 0.18 | 763 | 457 | 16 | -0.38 | 17 | 0.63 |
| #3 | 0.58 | 0.54 | 0.10 | 0.17 | 5948 | 1780 | 34 | -0.60 | 48 | 0.63 |
| #18 | 0.51 | 0.54 | 0.09 | 0.23 | 3869 | 1349 | 31 | -0.53 | 35 | 0.59 |
| #38 | 0.57 | 0.57 | 0.14 | 0.21 | 1304 | 662 | 20 | -0.47 | 14 | 0.58 |

**Supplementary Table 10.** Details of simulation systems. Oncogenic hotspot mutations are in red.

<table>
<tr><th>System</th><th>Mutation site in p110α</th><th>Mutation type</th><th>Simulation</th></tr>
<tr><td>Wild type</td><td>—</td><td>—</td><td rowspan="12">• Started with an inactive PI3Kα conformation in solution<br>• Performed 3 replica simulations, each with 1 μs simulation<br>• TIP3P water in a cubic box</td></tr>
<tr><td>R88Q</td><td>ABD</td><td>Moderate</td></tr>
<tr><td>R93W</td><td>ABD</td><td>Weak</td></tr>
<tr><td>E453K</td><td>C2</td><td>Weak</td></tr>
<tr><td>E545K</td><td>Helical</td><td>Hotspot</td></tr>
<tr><td>M1043I</td><td>KD (C-term)</td><td>Weak</td></tr>
<tr><td>H1047R</td><td>KD (C-term)</td><td>Hotspot</td></tr>
<tr><td>R93W/E545K</td><td>ABD/Helical</td><td>Weak/Hotspot</td></tr>
<tr><td>E453K/E545K</td><td>C2/Helical</td><td>Weak/Hotspot</td></tr>
<tr><td>M1043I/E545K</td><td>KD/Helical</td><td>Weak/Hotspot</td></tr>
<tr><td>R93W/H1047R</td><td>ABD/KD</td><td>Weak/Hotspot</td></tr>
<tr><td>E453K/H1047R</td><td>C2/KD</td><td>Weak/Hotspot</td></tr>
<tr><td>Wild type</td><td>—</td><td>—</td><td rowspan="4">• Started with an active PI3Kα conformation with truncated nSH2 in the membrane<br>• Performed 3 replica simulations, each with 1 μs simulation<br>• Anionic lipid bilayer composed of DOPC:DOPS:$PIP_2$ (28:6:1 molar ratio)</td></tr>
<tr><td>E726K</td><td>KD (N-term)</td><td>Weak</td></tr>
<tr><td>H1047R</td><td>KD (C-term)</td><td>Hotspot</td></tr>
<tr><td>E726K/H1047R</td><td>KD/KD</td><td>Weak/Hotspot</td></tr>
</table>
